# Gene dosage screens in yeast reveal core signalling pathways controlling heat adaptation

**DOI:** 10.1101/2020.08.26.267674

**Authors:** Cosimo Jann, Andreas Johansson, Justin D. Smith, Leopold Parts, Lars M. Steinmetz

**Affiliations:** European Molecular Biology Laboratory (EMBL), Genome Biology Unit, Heidelberg, Germany; ETH Zurich, Department of Biology, Institute of Biochemistry, Zurich, Switzerland; Department of Genetics, Stanford University School of Medicine, Stanford, California, USA; Stanford Genome Technology Center, Stanford University, Palo Alto, California, USA; Wellcome Sanger Institute, Hinxton, UK; Department of Computer Science, University of Tartu, Tartu, Estonia

**Keywords:** CRISPRi/a, CRISPR/dCas9 screen, heat shock response, thermotolerance, Hsf1, HOG pathway, PKA signalling

## Abstract

Heat stress causes proteins to unfold and lose their function, jeopardizing essential cellular processes. To protect against heat and proteotoxic stress, cells mount a dedicated stress-protective programme, the so-called heat shock response (HSR). Our understanding of the mechanisms that regulate the HSR and their contributions to heat resistance and growth is incomplete. Here we employ CRISPRi/a to down- or upregulate protein kinases and transcription factors in *S. cerevisiae*. We measure gene functions by quantifying perturbation effects on HSR activity, thermotolerance, and cellular fitness at 23, 30 and 38°C. The integration of these phenotypes allowed us to identify core signalling pathways of heat adaptation and reveal novel functions for the high osmolarity glycerol, unfolded protein response and protein kinase A pathways in adjusting both thermotolerance and chaperone expression. We further provide evidence for unknown cross-talk of the HSR with the cell cycle-dependent kinase Cdc28, the primary regulator of cell cycle progression. Finally, we show that CRISPRi efficiency is temperature-dependent and that different phenotypes vary in their sensitivity to knock-down. In summary, our study quantifies regulatory gene functions in different aspects of heat adaptation and advances our understanding of how eukaryotic cells counteract proteotoxic and other heat-caused damage.

## Introduction

When exposed to high temperature, cells need to resist the proteotoxicity due to protein misfolding and aggregation^1,2^and other heat-caused damage^3^. They do so by eliciting a series of stress-protective events, referred to as the heat shock response (HSR)^4^. The HSR is highly conserved across eukaryotes and characterized by the fierce production of heat shock proteins (HSPs) which mostly function in maintaining protein homeostasis^5,6^. Dysregulation of the HSR results in altered chaperone capacity and is linked to neurodegenerative diseases^7,8^ and aging^9^ where decreased HSR activity aggravates proteotoxicity. Cancers also hi-jack and increase HSR activity to cope with their proteotoxic burden^10^. In the last two decades, heat-induced changes to the transcriptome and proteome have been well characterized^11–17^. In budding yeast, a shift from 30 to 37°C causes around a thousand genes to change transcription^11,17^, while exposure to 42°C results in expression changes for more than 50% of the yeast genome (∼3100 genes), with higher magnitude and longer upkeep compared to 37°C^17^. Much less is known about the mechanisms that enable regulation of this response which is essential to safeguard cellular survival.

Heat shock factor 1 (Hsf1) is considered the master HSR regulator, in yeast acting together with the general stress response factors Msn2 and Msn4^18,19^. Hsf1 is regulated through titration by chaperones^20–23^ and hyperphosphorylation^24,25^. Dissection of the Hsf1-driven HSR in yeast^26^ and human cells^27^ revealed new mechanisms controlling Hsf1. However, recent studies demonstrate that the HSR remains largely unchanged when Hsf1 is absent in yeast and mammalian cells, and mainly driven by other transcriptional regulators^15,28^. In addition, even for Hsf1, Msn2 and Msn4, the most prominent transcription factors (TFs) of the yeast HSR, the protein kinases (PKs) mediating their heat-induced hyperphosphorylation and activation remain elusive^29^. Individual TFs are likely controlled by an interplay of signalling pathways that, apart from transcription^15^, may also affect mRNA localization^30^, stability^31^ and translation^32^.

HSR overlaps with oxidative and general stress responses^11,19^ which trigger cell cycle arrest and the cell wall integrity (CWI) pathway^33,34^. The HSR also inhibits target of rapamycin (TOR) signalling^35^ and is itself repressed by protein kinase A (PKA)^36,37^. High temperature further activates the high osmolarity glycerol (HOG) pathway, although its role is unknown^38–40^. A comprehensive understanding of how signalling programmes integrate to regulate the HSR is missing. In addition, it is unclear which molecular branches of the HSR contribute to cellular protection, given that the bulk of heat-induced genes^17,28^ is dispensable for tolerance to both acute and anticipated stress^41–43^.

Here we dissect HSR regulation by CRISPR interference and activation (CRISPRi/a) systems that employ a catalytically inactive Cas9 nuclease fused to transcriptional repression or activation domains^44,45^. Only a handful of studies reported the use of these technologies for functional genomic screens in *S. cerevisiae*, mostly assaying effects on growth^46–50^. We employ inducible CRISPRi/a^46,51^ to modulate the abundance of protein kinases (PKs) and transcription factors (TFs), key regulators of almost every cellular pathway and trait, and screen for gene functions in cellular fitness at temperatures 23°C, 30°C and 38°C, HSR activity and thermotolerance. We discover a handful of genes capable of tuning thermotolerance by altering chaperone expression, including principal regulators of the HSR, the unfolded protein response (UPR), as well as the HOG and PKA pathways. We further find that CRISPRi effect size is temperature-dependent and that diverse traits are differentially sensitive to knock-down. Altogether, our study reveals the HSR as a complex programme, regulated by multiple molecular pathways and coupled with diverse cellular mechanisms to confer a rapid and precisely tuned adaptation to heat.

## Results

### CRISPRi/a efficiently modulate gene expression

We first validated the performance of the employed perturbation and reporter systems. To confirm CRISPRi effects on growth, we repressed the essential *HSF1* gene. As expected, this resulted in decreased growth rate, with varied effect size for three gRNAs differing in target sequence and distance to the transcription start site (TSS) (Fig. 1a and b). Effects were specifically observed in the presence of the gRNA-inducing compound anhydrotetracycline (ATc) (Supplementary Fig. S1).

**Figure 1.**
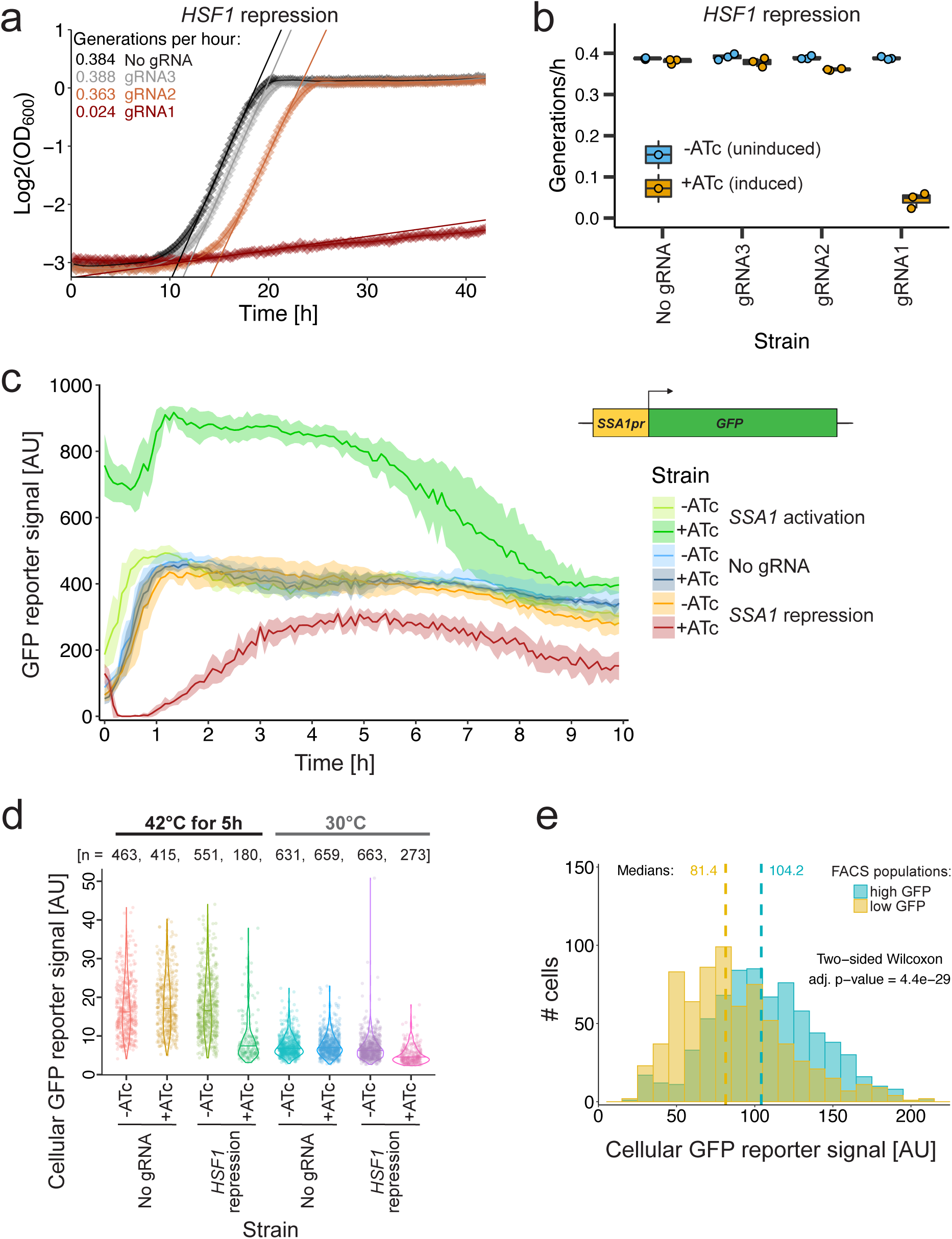
Evaluation of CRISPRi/a effects and the HSR reporter. **a**, Growth curves for repression of *HSF1* with three different gRNAs. The optical density at 600 nm (OD600) of strains grown with ATc to induce CRISPRi was measured over time. Lines denote linear fits. **b**, Generation times of different *HSF1* CRISPRi strains (x-axis), measured in n=3 replicate well cultures. **c**, Normalized GFP signal of *SSA1* CRISPRa (act) and CRISPRi (rep) strains over time. Cultures were grown at 30°C, calibrated at OD600=0.3 and exposed to 40°C throughout the experiment. The y-axis denotes GFP intensity normalized by OD600. Lines denote means and ribbons denote standard deviations of n=6 replicate well cultures, respectively. The chromosomally inserted HSR reporter is depicted as inlay. **d**, Cellular reporter signal of CRISPRi strains at 30°C and exposed to 42°C for 5h. Dots denote single cells imaged by fluorescence microscopy. Horizontal lines mark medians. **e**, Cellular GFP intensity of CRISPRi strains (merged data for TF and PK libraries) sorted for high or low cellular GFP intensity after heat shock, regrown for more than 20 generations, and imaged by microscopy. The cellular GFP intensity (x-axis) is shown for bins of size 10. Dashed lines denote medians.

We quantify HSR activity with a heat-responsive reporter based on the truncated promoter of the *SSA1 HSP70* gene^52,53^ driving expression of an ultra-fast maturing GFP^54^. We validated our CRISPRi/a systems by targeting this promoter, achieving efficient knock-down and overexpression of Hsp70 protein, respectively (Fig. 1c). The ATc-induced CRISPRa strain had fivefold increased reporter signal already before heat shock (t=0), as expected for strong activation (dark green curve in Fig. 1c). Interestingly, heat exposure (t>0) resulted in ∼30 min delayed *SSA1* expression compared to the non-induced CRISPRa strain. The *SSA1* promoter was thus not immediately induced if Hsp70 protein levels were already elevated, in line with its ability to inhibit Hsf1^20,25^.

As a proof of concept for using the HSR reporter for functional genomics, we tested its responsiveness to Hsf1. Repression of Hsf1 decreased Ssa1 protein levels in heat and non-stress conditions (Fig. 1d), as expected from *SSA1* mRNA changes after Hsf1 depletion^28^ and chromatin-immunoprecipitation (ChIP) of the *SSA1* promoter together with Hsf1 protein^55^. The GFP-based reporter is selectable by Fluorescence-Automated Cell Sorting (FACS) and CRISPRi effects were inherited over at least 20 generations (Fig. 1e), indicating excellent suitability for genetic screens.

### Gene dosage effects on cellular fitness

We first characterised CRISPRi effects on fitness (Fig. 2a). We repressed sets of either 129 protein kinases^56^ or 161 transcription factors^57^ with up to six gRNAs per gene (Supplementary Fig. S2). Out of 1573 gRNAs in both libraries, 271 were significantly depleted (two-fold depletion, FDR<0.05) after two days of competitive growth at 30°C (Supplementary Fig. S3). Approximately 40% of gRNAs were effective (Fig. 2b), based on the dropout of essential genes defined as non-viable deletions according to the *Saccharomyces* Genome Database (SGD)^58^. CRISPRi efficiency depends on the GC content and secondary structure of gRNAs (Supplementary Fig. S4), and the chromatin accessibility at the targeted genomic locus (Supplementary Fig. S5) which supports and complements previous findings^46,47^. Based on the distance between TSS and gRNA target locus, the optimal range is between TSS-150 to TSS+25 nucleotides, with minor variation between target strands (Supplementary Fig. S4b).

**Figure 2.**
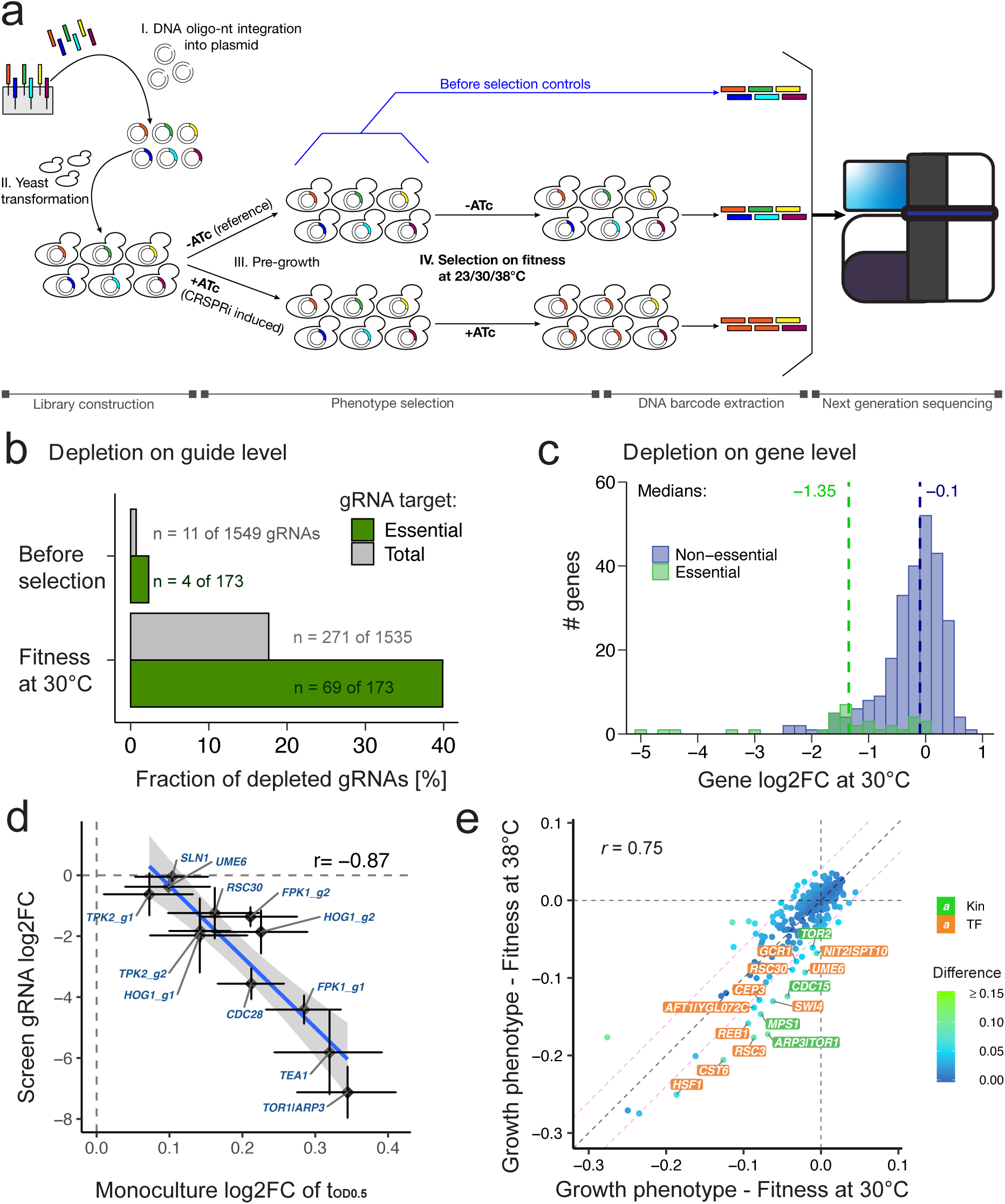
CRISPRi effects are reproducible, capture positive controls and novel fitness modulators. **a**, Experimental set-up for competitive growth screens. Briefly, a computationally designed oligonucleotide library was cloned into plasmids and transformed into yeast. CRISPRi induced (+ATc) and reference yeast cultures (-ATc) were then profiled for fitness, followed by plasmid DNA extraction and gRNA barcodes sequencing. **b**, Frequency of CRISPRi strain dropout from populations before selection (after 10 h pre-growth, see Methods) and after two days selection at 30°C. Fractions of gRNAs with log2FC<=-1 and FDR<0.05 are shown for total and essential genes (merged for PK and TF library data). Numbers denote displayed ratios. **c**, Depletion of essential (green) and non-essential genes (blue) for fitness at 30°C. Histogram bins have size 0.2 and dashed lines denote medians. Gene log2FCs represent mean gRNA log2FCs per gene. **d**, Comparison of screen gRNA log2FC with monoculture log2-scale effects on tOD0.5 (time until cultures reach half-maximal OD) which reports on growth rate and lag time. Error bars denote standard deviations of n=3 well cultures (x-axis) and n=2 replicates for the fitness at 30°C screen (y-axis). Spearman correlation is shown. Repressed genes are labelled. If multiple gRNAs were used, these are included in labels with g[1-9]. Dashed grey lines are intercepts marking a fold change of 0. The linear model fit was generated with the R ggplot2 function geom_smooth, using default parameters and method=“lm”. Hsf1 was not included since maximum OD values are not meaningful for severe growth defects (see gRNA1 in Fig. 1a). **e**, CRISPRi effects on cellular fitness at 30 versus 38°C. Dots denote generation-normalized gene log2FCs coloured by difference and Spearman correlation is shown. Genes with heat-sensitive phenotypes were labelled in green (PKs) and orange boxes (TFs). Dashed purple lines mark diagonals indicating difference thresholds at -/+ 0.04. Dashed grey lines are intercepts marking a fold change of 0 and the diagonal.

Cellular fitness decreased upon repression of 68 genes (Fig. 2c, Supplementary Tab. 1). These were enriched for essential functions (34% compared to 12% in the background) with roles in ribosome biogenesis, cell cycle and chromosome segregation (Supplementary Fig. S6). Out of the 34 targeted essential genes, 31 had at least one effective gRNA, and 23 were depleted with two or more supporting gRNAs (Supplementary Fig. S7). In general, fitness effects correlated with knock-out (KO) screens, despite differences in assay conditions and readouts (Supplementary Fig. S8), and outperformed heterozygous deletions in detecting gene essentiality (Supplementary Fig. S9). We measured novel fitness-modulatory roles for eight open reading frames (ORFs) (*CAD1, FPK1, IKS1, NHP6A, RSC30, SCH9, TEA1, TPK2*) and two ambiguous loci where multiple TSS were potentially targeted (*FUS3|PEP1, MMO1|PHD1*).

To validate screen performance, we selected ten ORFs for further characterization; six known to affect growth and four measured with new functions. We observed high correlation between screen fold changes and individually determined growth rates (Spearman *r*=0.77; Supplementary Fig. S10), and half-maximal OD intervals (tOD0.5) which additionally report on lag time (Spearman *r*=-0.87; Fig. 2d). This follow-up confirmed novel roles in fitness regulation for all four genes included (*FPK1, RSC30, TEA1, TPK2*).

### Genetic requirements for growth at high temperature

Having validated our assay at 30°C, we further screened for fitness effects at 23 and 38°C, detecting 18 and 30 depleted genes, respectively (Supplementary Tab. 1). Effect size and statistical power were reduced compared to the 30°C screen due to less generations of selection, considering that doubling time was lowered by ∼60% at 23°C and by ∼30% at 38°C (Supplementary Fig. S11-12). Generation-normalized fold changes correlated well between temperatures, demonstrating high reproducibility not only for read counts (Supplementary Fig. S13) but also for CRISPRi effects (Supplementary Fig. S14). We detected 15 genes causing heat sensitivity upon repression, derived from reduced fitness at 38 compared to 30°C (Fig. 2e), nine of which were known, such as Hsf1 and Swi4 which control HSR transcription^59,60^, Reb1 which enhances Hsf1 transactivation^61^, Ume6 which promotes Msn2/4-dependent transcription as part of the Rpd3L histone deacetylation complex^62^, and Cst6 with a yet unknown but predicted role in the HSR^63^. Quantification of mRNA and gRNA levels in Hsf1 and Ume6 CRISPRi strains over time and temperatures confirmed efficient repression in all conditions, showing that heat sensitivity is not simply due to stronger repression at higher temperature (Supplementary Fig. S15). Novel roles in heat sensitivity were detected for Cep3, Gcr1, Tor2 and Rsc30. Supporting this, Gcr1 controls the expression of glycolysis genes^64^ and Tor2 regulates cytoskeleton organization^65^, both important for thermal adaptation^3^. Rsc30 is part of the RSC complex which translocates from ORFs to promoters in heat to facilitate nucleosome dissociation^66^ and Hsf1-mediated transcription^61^, akin to the known hit Rsc3 (Fig. 2e).

### Modulators of the heat shock response

Most transcripts induced during heat shock appear to not serve protective functions and it is therefore debated if they compensate for loss-of-function effects due to protein instability^41^. Combining published RNAseq time course with thermal proteome profiling data, we found that heat-induced transcripts encode proteins with higher than average thermal stability, while proteins with low stability are down-regulated (Supplementary Fig. S16). This suggests that the HSR is a purposeful programme to enhance heat resistance rather than overproducing proteins that go astray.

To identify components controlling the HSR pathway, we established a flow cytometry assay (Fig. 3a) based on the Hsp70 reporter (introduced in Fig. 1c). This allowed us to measure impacts on chaperone expression independent of fitness effects (Fig. 3b & Supplementary Fig. S11). Out of the 290 TFs and PKs, we found twenty to decrease and seven to increase HSR activity upon repression, implying functions in promoting and inhibiting the HSR, respectively (Supplementary Tab. 1). CRISPRi effects on the Hsp70 promoter cannot be fully explained by previous screens with artificial promoters based on either heat shock elements (HSEs) recognized by Hsf1 or stress response elements bound by Msn2/4, using deletion mutants and a one hour heat shock at 37°C (Supplementary Fig. S17)^26^. Most of our hits were thus not known to tune chaperone expression during the HSR, although we found that individual roles were supported by studies that probed 35 or 68 gene deletions with an *HSP12-GFP* reporter gene^67,68^.

**Figure 3.**
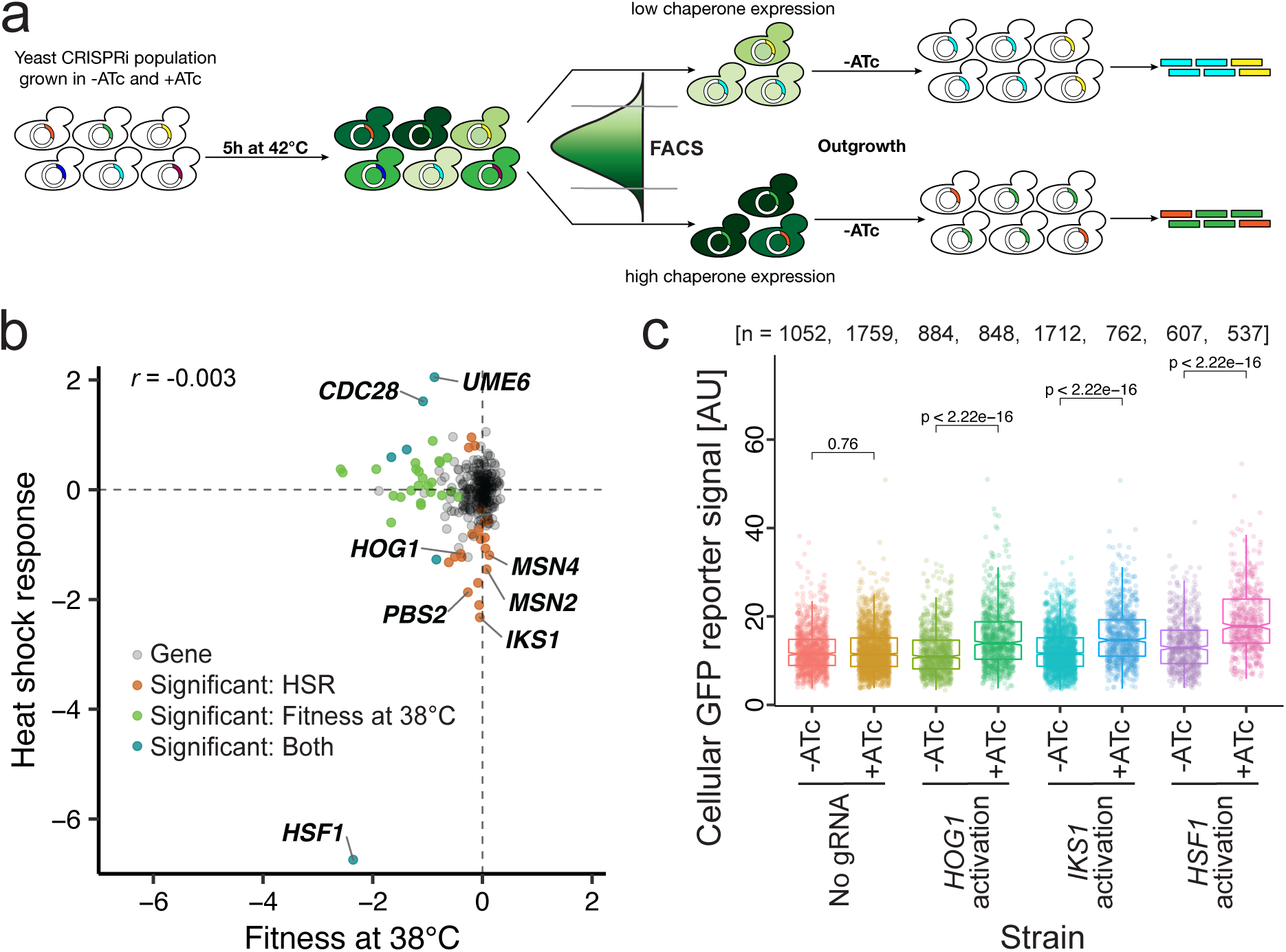
Modulators of HSR activity. **a**, Schematic of the reporter-based HSR screen. After heat exposure, cells with extremely high and low GFP intensity were collected by FACS and identified through barcode sequencing. **b**, Comparison of gene log2 fold changes for cellular fitness at 38°C versus HSR with Spearman correlation. Dots denote genes (n=290), coloured by screen effects as indicated in figure legend. Selected HSR modulators are labelled. **c**, Cellular *SSA1pr-GFP* reporter intensity (y-axis) of individual CRISPRa strains cultured with or without ATc (x-axis) after 5h exposure to 42°C, and imaged with fluorescence microscopy. Dots denote single cells. Used gRNAs are Hog1_g1, Iks1_g5, Hsf1_g1. Two-sided Wilcoxon adjusted p-values are depicted for tests between samples.

Hsf1 was the most potent HSR activator (Fig. 3b), in agreement with single cell microscopy results (Fig. 1d). Additionally, Msn2 and Msn4 were measured as strong HSR stimulators, further validating the experimental setup (Fig. 3b). The two regulators of the environmental stress response are redundant for growth and thermotolerance, as discovered by Martínez-Pastor et al. (1996)^69^ and confirmed by our study. In contrast, repression of either factor decreased Hsp70 expression which demonstrates their non-redundant roles in stimulating transcription as part of the HSR^11^. Notably, the second strongest HSR stimulator in the panel was Iks1, a putative kinase of unknown function which is transcriptionally induced at 37°C^70^ and during proteotoxic stress^71^ (Fig. 3b). We measured further genes that promote HSR activity, derived from decreased reporter signal upon repression, that encode chromatin remodellers (Rsc30, Nhp6A), activators of the plasma membrane ATPase (Ptk2, Hrk1) stress-related kinases (Mck1, Rim15, Yak1), and surprisingly also central components of the UPR (Ire1), the high osmolarity glycerol (HOG) pathways (Hog1, Pbs2, Rck2) and the protein kinase A (PKA) subunit Tpk2. Additionally, we determined found four kinases (Cdc28, Hrr25, Mps1, Sch9) and three TFs (Rim101, Sok2, Ume6) with roles in alleviating HSR activity. We confirmed that Cdc28 counteracts the HSR by FACS (Supplementary Fig. S18a) which further agrees with screen measurements for its regulators (Supplementary Fig. S18b). All three HSR-antagonizing TFs act as transcriptional repressors^72–74^, in line with their inhibiting roles.

Microscopy follow-ups confirmed screen results for the repression of Hog1, Iks1 and Ume6 (Supplementary Fig. 19). In addition, we show that CRISPRi screen phenotypes can be reversed using CRISPRa strains for Hog1, Iks1 and Hsf1 by CRISPRa (Fig. 3e). HSR-stimulating kinases, such as Iks1, Hog1, Rim15 and Yak1 potentially activate a potent TF. Rim15 and Yak1 phosphorylate both Hsf1 and Msn2 upon glucose starvation^75–77^ and our results suggest these roles also as part of the HSR. The opposite phenotypes measured for Sch9 CRISPRi strains further agree with the role of Sch9 in inhibiting Rim15^78^, Yak1^79^ and Hsf1 in starvation stress^80^. In line with our findings, Hog1 has recently been reported to phosphorylate Hsf1 in osmostress^81^. Interestingly, the human Hog1 MAPK orthologue p38 also phosphorylates and activates Hsf1 upon treatment with an Hsp90 inhibiting compound^82^.

To get insights into HSR-regulated processes, we determined PK interactors from phospho-proteomics^83^ and TF target genes from ChIP data^84^ (Supplementary Fig. S20). Interactors of HSR-modulating PKs were enriched for functions in mitogen-activated protein kinase (MAPK) signalling (p-value=3.3e-05) and cell cycle regulation (p-value=1.8e-03) (Supplementary Fig. S20d). Target genes of HSR-regulating TFs had roles in responses to heat and oxidative stress (p-value=7.2e-04), the fungal cell wall (p-value=6.5e-03) and trehalose metabolism (p-value=4.3e-03) (Supplementary Fig. S20e). This target-based analysis thus not only proved useful in recapitulating paramount mechanisms of heat resistance that are remodelled as part of the HSR^18^, but also implies that these processes are, at least partially, controlled by the same TFs that regulate chaperone expression.

### Modulators of thermotolerance

We were curious if genes adjusting the HSR pathway have stress-protective roles. We thus screened for thermotolerance, the cells’ ability to survive a sudden and lethal heat shock (Fig. 4a). Briefly, yeast populations were selected at 50°C for 150 min (Supplementary Fig. S21) and recovered without maintaining CRISPRi perturbations. Comparing sequencing barcodes before and after heat shock, we identified twelve genes to promote and three to counteract thermotolerance, with minor or no impact on fitness (Supplementary Fig. S22 & Supplementary Tab. 1). Most thermotolerance modulators also physically interact with each other (14 of 15) holding the potential for cross-talk to fine-tune mutual activities (Fig. 4b).

**Figure 4.**
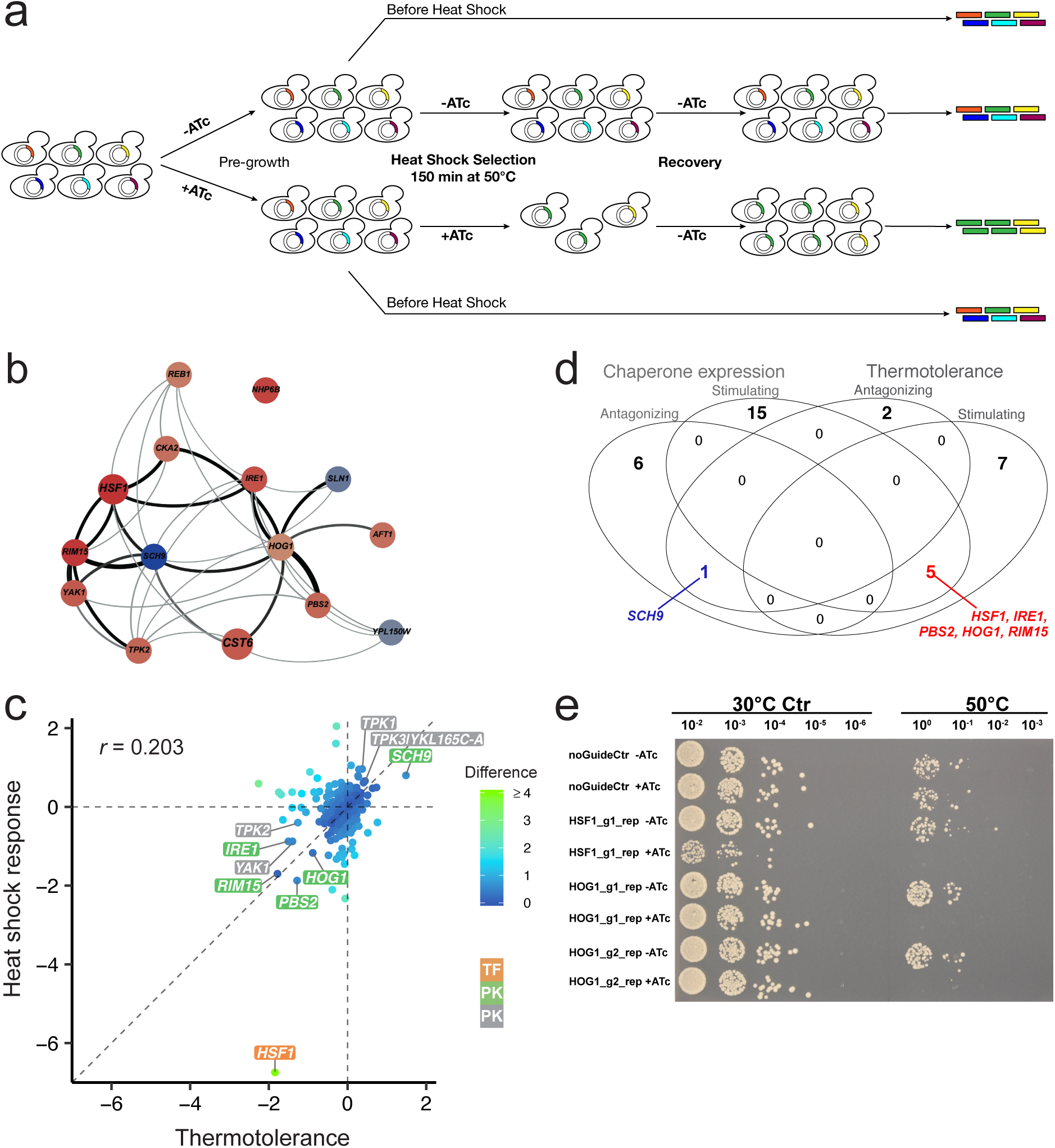
Shared regulation of thermotolerance and HSR. **a**, Schematic of the thermotolerance screen. **b**, Protein-protein interaction network of thermotolerance modulators based on STRING^108^. Line thickness indicates interaction confidence, and node size the screen p-value. Node colour denotes increasing (blue) or decreasing (red) roles on thermotolerance based on screen results with a light to dark gradient indicating low to strong effect size. **c**, Gene fold change comparison between the thermotolerance and HSR screens. Significant modulators are labelled in orange (TFs) or green boxes (PKs). Genes in grey boxes encode PKs that missed the stringent significance thresholds. Dashed lines are intercepts marking a fold change of 0 and the diagonal. **d**, Venn diagram showing overlap of genes altering thermotolerance and Hsp70 expression. **e**, Dilution spot plating of individual CRISPRi strains grown with and without ATc at 30°C, or exposed to 50°C for 150 min.

Decreased thermotolerance was observed for repression of three TFs that also had heat sensitive fitness (Hsf1, Reb1, Cst6), and for PKs in stress signalling (Yak1, Rim15) and the HOG (Hog1, Pbs2), UPR (Ire1) and PKA pathways (Tpk2) (Supplementary Fig. S22). Increased thermotolerance was measured for CRISPRi strains of Sch9, a PK controlled by target of rapamycin (TOR) signalling^85^, the Sln1 sensor kinase of the HOG pathway^86^, and the uncharacterized kinase Ypl150W^58^. Only three of the measured thermotolerance effects were known according to SGD, including the enhanced heat resistance of *sch9Δ* and the reduced tolerance of *pbs2Δ* and *rim15Δ* strains^43^. Strikingly, dilution spot plating of individual CRISPRi strains confirmed thermotolerance effects of Hsf1 and Hog1 (Fig. 4e), as well as for Tpk2, Pbs2, Cst6 and Rsc30 (Supplementary Fig. S23). Repression of the *TOR1|ARP3* locus was used as negative control that strongly decreased growth, but not thermotolerance. Interestingly, CRISPRa strains of Hsf1 had wildtype thermotolerance, suggesting that increased Hsf1 abundance may not alter heat resistance, and thermotolerance was decreased for activation of Pbs2 (Supplementary Fig. S23).

Genes found to modulate both thermotolerance and the HSR reporter modulators encode for Hsf1, stress-related kinases (Sch9, Rim15) and components of the UPR (Ire1) and HOG pathways (Hog1, Pbs2) (Fig. 4c & d). The Yak1 and Ypl150W PKs (Supplementary Fig. S24a & b), as well as the Tpk1/2/3 PKA subunits (Supplementary Fig. S25) were likely part of this overlap, although not fulfilling the strict significance requirements.

### CRISPRi effect magnitude depends on temperature and phenotype

We clustered effects across phenotypes to group genes by function (Fig. 5a), such as modulating fitness with moderate (clusters II, V, VI) or severe impact (VII & VIII) or adjusting the HSR pathway (II, III, IV, VII). Comparing CRISPRi magnitude across temperature, we found that effect size increased with temperature (Fig. 5b; Supplementary Fig. S26). Notably, the gRNA sequence GC content and secondary structure affects strain fitness with temperature-dependent contributions (Supplementary Fig. S27). Finally, we observed that phenotypes vary in their sensitivity to knock-down, as shown for Hsf1 (Fig. 5c). All six *HSF1*-targeting gRNAs severely decreased chaperone expression, implying that Hsf1 abundance is diminished strong enough to indirectly affect the assayed reporter. However, only the three most potent gRNAs affected fitness at 30°C, despite the essentiality of *HSF1* (Fig. 5c). Similar effect size gradients were observed for *UME6* and *RSC30* with only a few strong gRNAs (derived from HSR reporter impact) altering high temperature growth (Supplementary Fig. S28). Repression effects, even if they translate to protein level changes, do therefore not necessarily impact a robust downstream trait.

**Figure 5.**
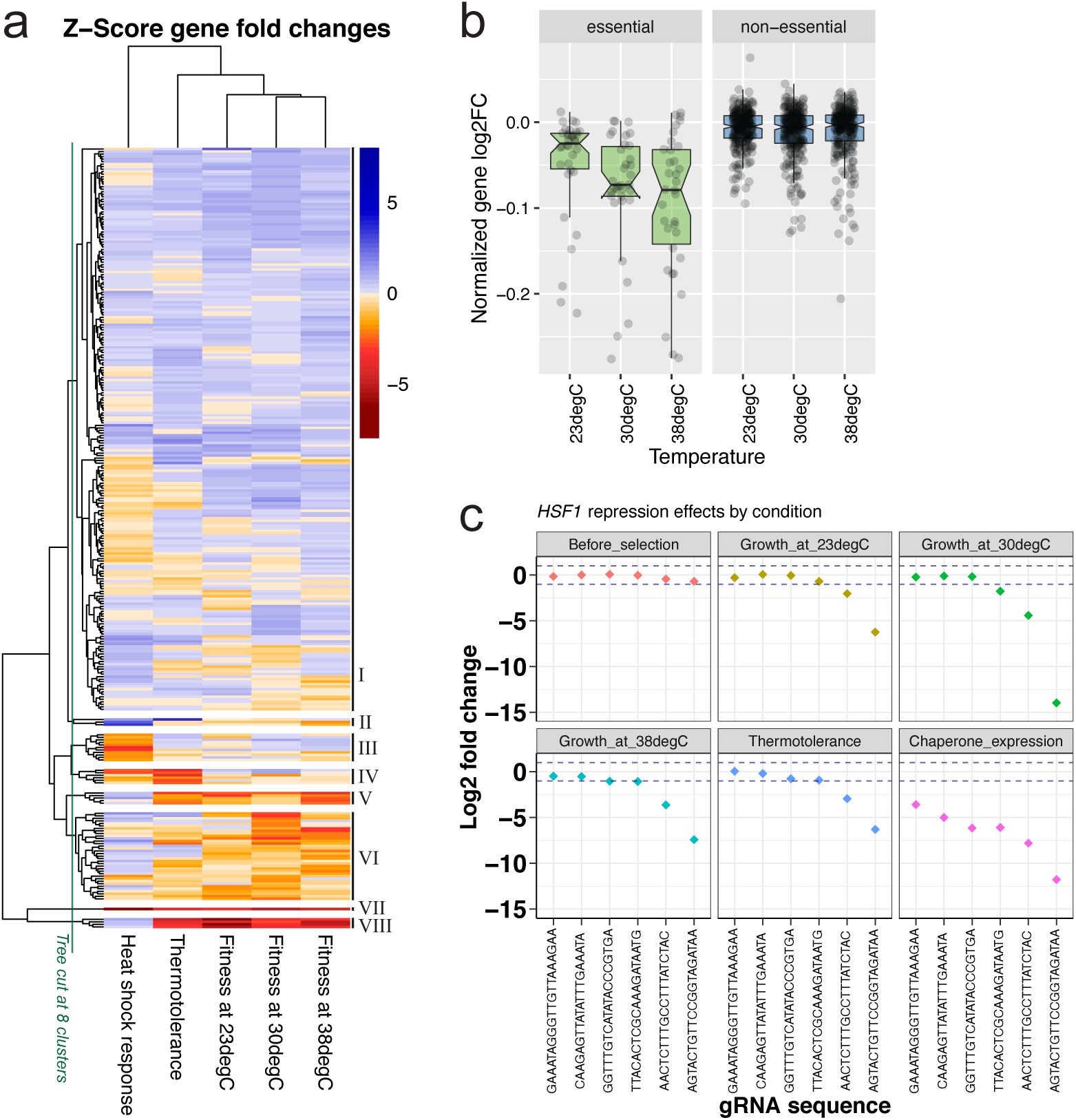
Knock-down sensitivity depends on growth and phenotype. **a**, Comparison of repression effects between screens as Z-score gene log2FCs (n=290). The hierarchical tree of rows is arbitrarily cut to form 8 clusters to group genes with shared effects, such as decreasing or increasing HSR activity (II or III, respectively), decreasing thermotolerance and HSR activity (IV), or severe fitness defects (VIII) upon repression. **b**, Temperature-dependent CRISPRi effects. Generation-normalized gene log2FCs of fitness screens at different temperatures, grouped for essential (n=34) and non-essential genes (n=255). c, Log2FCs (y-axis) of six HSF1-targeting gRNAs (x-axis) across screens. Dashed blue lines denote log2FC cut-offs 1 and −1.

## Discussion

We used CRISPRi screens to measure gene functions in temperature-associated growth, the heat shock response (HSR) pathway and thermotolerance. The integration of these diverse phenotypic readouts allowed us to reveal novel regulators of chaperone expression and link these to heat adaptation, discovering unknown functions of cell cycle regulators and the HOG, UPR and PKA pathways.

Our screens allowed us to pinpoint core regulatory components of the HSR capable of tuning thermotolerance by altering chaperone expression. We found central MAP kinases of the HOG pathway to stimulate thermotolerance and chaperone expression, thus explaining Hog1 activation in heat^38–40^ with a role in protein homeostasis (Fig. 6). We showed that the UPR-triggering sensor kinase Ire1^87^ increases the expression of a cytosolic Hsp70 (Fig. 4c, Supplementary Fig. S24). This suggests a role for the UPR in stimulating the HSR pathway, in line with the lowered activity of an artificial HSE reporter in the heat-exposed *ire1*Δ strain^88^. However, cytosolic HSPs were not previously known as UPR targets^89^. While PKA signalling is thought to counteract the HSR through inhibiting Msn2/4^90–92^ and by indirectly repressing Hsf1 presumably via Yak1 and Rim15^36,76,77^, we show that only the Tpk1 and Tpk3 subunits inhibit, while Tpk2 enhances Hsp70 production and thermotolerance (Supplementary Fig. S25a & b). This agrees with the increased Hsp12 expression and oxidative stress survival of *tpk1*Δ and *tpk3*Δ mutants, while both are decreased in *tpk2*Δ^67^. Additionally, differential Tpk1/2/3 roles are expected from their distinct physical interactors (Supplementary Fig. S25c) and phosphorylation patterns^40^.

**Figure 6.**
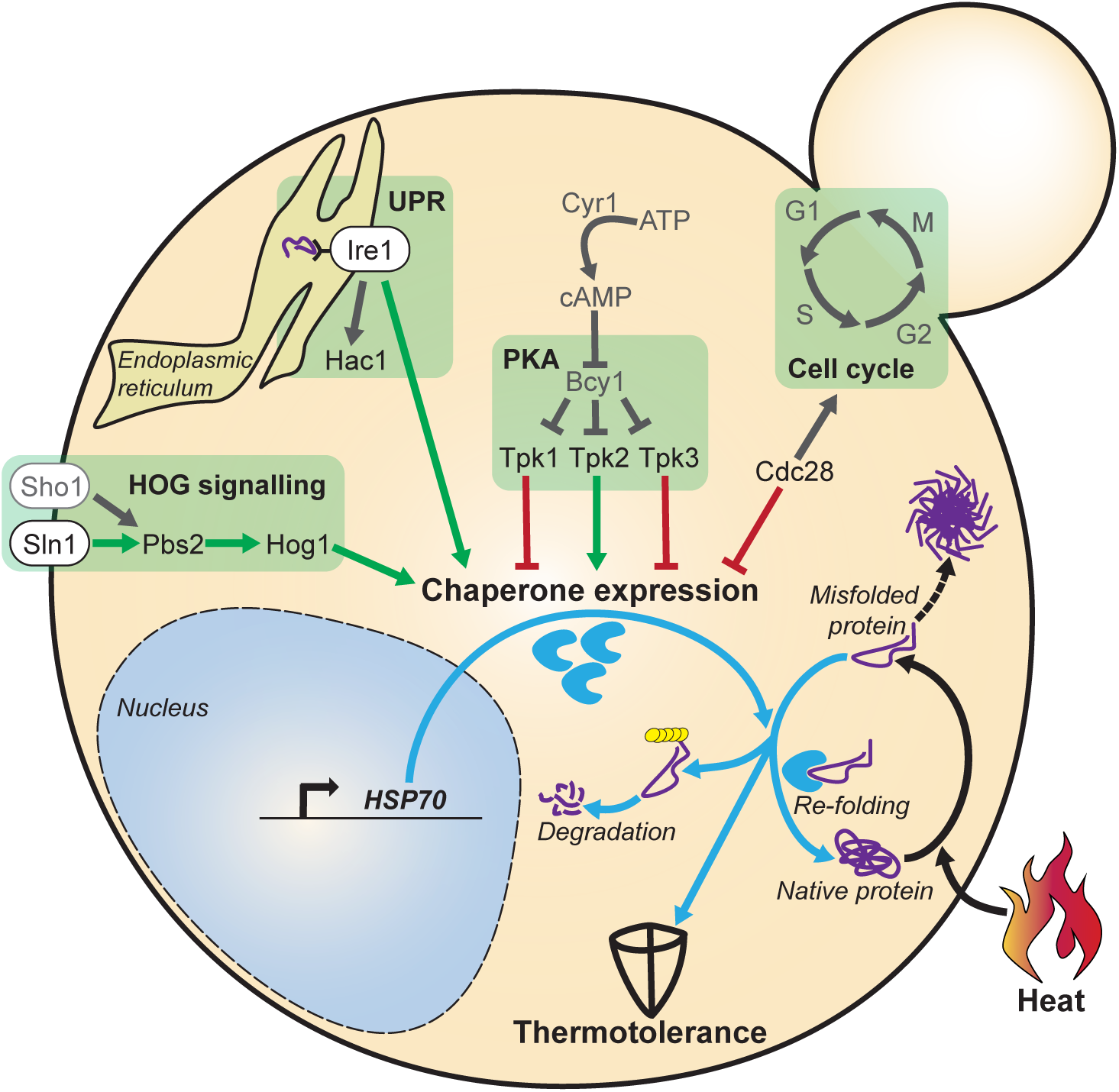
The HSR is regulated by multiple molecular pathways that sense diverse perturbations in different cellular compartments. Schematic overview of pathways controlling HSR activity based on screen-derived roles illustrated as stimulating (green arrows) or inhibiting (red stop indicators), and complemented by known processes from literature (in grey). Heat causes proteotoxic stress which is counteracted by cellular mechanisms, such as chaperone-mediated re-folding of misfolded proteins, and their degradation. Signalling pathway components of the HOG, UPR and PKA pathways modulate chaperone expression and thermotolerance. In addition, the roles of Cdc28 in promoting cell cycle progression and inhibiting HSR activity are shown. Further genetic roles measured in screens are omitted for clarity.

Notably, the strongest HSR-antagonizing PK (Cdc28) is the only cyclin-dependent kinase (CDK) both necessary and sufficient to drive the cell cycle in *S. cerevisiae*^93^. While cell cycle arrest is a known consequence of the HSR^94^, we show that key cell cycle regulators also signal back to the stress response in a mutually inhibitory manner (Supplementary Fig. S18; Fig. 6). During stress recovery, CDK promotes re-entry into mitosis and could simultaneously shut down the remains of a stress response to quickly resume proliferation. Supporting this, the HSP abundance is frequently regulated depending on cell cycle stage^95,96^. Additionally, given that Hog1 blocks mitosis by suppressing Cdc28-activators^97^ and its interacting cyclins in hyperosmotic conditions^98^, it presumably contributes to growth arrest also as part of the HSR.

We found repression effects to increase with temperature (Fig. 5b; Supplementary Fig. S26) which is likely due to both technical (CRISPRi system performance) and biological factors (sensitivity to knock-down caused by changes in transcription, translation and metabolism). We employed the *Streptococcus pyogenes* Cas9 fused to the human Mxi1 repressor which both evolved to operate around 37°C. Accordingly, Cas9 has higher *in vitro* DNA cleavage efficiency at 37°C compared to 22°C^99^. Temperature may also affect gRNA hybridization, as proposed for microRNAs^100^, in line with temperature-dependent contributions of gRNA GC content and secondary structure (Supplementary Fig. S27). This needs to be considered when using CRISPRi at extreme temperatures in yeast and other microorganisms. In our study, we paid special attention to potential biases in heat sensitivity, defined by contrasting fitness effects at 38 vs. 30°C. However, differences in strain dropout at these temperatures are minor compared to 23°C. The comparable mRNA depletion across temperatures in heat sensitive *HSF1* and *UME6* CRISPRi strains (Supplementary Fig. S15) and strong support by deletion strains reassured of measuring biological functions.

While repression is complementary to knock-out, we also measured novel fitness-modulatory roles (∼11% of 30°C fitness screen hits) which are unlikely to result from off-targets due to their support by two or more gRNAs. Additionally, effects were reproduced in screens at varied temperatures (Supplementary Fig. S14). We hypothesize that the sudden mRNA depletion can yield a stressed cell state while deletion mutants may trigger bypassing mechanisms when outgrown over several generations, such as acquiring adaptive mutations^101^. In CRISPR-KO screens, selection is usually performed immediately after introducing mutations and thus yield effects that are more severe^102^ or comparable to CRISPRi in human cells^103^, depending on assay setup. In future studies, it will be interesting to explore differential effects of knock-out and knock-down over time to explain the strengths and drawbacks of both techniques. Key advantages of the presented CRISPRi/a platforms are the inducible, reversible, and tunable modulation of transcription without altering genomic sequence and the ability to probe essential ORFs as demonstrated for *HSF1* (Fig. 1d & Fig. 5c).

A limitation of our HSR screens is given by the specificity of our reporter which monitors chaperone expression as a characteristic aspect of the HSR and may not report on other aspects. However, functional insights in PK phosphorylation targets and TF target genes illustrated the relevance of our screen hits for a variety of HSR-connected processes beyond HSP expression. Gene functions discovered by our screens can be directly applied in yeast biotechnology to generate strains with enhanced heat resistance or high temperature growth, i.e. to facilitate ethanol production^104,105^. If HSR-modulatory roles are conserved in human cells, they can be evaluated as therapeutic targets for disease treatment^7,10^. We anticipate that the established screen workflows and tools provide a basis to study diverse other reporters and molecular pathways, potentially expanding them to genome-wide scale and multiplexed application to infer genetic interactions and pathway connectivity.

## Material and Methods

### Chemicals, oligonucleotides, plasmids and strains

All chemical compounds, oligonucleotides, plasmids, and bacterial or yeast strains are listed in Additional File A1.

### Plasmid and strain construction

The Tet-inducible dCas9-MxiI and dCas9-nGal4-VP64 plasmids are available on AddGene (#73796 and #71128). Chemically synthesized gRNA oligonucleotide libraries were purchased from CustomArray, Inc. (GenScript) amplified by PCR and integrated into the NotI site of the pRS416 dCas9-Mxi1 plasmid via Gibson Assembly^106^ with 30bp homology regions, or ligation with T4 DNA Ligase. Single gRNA oligos and primers were purchased from Sigma Aldrich, and cloned into the NotI site of plasmids. Plasmids were transformed into *E. coli* NEB10beta chemo- or electrocompetent cells (New England Biolabs). Plasmid extraction from *E. coli* cultures was performed with the QIAprep Spin Miniprep kit (Qiagen) and was combined with FastPrep (MP Biomedicals) to break the cell wall of *S. cerevisiae* using ∼50μl volume of autoclaved glass beads (Sigma Aldrich) per cell pellet. FastPrep was run three-times at 5,5 m/s for 20s with 1min pausing. PCR was done using Phusion high fidelity polymerase (Thermo Fisher Scientific). Quality control of isolated DNA was performed with NanoDrop1000 (Thermo Fisher Scientific), Qubit spectro-fluorometer (Invitrogen) and High Sensitivity DNA Bioanalyzer chips (Agilent). For confirmation of sequence identity, plasmids and PCR products were submitted for Sanger sequencing to Eurofins Genomics. Template plasmids of PCR reactions were digested with DpnI for removal of bacterial DNA, and products were then used for Gibson Assembly or ligation. For library generation, multiple *E. coli* transformations were performed in parallel, pooled, and colonies of several selection plates were scraped together in a dense culture with LB-Ampicillin to ensure >20x coverage of libraries. Chemical transformation of *S. cerevisiae* was done as described previously^107^. Transformants were selected on synthetic complete uracil-dropout media (SC-Ura) agar plates. Single colonies were picked, confirmed by colony PCR and cultured for individual strain experiments. Transformations were sequence-verified by Sanger sequencing. Cell libraries were generated by washing off transformant colonies when reaching small size ∼36h after plating. For long-term storage at −80°C, 25% glycerol stocks were prepared.

### Plate reader growth and fluorescence assays

Sequence-verified strains were cultured in 96 well round bottom plates (Thermo Fisher Scientific) filled with 100 μL in SC-Ura dropout media with or without 250 ng/ml ATc. Yeast cultures were inoculated with OD600=0.005 for growth measurements or OD600=0.3 for fluorescence measurements (GFP channel: 488 nm excitation and 512 nm emission wavelength) in 15 min intervals with Genios (Tecan) or Synergy HTX (Biotek) plate readers according to the manufacturer’s instructions. The maximum growth rate and the time until half-maximum OD600 were determined by fitting a linear model and calculating its slope using the cellGrowth R package (V. 1.30.0)^108^.

### RNA extraction from yeast cells and qRT-PCR

Exponential phase yeast cultures were diluted to OD600=0.3 and cultured with 250 ng/ml ATc or without ATc for either 4 or 6 hours at temperatures as indicated. Cells were collected using a vacuum filter device and instantly frozen in liquid nitrogen. Cells were then resuspended in Trizol (Invitrogen) and RNA was extracted using the Quick RNA Kit (Zymo Research) according to manufacturer’s instructions. RNA was reverse-transcribed to cDNA using SuperScript III (Invitrogen) with RNasin (Promega, Life Technologies), Oligo(dT)18 (Thermo Fisher Scientific) and a reverse primer specific for the 3’ region of guide RNAs (see Additional File A1). This cDNA was diluted 1:10 or 1:20 and then used for SYBR Green quantitative reverse transcription PCR (qRT-PCR) using PowerUp SYBR Green PCR Master Mix (Thermo Fisher Scientific) and the Applied Bio-Systems QuantStudio 6 Flex Real-Time PCR System (Thermo Fisher Scientific). Analysis was performed with R code as follows: The log2 fold change between +ATc samples and -ATc reference samples was computed as the negative delta delta Ct (-ddCt) with ddCt = ((average transcript Ct) - (average *ACT1* house-keeping control Ct) of +ATc condition) – ((average transcript Ct) - (average *ACT1* house-keeping control Ct) of -ATc reference condition). Average Ct values were calculated from triplicates. Primers were designed to yield products of 75-130 nucleotides and are listed in Additional File A1.

### Microscopy and image analysis

To quantify cellular reporter gene expression, cells were imaged with the Zeiss CellObserver microscope (Carl Zeiss AG) during cultivation at 30°C and after 5h exposure to 42°C. Images were acquired with the ZEN Black software, and analysis was performed with KNIME to quantify cellular GFP signals^109^ and R^110^ for data visualization.

### Choice of HSR reporter

In search of a suitable reporter for HSR activity, we prioritized genes by their stress-related expression^11, 111^. We evaluated 44 Green Fluorescent Protein (GFP) tag strains^112^ on fluorescence induction after heat shock and found the *SSA1-GFP* fusion strain to yield the highest and most stable reporter signal in this panel (Supplementary Fig. S29). To quantify *in vivo* HSR activity, and for all screens presented, we employed a diploid BY4743 strain^113^ harbouring a chromosomally integrated reporter, consisting of the highly heat-responsive *SSA1* promoter with a Δ-280 bp truncation^52^ which controls expression of a fast-maturing GFP^54^. This reporter was chosen over artificial TF-specific promoters^26,114^ to measure effects of various transcription factors that affect Hsp70 expression.

### Design of gRNA libraries

Guide RNA oligonucleotides were designed to target 161 TFs^57^ (retrieved from yetfasco.ccbr.utoronto.ca on 16/04/2014 using DNA-binding=1 and dubious=false parameters) and 129 PKs^56^ (retrieved from yeastkinome.org on 16/04/2014). Libraries consist of 885 and 668 gRNAs, respectively. Each gene was covered by up to six different gRNAs to minimize off-target effect calling (Supplementary Fig. S2a). Guide RNAs were designed considering the distance of their midpoint (of the 20nt target sequence) to the respective TSS^115^ and nucleosome occupancy^116^. Blast (blast.ncbi.nlm.nih.gov/Blast.cgi) and ECRISP (version 5.4)^117^ were used to check potential off-target binding sites in the yeast genome, allowing for two mismatches at most. The gRNA design pipeline was published as part of Smith et al. (2016) and is available at lp2.github.io/yeast-crispri/. Potentially regulated TSS in close proximity to the intentional target TSS (Supplementary Fig. S2b) are included in the gene name. Specifically, if two or more gRNAs designed for “gene1” potentially targeted the TSS of another “gene2” within 150 nt distance^47^, the target locus is annotated as “gene1|gene2”.

### Screens

Plasmid libraries targeting sets of either TFs or PKs were transformed and profiled individually. Screens were performed with 30 ml bulk yeast populations in 150 ml flasks. Populations were pre-grown at 30°C with or without 250 ng/ml ATc for 10 h (∼3 generations) before selection to enable acquisition of CRISPRi-mediated changes on protein and phenotype level. This pre-growth had minor effects on strain composition of populations (Supplementary Fig. S3) so that almost every gRNA barcode in the design (>98%) was probed.

### Competitive growth screens

For competitive growth selection, pre-grown cultures were diluted to OD600=0.005 and grown over 1.5–2 days at temperatures 23, 30 or 38°C. Fitness screens were performed with two replicates. Due to anhydrotetracycline instability at high temperature, we confirmed that the compound maintains its biological activity in the used concentration over at least 3 days at 38°C (Supplementary Fig. S30).

### HSR screens and FACS

For HSR screens, pre-grown cultures were diluted to OD600=0.3, exposed to 42°C for 5 h to induce the *SSA1pr-GFP* reporter and sorted with flow cytometry. Sorting was performed immediately after heat shock to measure effects during the stress as opposed to recovery. 250.000 – 500.000 cells within the top and bottom 5% of cellular GFP reporter intensity were collected by FACS. Gating was used to select cells representing the bulk population in forward and sideward scattering, and to exclude dividing cells (Supplementary Fig. S31). Sorted cells were recovered for ∼5 generations without CRISPRi induction and accounting for growth arrest. HSR screens were performed in three replicates. Flow cytometry was performed using a MoFlo cell sorter (Beckman Coulter Inc.), equipped with a 70 μm nozzle. A Sabre argon ion laser (Coherent Inc.), tuned to 488nm (200mW) was used as primary laser. Laser illumination, optical configuration and sorting parameters were optimized with Flow-Check fluorospheres (Beckman Coulter Inc.). Single cells were measured and sorted in purify one-drop mode. Data was acquired with Moflo Summit and analysed with R code^110^.

### Thermotolerance screens

For thermotolerance selection, cultures were diluted to OD600=0.3, exposed to 50°C for 150 min and recovered for ∼7 generations considering growth arrest and rate during recovery (Supplementary Fig. S21). Two samples did not pass quality control and were excluded from analysis (PK after heat shock +ATc Rep2 & TF after heat shock -ATc Rep2 in Supplementary Fig. S13). Thermotolerance screens were performed in two replicates for the TF and four replicates for the PK library.

### Next generation sequencing

QuBit (Thermo Fisher Scientific) was used to quantify extracted plasmid DNA. To amplify gRNA barcodes, PCR was performed with ∼5 ng plasmid DNA as template and primers that add inline barcodes and Illumina P5 and P7 adapters (listed in Additional file A1). PCR products of all samples were run on 1% Agarose gel with SYBR Safe (Invitrogen) to control DNA amount, size and purity. Gel bands of PCR products were excised and DNA purified with the MinElute kit (Qiagen). After DNA quantification by QuBit, equal amounts of PCR products were pooled. This sequencing library was size-selected with an eGel (Thermo Fisher Scientific) and controlled for purity on a DNA-Bioanalyzer high sensitivity chip (Agilent). Illumina sequencing was performed in paired-end and 75-100 base pairs read length on NextSeq500 machines with 15% PhiX spike-in.

### Sequencing data analysis

Sequencing data was demultiplexed with Jemultiplexer. Base calling quality was controlled with FastQC. Reads were trimmed and aligned to a reference FASTA file with DNA barcodes using the Burrows-Wheeler algorithm to compute read counts. Computational analysis was performed with the edgeR R package^118^ although other count-based packages can alternatively be used. Fold changes of fitness screens were calculated as contrasts of read counts between +ATc and -ATc populations that both underwent competitive growth selection (Fitness effect). Fold changes of the thermotolerance screen are ratios between +ATc samples after versus before heat shock (thermotolerance effect), and for the HSR screen denote ratios between +ATc samples sorted for high versus low cellular reporter intensity (HSR effect). For all screens, we provide sample correlations of read counts (Supplementary Fig. S13), as well as for the computed gRNA log2FCs (Supplementary Fig. S32) and gene log2FCs (Supplementary Fig. S33). Gene log2FCs were computed as the mean log2FC of gRNAs per gene and also be calculated without prior calculation of gRNA log2FCs directly from the geometric mean of gRNA reads per gene. A single analysis workflow thus enables computation of log2FCs and adjusted p-values/FDRs for genes and individual gRNAs (Supplementary Fig. S34). We benchmarked approaches to calculate gene scores. We report gene scores as mean log2FC of all gRNAs per gene since it performed better or as well as other measures, including median and rank-based scores (see Supplementary Fig. S35 for correlation plots and Supplementary Fig. S36 for receiver operating characteristics and precision-recall curves) and has higher robustness against noise and off-targets. Significant gene functions are supported by a gene log2FC with FDR<0.05 and at least two gRNAs with an absolute log2FC>=1 and FDR<0.05. Fold changes between non-induced samples (-ATc) are helpful to determine the background variation without CRISPR-based perturbation for each screen (Supplementary Fig. S37).

### Gene ontology (GO) enrichments and target gene analysis

GO enrichment analyses were performed using the gProfiler^119^ and gProfiler2 R packages^120^. Significant genes are queried with all target genes of the library as a statistical background. Major cellular processes and functions are reported as enriched with adjusted Benjamini Hochberg FDRs specified. TF target genes were determined based on Chromatin Immuno-Precipitation on chip (ChIP-chip) data^84^. Target genes bound by at least two TFs identified as significant modulators were used for GO enrichment, using the *S. cerevisiae* genome as background. Phosphorylation targets of protein kinases were determined using the phosphogrid 2.0 database^83^. Physical interactors of significant PKs were used for enrichment analysis with a background consisting of all identified *S. cerevisiae* protein kinase targets.

### Dilution spot plating thermotolerance assays

Individual CRISPRi/a strains were cultured in SC-Ura with or without 250 ng/ml ATc for 1 day, diluted to OD600=0.3 and exposed to a 50°C in a table incubator (Eppendorf AG) for either 150 min or 90 min as indicated. A dilution series was prepared in SC-Ura and 10 μl of each dilution was plated on SC-Ura agar plates for the recovery of cells that survived the treatment without CRISPRi induction. After 2-3 days incubation at 30°C, photographs of plates were taken and colonies counted.

### Computational preparation and visualization

Figures were prepared using Adobe Illustrator 2019. Data was processed in R (V. 3.4.1)^110^ with the tidyverse^121^ and dplyr^122^ packages, and plots were generated with the LSD (V. 3.0) ^123^, ggplot2 (V. 3.1.0)^124^ and ggally (V. 1.3)^125^ packages. Minimum free energies of RNA secondary structure were computed using the ViennaRNA package (V. 2.0)^126^. Networks were generated with Gephi^127^, using protein-protein interaction data from the STRING database^128^.

### Statistics

For screens, multiple testing adjusted p-values (Benjamini Hochberg FDRs) were calculated with standard edgeR functions^118^ as described. For microscopy data comparisons between sample populations, a two-sided Wilcoxon adjusted p-value was computed using the ggpubr (V. 0.3.0) R package^129^. Boxplots are shown with a middle line corresponding to the median, and the lower and upper hinges denoting the first and third quartiles, respectively. For all experiments with multiple data points, these represent distinct samples and not repeated measurements.

### Code availability

The KNIME image analysis workflow is available on request. The R code for screen analysis can be downloaded from https://github.com/IAmTheMatrix/CRISPRi_Screen_Analysis/.

### Data availability

Demultiplexed Illumina sequencing data has been uploaded to Gene Expression Omnibus, available through GSE155455. The raw read counts and computed fold changes of gRNA barcodes and genes are provided as Additional Files A2, A3 and A4, respectively. Source data underlying figures is provided in Additional File A5.

## Supplementary Information

Supplementary Figures S1-S37 and Supplementary Table 1 are provided as two pdf files. Additional Files A1-A5 are xlsx files listing reagents, strains, DNA sequences, providing raw and processed sequencing data and source data to figures.

## Acknowledgements

The authors are grateful to Tanja Specht and Marko Kaksonen (University of Geneva) for sharing yeast GFP collection strains, and to Marc Sherman and Barak Cohen (Washington University, St. Louis) for kindly providing the SSA1pr-GFP reporter strain. The authors thank Mart Loog (University of Tartu), Timo Mühlhaus and Michael Schroda (University of Kaiserslautern) for productive discussions, Lin Gen and Aaron Brooks for computational support, and the EMBL Advanced Light Microscopy, Genomics and Flow Cytometry facilities for their excellent support, especially Diana Ordonez, Malte Paulsen and Vladimir Benes. This work was supported by an Advanced Investigator grant from the European Research Council (ERC) under the European Union’s Horizon 2020 research and innovation programme (AdG-742804 to L.M.S.) by the Deutsche Forschungsgemeinschaft (DFG, German Research Foundation; project STE 1422/4-1 to L.M.S.), by Estonian Research Council (IUT 34-4 to L.P.), a Marie Curie International Outgoing Fellowship to L.P., and Wellcome (grant number 206194 supporting L.P.). C.J. was supported by an Interdisciplinary Sciences Fellowship of the Joachim Herz Foundation.

## Competing Interests

The authors declare no competing interests.

## Contributions

C.J. conceived the study, designed and performed experiments, analysed data and wrote the manuscript. A.J. assisted with thermotolerance screens. L.P. designed and J.D.S generated gRNA libraries. A.J., J.D.S., L.P. and L.M.S. provided valuable advice and revised the manuscript. L.P. and L.M.S. supervised the study.

## Notes

### Competing Interest Statement

The authors have declared no competing interest.

### Summary of Updates

Updated manuscript title in pdf file.

