## Supplementary Table T1 for "Gene dosage screens in yeast reveal core signalling pathways controlling heat adaptation"

**Supplementary Table 1. Significant CRISPRi effects across screens.**

Significant genes (gene fold change FDR<0.05 and at least two gRNAs with absolute log2FC>=1 and FDR<0.05) are shown across screens with their gene log2FC (mean of all gRNA log2FCs per gene) and their maximum gRNA log2FC. The ID specifies if the target gene encodes a TF or PK. Effects on heat sensitivity and thermotolerance denote that effects on either fitness at 38°C or heat shock survival are not explained by general fitness effects at 30°C. For multiple TSS potentially targeted, respective genes are reported and separated by a vertical dash. Rows are coloured green if containing an essential gene. Tables are ordered by gene log2FC.

**Fitness at 23°C**

| Gene | Gene log2FC | Max. gRNA log2FC | ID |
|---|---|---|---|
| STB5 | -0.35 | -1.07 | TF |
| SWI4 | -0.55 | -1.58 | TF |
| REB1 | -0.56 | -1.25 | TF |
| MMO1\|PHD1 | -0.57 | -1.62 | TF |
| GCN4\|YEL008W | -0.58 | -1.56 | TF |
| AFT1\|YGL072C | -0.63 | -1.33 | TF |
| MCM1 | -0.71 | -1.88 | TF |
| RDS1 | -0.73 | -3.36 | TF |
| PHO85 | -0.77 | -1.61 | PK |
| BUD32 | -0.78 | -2.78 | PK |
| TEA1 | -0.90 | -3.27 | TF |
| CLA4 | -0.93 | -3.76 | PK |
| CST6 | -1.11 | -3.34 | TF |
| CEP3 | -1.30 | -5.01 | TF |
| HSF1 | -1.54 | -6.23 | TF |
| IPL1\|RKM1 | -1.73 | -3.08 | PK |
| SCC4\|SPT15 | -2.46 | -5.70 | TF |
| HRR25 | -2.61 | -7.26 | PK |

**Fitness at 30°C**

| Gene | Gene log2FC | Max. gRNA log2FC | ID |
|---|---|---|---|
| PTK1 | -0.33 | -1.74 | PK |
| GIN4 | -0.36 | -1.43 | PK |
| CAD1 | -0.37 | -1.10 | TF |
| IKS1 | -0.38 | -1.49 | PK |
| RFX1\|YLR177W | -0.43 | -2.96 | TF |
| CAK1 | -0.48 | -1.86 | PK |
| NHP6A | -0.49 | -1.08 | TF |
| HOG1 | -0.51 | -1.98 | PK |
| TPK2 | -0.53 | -1.83 | PK |
| MMO1\|PHD1 | -0.55 | -2.62 | TF |
| STP4 | -0.56 | -1.74 | TF |
| PTK2 | -0.57 | -1.46 | PK |
| GCR1 | -0.59 | -1.82 | TF |
| KNS1 | -0.60 | -1.64 | PK |
| RSC30 | -0.68 | -1.55 | TF |
| STB4 | -0.70 | -1.79 | TF |
| STP1 | -0.71 | -1.73 | TF |
| SAT4 | -0.76 | -1.78 | PK |
| EKI1\|SWI5 | -0.78 | -2.05 | TF |
| RTG3\|SFT2 | -0.78 | -1.18 | TF |
| CDC15 | -0.80 | -3.42 | PK |
| STB3 | -0.81 | -5.07 | TF |
| DUN1 | -0.86 | -1.25 | PK |
| YCK3 | -0.88 | -1.48 | PK |
| MCK1 | -0.91 | -5.32 | PK |
| KIN28\|YDL109C | -0.91 | -2.71 | PK |
| TDA1 | -0.92 | -2.34 | PK |
| TEL1 | -0.92 | -2.14 | PK |
| SCH9 | -0.95 | -2.64 | PK |
| PKC1 | -0.99 | -2.99 | PK |
| NRG1 | -1.04 | -6.57 | TF |
| FHL1 | -1.13 | -2.76 | TF |
| SWI4 | -1.14 | -4.44 | TF |
| HAL5 | -1.15 | -3.56 | PK |
| GCN4\|YEL008W | -1.16 | -4.68 | TF |
| STB5 | -1.17 | -3.25 | TF |
| PHO85 | -1.17 | -2.84 | PK |
| YJL055W\|ZAP1 | -1.23 | -4.64 | TF |
| ARP3\|TOR1 | -1.27 | -7.12 | PK |
| CBF1 | -1.31 | -3.43 | TF |
| TBF1 | -1.34 | -5.73 | TF |
| MCM1 | -1.36 | -3.51 | TF |
| FUS3\|PEP1 | -1.40 | -4.49 | PK |
| RAD53\|YPL152W-A | -1.42 | -4.36 | PK |
| MPS1 | -1.43 | -3.29 | PK |
| CDC28 | -1.44 | -3.56 | PK |
| ANR2\|ELM1 | -1.45 | -1.98 | PK |
| CEP3 | -1.47 | -4.21 | TF |
| CDC7 | -1.51 | -3.11 | PK |
| HSL1 | -1.53 | -7.38 | PK |
| SFP1\|YLR402W | -1.53 | -2.66 | TF |
| TEA1 | -1.55 | -5.81 | TF |
| FPK1 | -1.58 | -4.41 | PK |
| CBK1 | -1.58 | -8.80 | PK |
| AFT1\|YGL072C | -1.59 | -2.94 | TF |
| RSC3 | -1.60 | -4.65 | TF |
| RAP1 | -1.67 | -5.83 | TF |
| CDC5\|YMR001C-A | -1.69 | -5.16 | PK |
| REI1 | -1.72 | -4.69 | TF |
| REB1 | -1.73 | -3.16 | TF |
| NHP6B\|YBR090C | -2.11 | -5.91 | TF |
| BUD32 | -2.26 | -7.47 | PK |
| CST6 | -2.33 | -7.74 | TF |
| CLA4 | -2.38 | -9.21 | PK |
| HSF1 | -3.44 | -13.98 | TF |
| IPL1\|RKM1 | -4.33 | -8.64 | PK |
| SCC4\|SPT15 | -4.61 | -9.69 | TF |
| HRR25 | -5.10 | -10.40 | PK |

**Fitness at 38°C**

| Gene | Gene log2FC | Max. gRNA log2FC | ID | Heat sensitivity |
|---|---|---|---|---|
| TBF1 | -0.44 | -2.07 | TF | - |
| TOR2 | -0.57 | -2.11 | PK | increased |
| NIT2\|SPT10 | -0.64 | -2.41 | TF | increased |
| GCR1 | -0.74 | -1.67 | TF | increased |
| HSL1 | -0.77 | -1.98 | PK | - |
| ANR2\|ELM1 | -0.79 | -1.18 | PK | - |
| RSC30 | -0.84 | -1.66 | TF | increased |
| UME6 | -0.87 | -1.79 | TF | increased |
| CDC5\|YMR001C-A | -0.91 | -2.38 | PK | - |
| YJL055W\|ZAP1 | -0.93 | -2.27 | TF | - |
| SFP1\|YLR402W | -0.94 | -2.14 | TF | - |
| CBF1 | -0.99 | -2.25 | TF | - |
| CDC28 | -1.08 | -3.38 | PK | - |
| CDC7 | -1.09 | -1.81 | PK | - |
| RAP1 | -1.12 | -3.52 | TF | - |
| BUD32 | -1.12 | -4.05 | PK | - |
| CLA4 | -1.16 | -4.19 | PK | - |
| CDC15 | -1.16 | -3.03 | PK | increased |
| CEP3 | -1.21 | -4.48 | TF | increased |
| SWI4 | -1.22 | -2.65 | TF | increased |
| AFT1\|YGL072C | -1.30 | -2.28 | TF | increased |
| MPS1 | -1.38 | -5.83 | PK | increased |
| REB1 | -1.49 | -3.38 | TF | increased |
| ARP3\|TOR1 | -1.62 | -5.68 | PK | increased |
| HRR25 | -1.66 | -4.47 | PK | - |
| RSC3 | -1.66 | -3.82 | TF | increased |
| CST6 | -1.94 | -4.34 | TF | increased |
| HSF1 | -2.36 | -7.42 | TF | increased |
| SCC4\|SPT15 | -2.55 | -5.87 | TF | - |
| IPL1\|RKM1 | -2.59 | -4.27 | PK | - |

**Thermotolerance**

| Gene | Gene log2FC | Max. gRNA log2FC | ID | Thermotolerance |
|---|---|---|---|---|
| SCH9 | 1.48 | 2.41 | PK | increased |
| SLN1 | 0.76 | 1.20 | PK | increased |
| YPL150W | 0.64 | 1.36 | PK | increased |
| CLA4 | -0.42 | -2.12 | PK | - |
| CDC15 | -0.52 | -1.71 | PK | - |
| BUD32 | -0.53 | -3.09 | PK | - |
| TDA1 | -0.67 | -1.86 | PK | - |
| PKC1 | -0.72 | -1.70 | PK | - |
| CEP3 | -0.85 | -3.47 | TF | - |
| HOG1 | -0.88 | -2.38 | PK | decreased |
| REB1 | -1.06 | -2.24 | TF | decreased |
| CKA2\|SLD7 | -1.16 | -3.25 | PK | decreased |
| AFT1\|YGL072C | -1.18 | -2.09 | TF | decreased |
| TPK2 | -1.26 | -2.53 | PK | decreased |
| PBS2 | -1.28 | -2.14 | PK | decreased |
| YAK1 | -1.40 | -2.12 | PK | decreased |
| CST6 | -1.41 | -5.41 | TF | decreased |
| IRE1 | -1.50 | -2.94 | PK | decreased |
| NHP6B\|YBR090C | -1.64 | -4.01 | TF | decreased |
| SCC4\|SPT15 | -1.69 | -3.37 | TF | - |
| IPL1\|RKM1 | -1.75 | -3.53 | PK | - |
| RIM15 | -1.78 | -2.48 | PK | decreased |
| HSF1 | -1.84 | -6.31 | TF | decreased |
| HRR25 | -2.26 | -4.61 | PK | - |

**Heat shock response**

| Gene | Gene log2FC | Max. gRNA log2FC | ID |
|---|---|---|---|
| UME6 | 2.05 | 3.29 | TF |
| CDC28 | 1.61 | 4.00 | PK |
| SOK2 | 0.95 | 1.34 | TF |
| SCH9 | 0.80 | 2.10 | PK |
| RIM101 | 0.77 | 1.23 | TF |
| MPS1 | 0.73 | 2.60 | PK |
| HRR25 | 0.59 | 1.59 | PK |
| ALK1 | -0.35 | -1.38 | PK |
| SWE1 | -0.57 | -1.51 | PK |
| CKA1 | -0.58 | -1.50 | PK |
| NHP6A | -0.73 | -1.71 | TF |
| STP3 | -0.81 | -1.44 | TF |
| IRE1 | -0.87 | -1.99 | PK |
| RIO1 | -0.90 | -3.77 | PK |
| RCK2 | -1.07 | -1.81 | PK |
| HOG1 | -1.17 | -4.59 | PK |
| MSN4 | -1.19 | -2.16 | TF |
| RTG3\|SFT2 | -1.22 | -2.05 | TF |
| PTK2 | -1.23 | -3.13 | PK |
| RSC30 | -1.27 | -2.18 | TF |
| MCK1 | -1.32 | -4.28 | PK |
| MSN2 | -1.45 | -2.64 | TF |
| RIM15 | -1.70 | -2.78 | PK |
| PBS2 | -1.87 | -3.53 | PK |
| HRK1 | -2.10 | -3.83 | PK |
| IKS1 | -2.33 | -3.79 | PK |
| HSF1 | -6.74 | -11.80 | TF |
