## Supplementary Figures S1-37 for "Gene dosage screens in yeast reveal core signalling pathways controlling heat adaptation": Supplementary_Figures.pdf

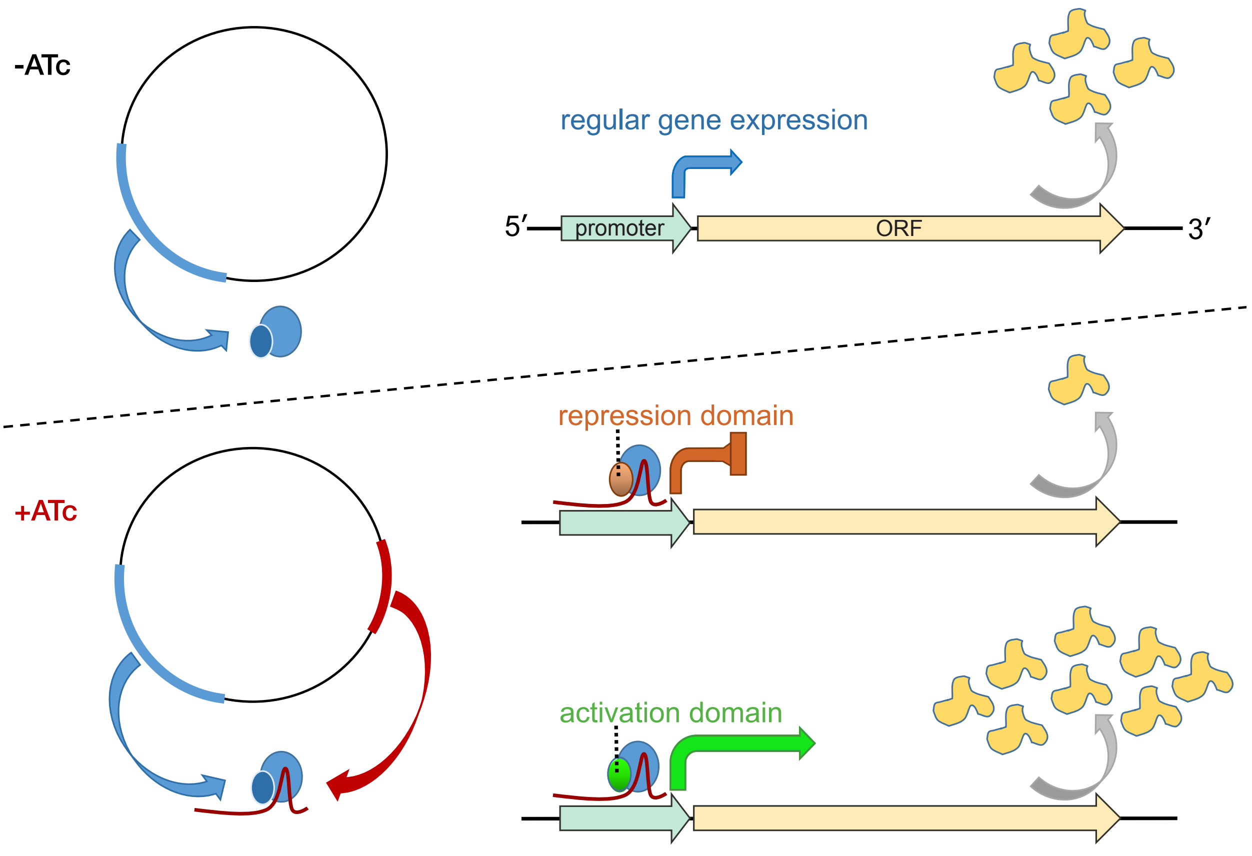


**Suppl. Fig. S1. CRISPRa/i mechanism.**
The CRISPRa/i systems are based on single plasmids. The dCas9-Mxi1 or dCas9-Vp64 components are constitutively expressed. In presence of Anhydrotetracycline (ATc), expression of the guide RNA is induced which can form a complex with the dCas9 proteins to localize them to a specific promoter region which is targeted for transcriptional activation or repression.

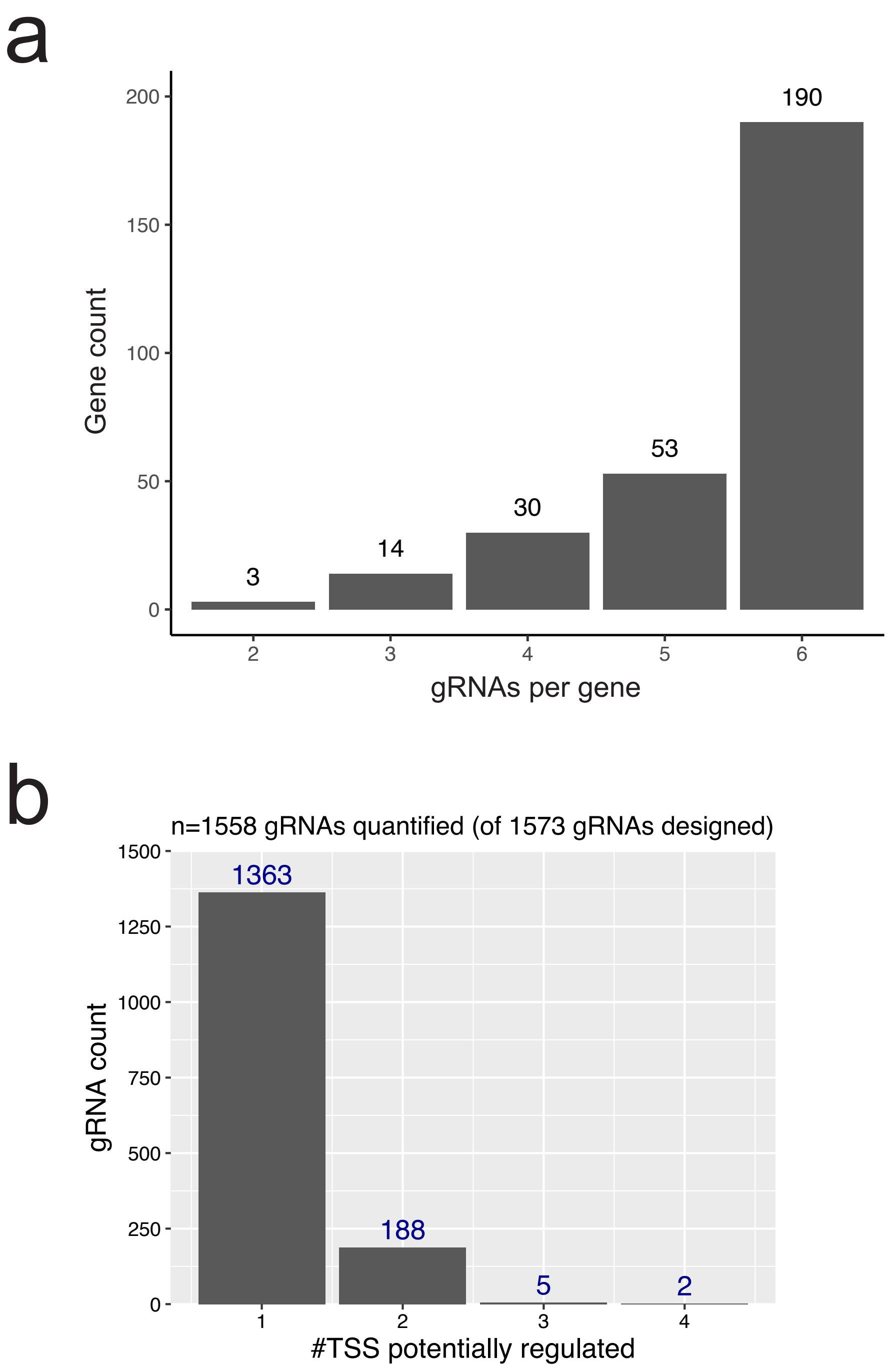


**Suppl. Fig. S2. Gene and TSS coverage.**
a) Guide RNA coverage of targeted genes (n=290). The number of target genes (y-axis) is shown with the number of gRNAs targeting the respective gene (x-axis).
b) Number of potentially regulated transcriptional start sites (TSS) for all gRNAs, based on the nucleotide distance between TSS and gRNA midpoint (see Methods).

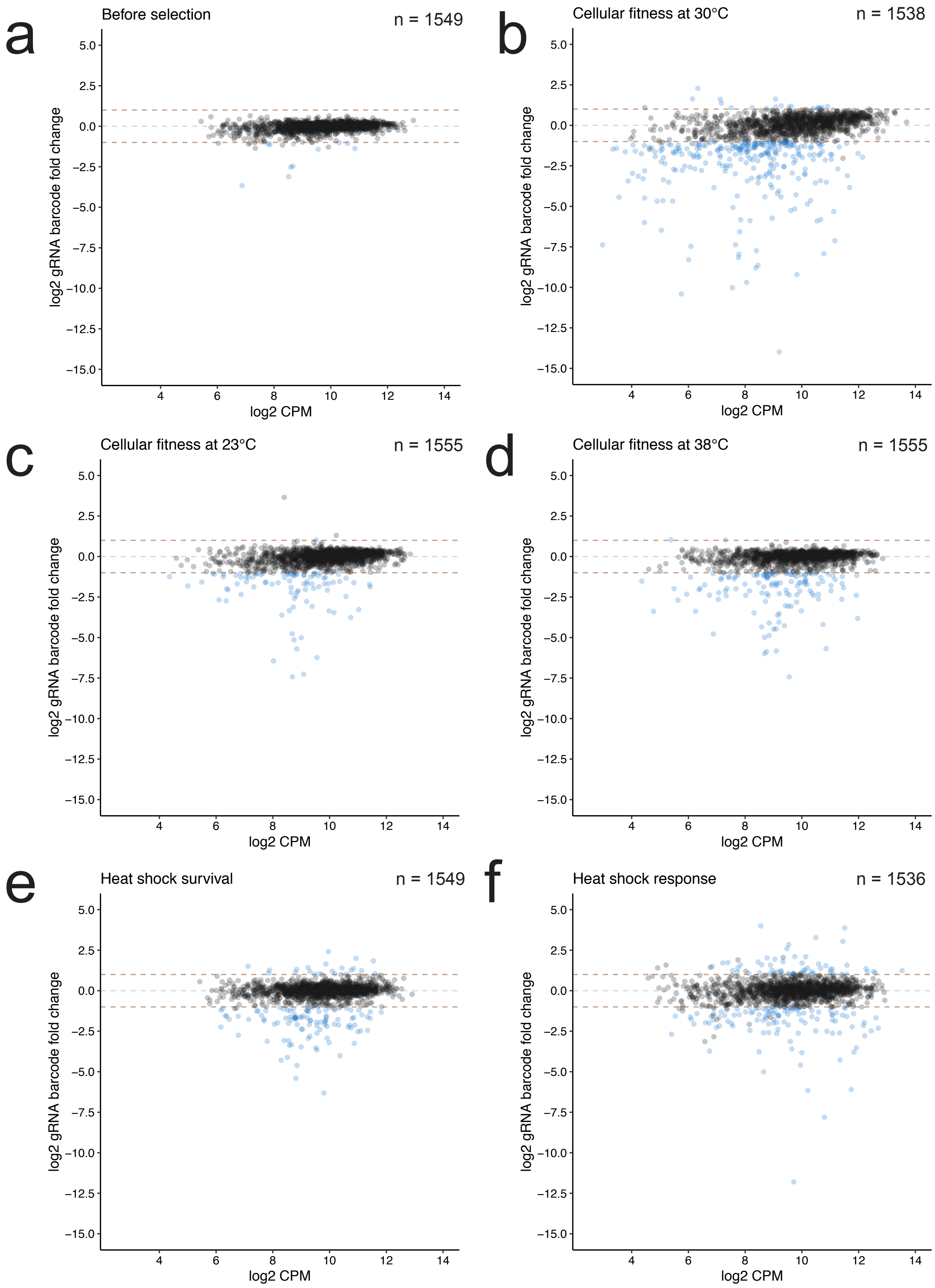


**Suppl. Fig. S3. gRNA fold change vs counts per million coverage (CPM).**
Guide log2FCs (y-axis) and coverage in counts per million (x-axis) shown across screen conditions. Blue dots denote gRNA barcodes (n = 1526 of 1573 strains quantified across all screens) with significant log2FC exceeding thresholds of +1 or -1 as marked by dashed brown lines. Dashed grey line marks the y intercept at log2FC=0.

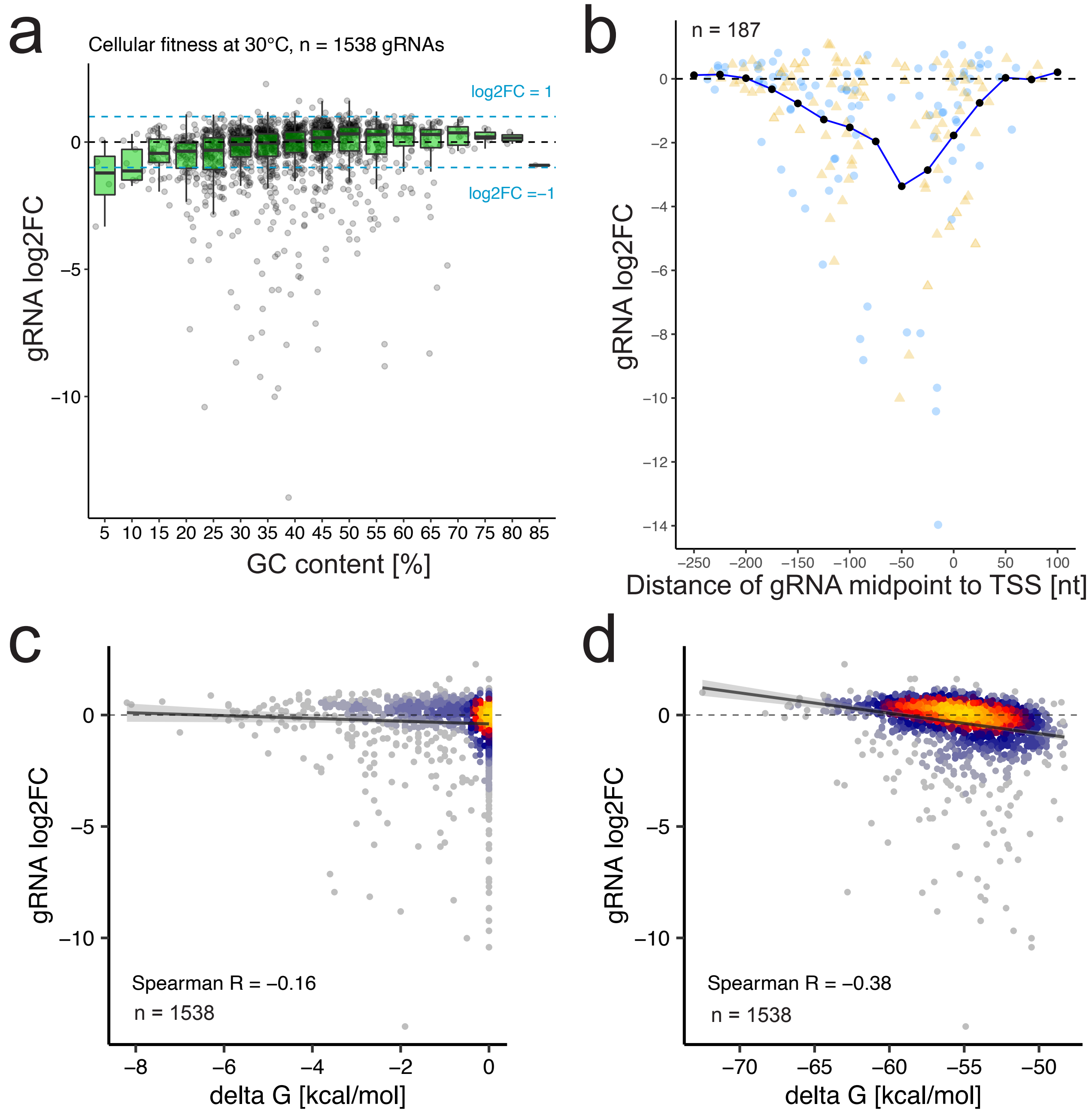


**Suppl. Fig. S4. Deriving gRNA design rules based on guide features.**
a) Log2FC related to the GC content of the genomic target region of gRNAs, for all gRNAs quantified in the 30°C fitness screen (n=1538). Boxplots show median gRNA log2FC and quantiles. Log2FC cutoffs of 1 and -1 are shown as dashed, lightblue lines. b) Log2FC related to the distance between the midpoint of the 20nt gRNA targeting sequence to the transcriptional start site (TSS) for essential gRNAs quantified in the 30°C fitness screen (n=187). Orange dots and blue diamonds denote gRNAs targeted to Watson and Crick DNA strands. Black dots and connecting line represent median values in bins of 50 nt that overlap by 25 nt. c) Log2FC related to the minimum free energy of RNAs secondary structure folds for the 20 nt genomic targeting region of gRNAs, for all gRNAs quantified in the 30°C fitness screen (n=1520). Minimum free energies (deltaG) are computed with Vienna RNAfold package (version 2.0) (Lorenz et al., 2011) for all gRNAs (n=1520). d) Similar plot as in (c), taking into account the 84 nt RNA-Polymerase III linker that may not be completely removed from gRNAs, as well as the gRNA structural region that may engage in folds.

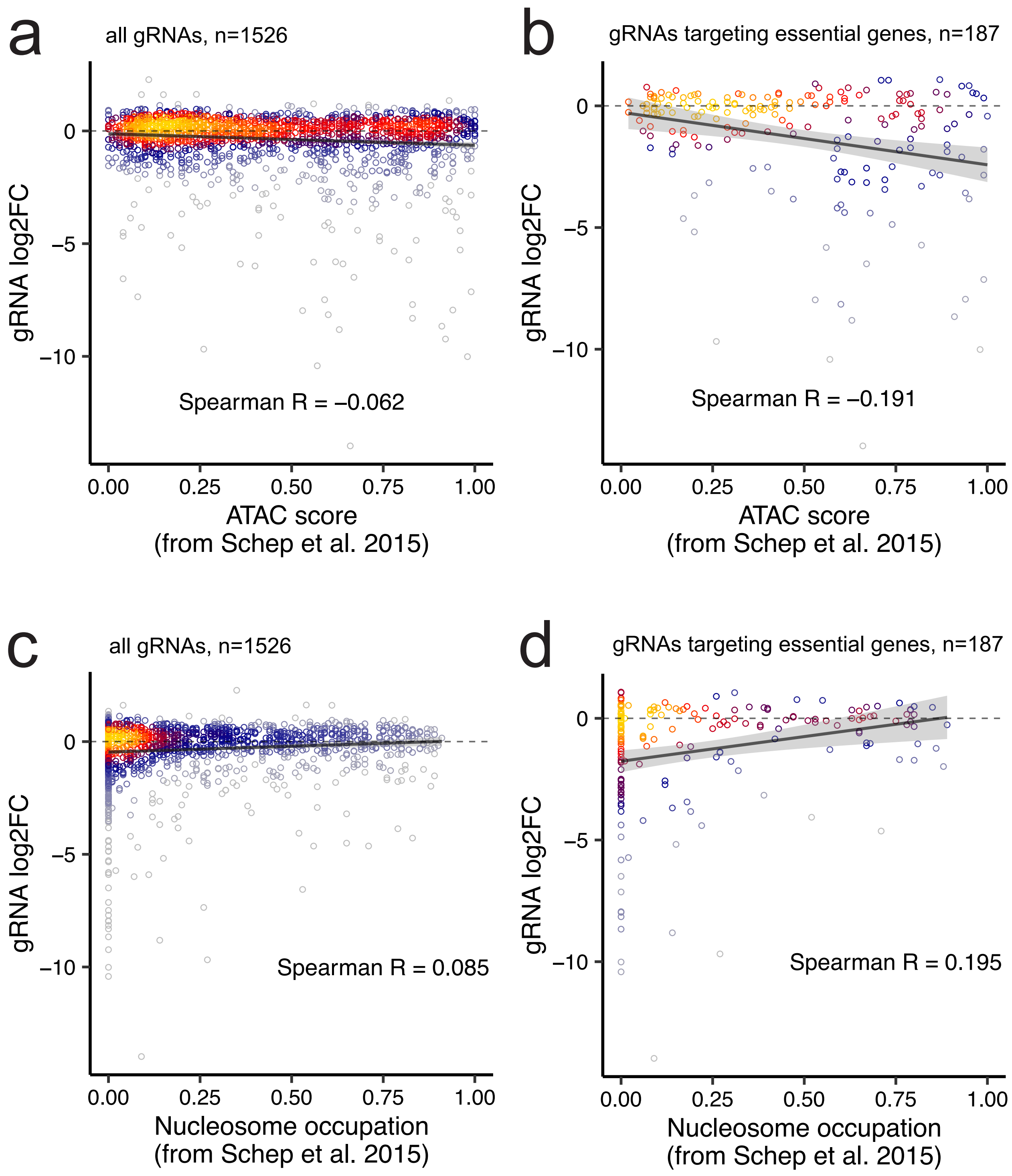


**Suppl. Fig. S5. Deriving gRNA design rules based on genomic target region features.**
a) Log2FC related to the ATAC-based local chromatin openness for all gRNAs quantified in the 30°C fitness screen (n=1520). ATAC score: ATAC-seq read counts averaged in sliding 50 bp range and normalized to the largest value within a 1 kilobase region. The ATAC score is a measure for chromatin openness relative to the neighbouring region. b) Similar plot as in (a) for all essential gRNAs quantified in the 30°C fitness screen (n=187).
c) MNase-seq derived nucleosome occupation for all gRNAs quantified in the 30°C fitness screen (n=1520). The nucleosome occupation score is a measure for the probability of nucleosome occupancy as defined in Schep et al. (2015). d) Similar plot as in (c) for all essential gRNAs quantified in the 30°C fitness screen (n=187).

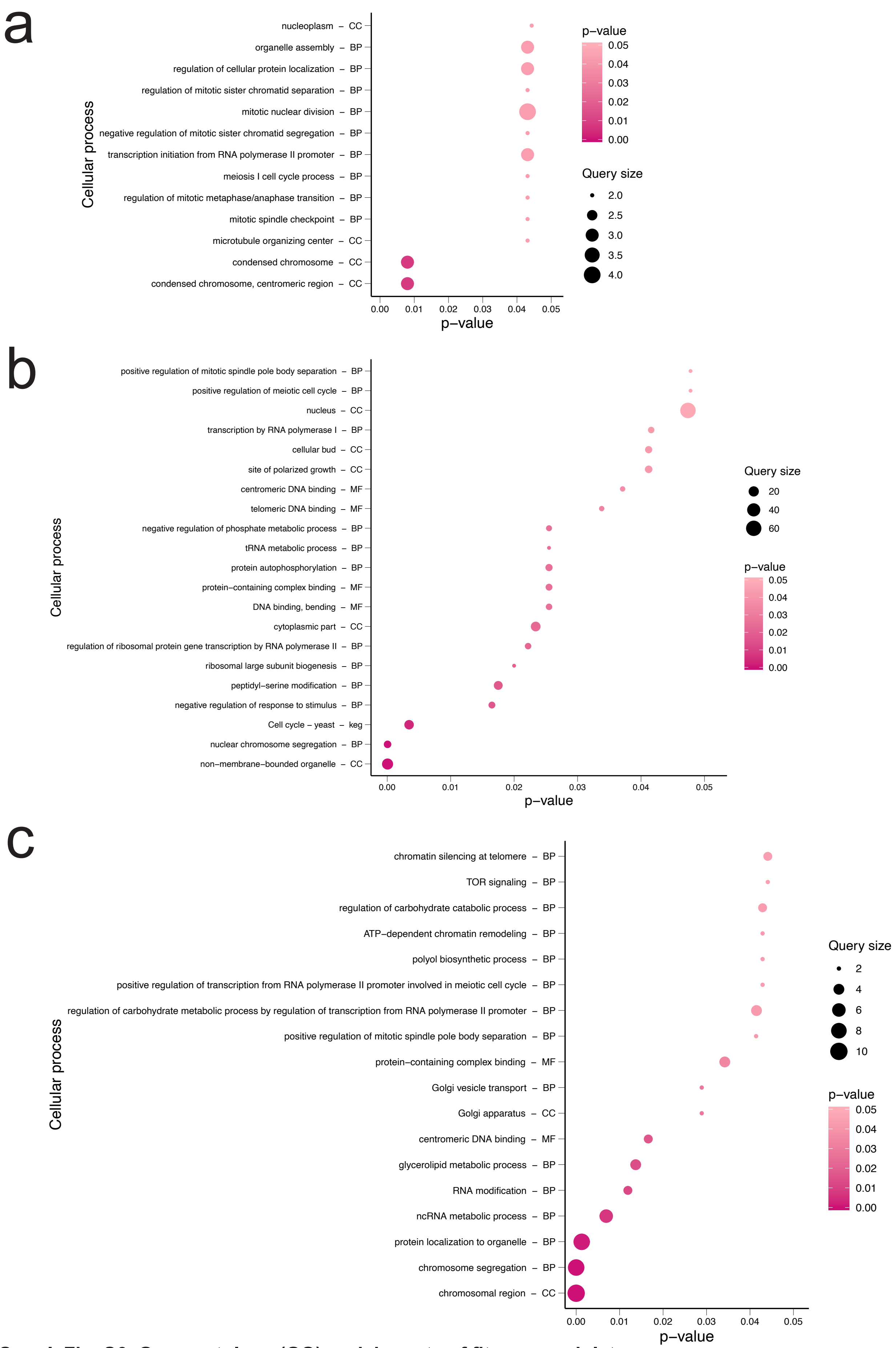


**Suppl. Fig. S6. Gene ontology (GO) enrichments of fitness modulators.**
GO enrichments of fitness-modulating genes are shown for competitive growth screens at a) 23°C (n=18), b) 30°C (n=68) and c) 38°C (n=30). GO enrichments were generated with all target genes as statistical background using the gProfiler R package (Reimand et al., 2016).

e

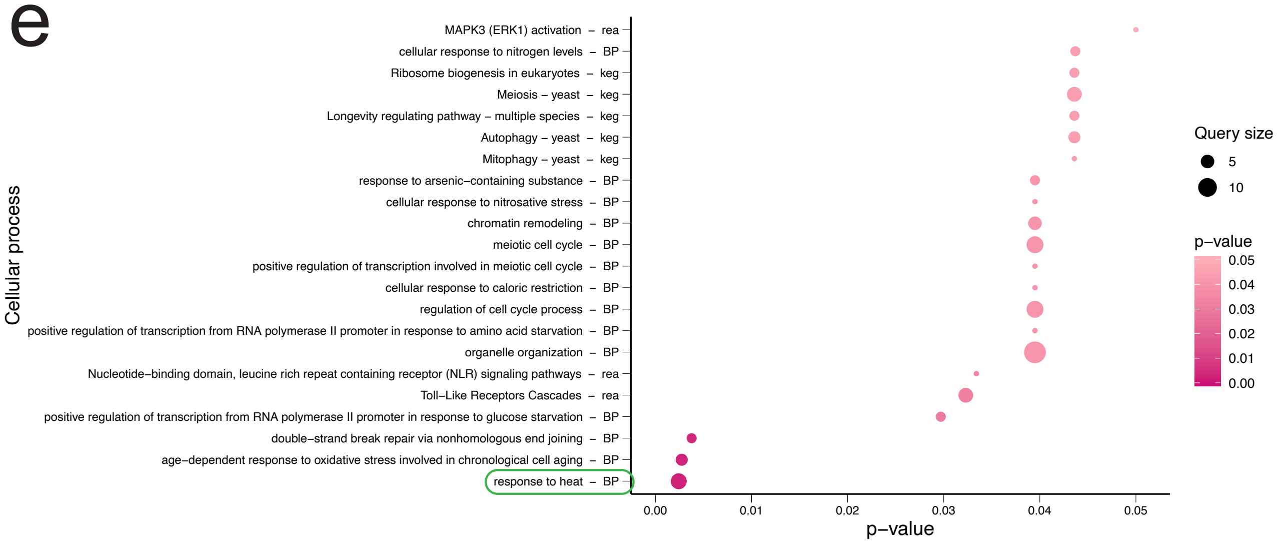


f

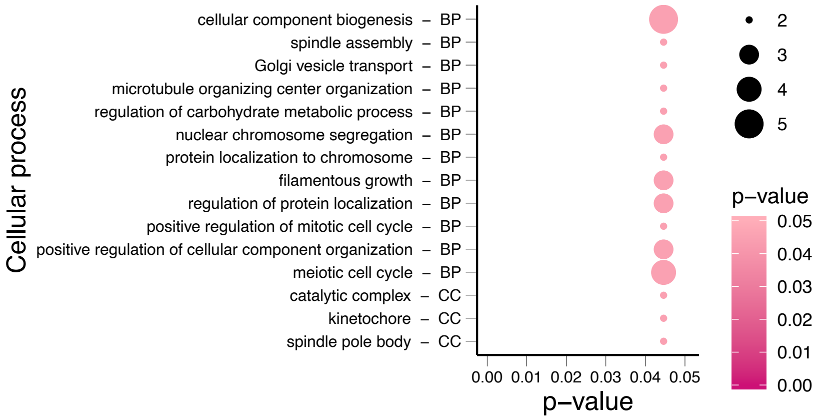


**Suppl. Fig. S6. GO enrichments of HSR and thermotolerance modulators. [continued for e & f]**
GO enrichments are shown for genes modulating e) the HSR (n = 27; GO term “response to heat” is marked) and f) thermotolerance (n = 15).

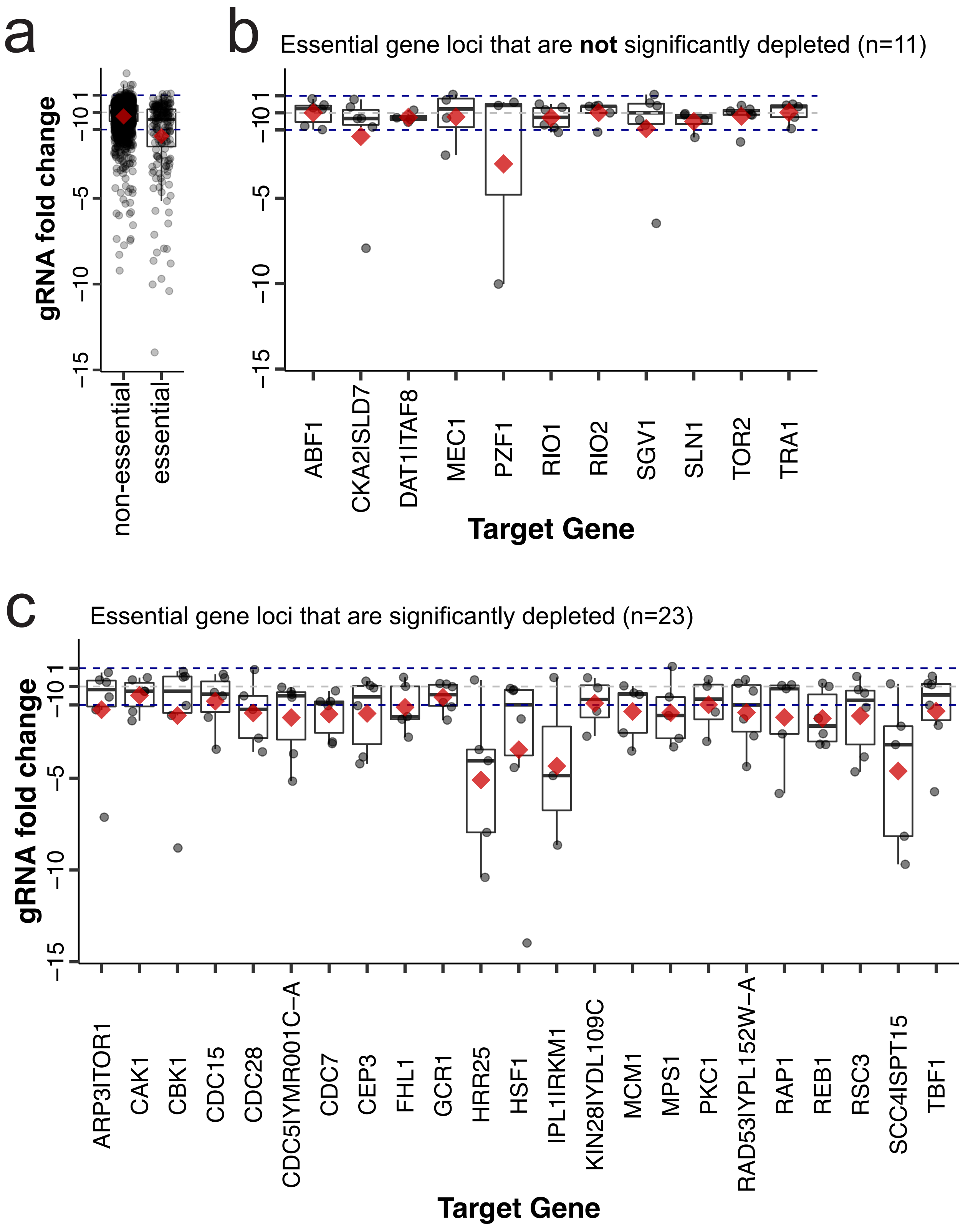


**Suppl. Fig. 7. CRISPRi strain depletion of essential genes in detail.**
Fitness effects on single strains (log2FCs of gRNA barcodes) are denoted as dots; for all non-essential and essential genes compiled (a), and in detail for all 34 essential genes targeted in the screens in separate groups of non-depleted (b) and significantly depleted genes (c). Red diamonds denote means, and for single genes in b and c represent the respective mean gene log2FCs. Boxplots show median gRNA log2FC and quantiles. Significant gene fold changes required an FDR <= 0.05 and support of at least two gRNAs with absolute log2FC >= 1 and FDR <= 0.05. Essentiality was defined by non-viable deletion mutants.

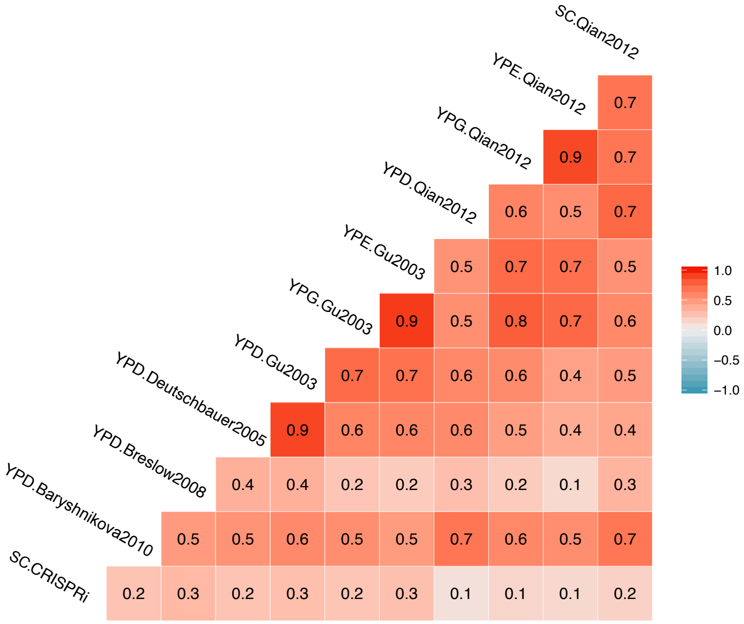


**Suppl. Fig. S8. Comparison of repression and homozygous deletion fitness effects.**
Pearson correlations between CRISPRi effects and several deletion collection screens are shown for all 290 target genes in the TF and PK libraries contained in the respective screens. Studies are indicated by the used growth medium, followed by first author name and publication year (Methods References 129-133). Assays vary in screen conditions and read-outs, as defined in respective articles. Scores were used as reported and without normalization.

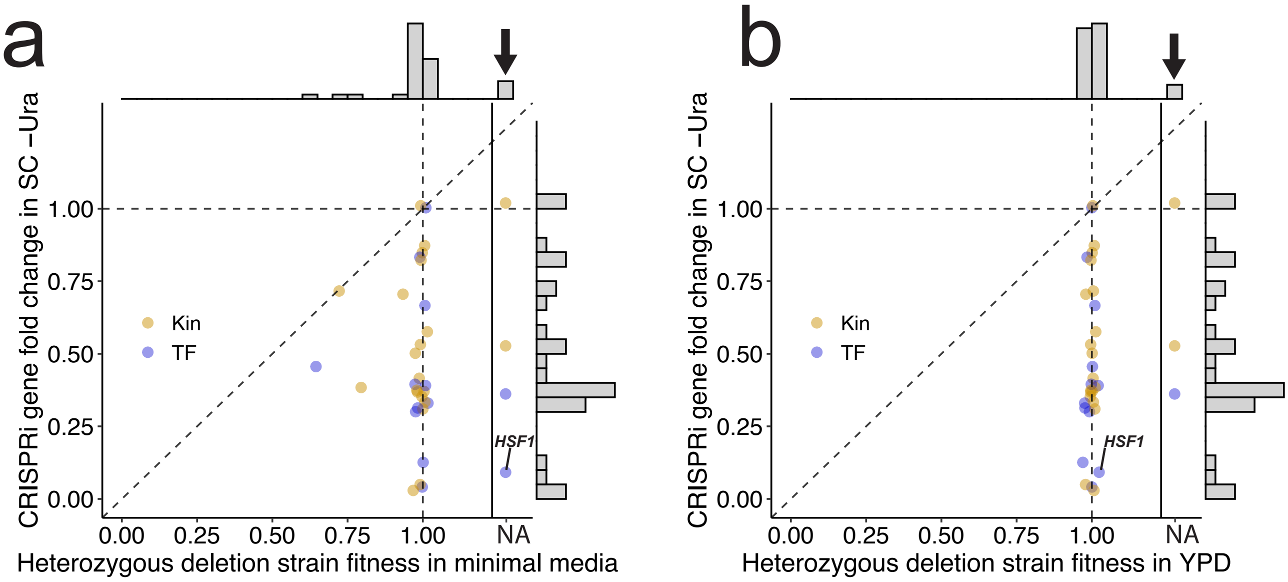


**Suppl. Fig. S9. Fitness effects of CRISPRi and heterozygous deletion strains.**
Comparison of CRISPRi gene fold changes of 34 essential genes (potentiated log2) with heterozygous deletion strain fitness from Deutschbauer et al. (2005) assayed in pooled format by DNA barcode hybridization in YPD (a) and minimal media (b). The black arrows point to four (a) and three essential genes (b) where values are missing in the data set of Deutschbauer et al. (2005) e.g. due to low coverage or the gene not being profiled. *HSF1* is labelled.

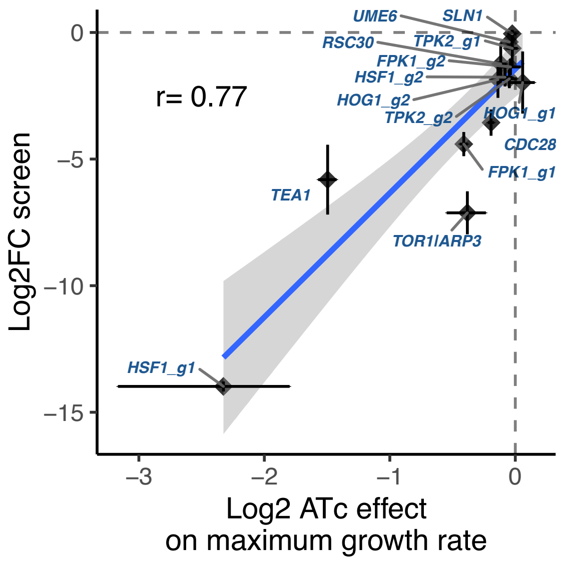


**Suppl. Fig. S10. Log2 gRNA fold change in screen versus log2 effect on growth rate in plate reader.** Comparison of screen guide log2FCs with log2-scale effects on growth rates determined from monoculture CRISPRi strains in plate reader OD600 measurements of triplicate wells (n=3), incl. Spearman correlation. Repressed genes are labelled and in cases where multiple guides are used in this or other experiments, gRNAs are labelled with g[number]. Dashed grey lines are intercepts marking a fold change of 0. Error bars denote standard deviations. The linear model fit is generated with the R ggplot2 function geom_smooth, using the default smoothing parameters with method=”lm”.

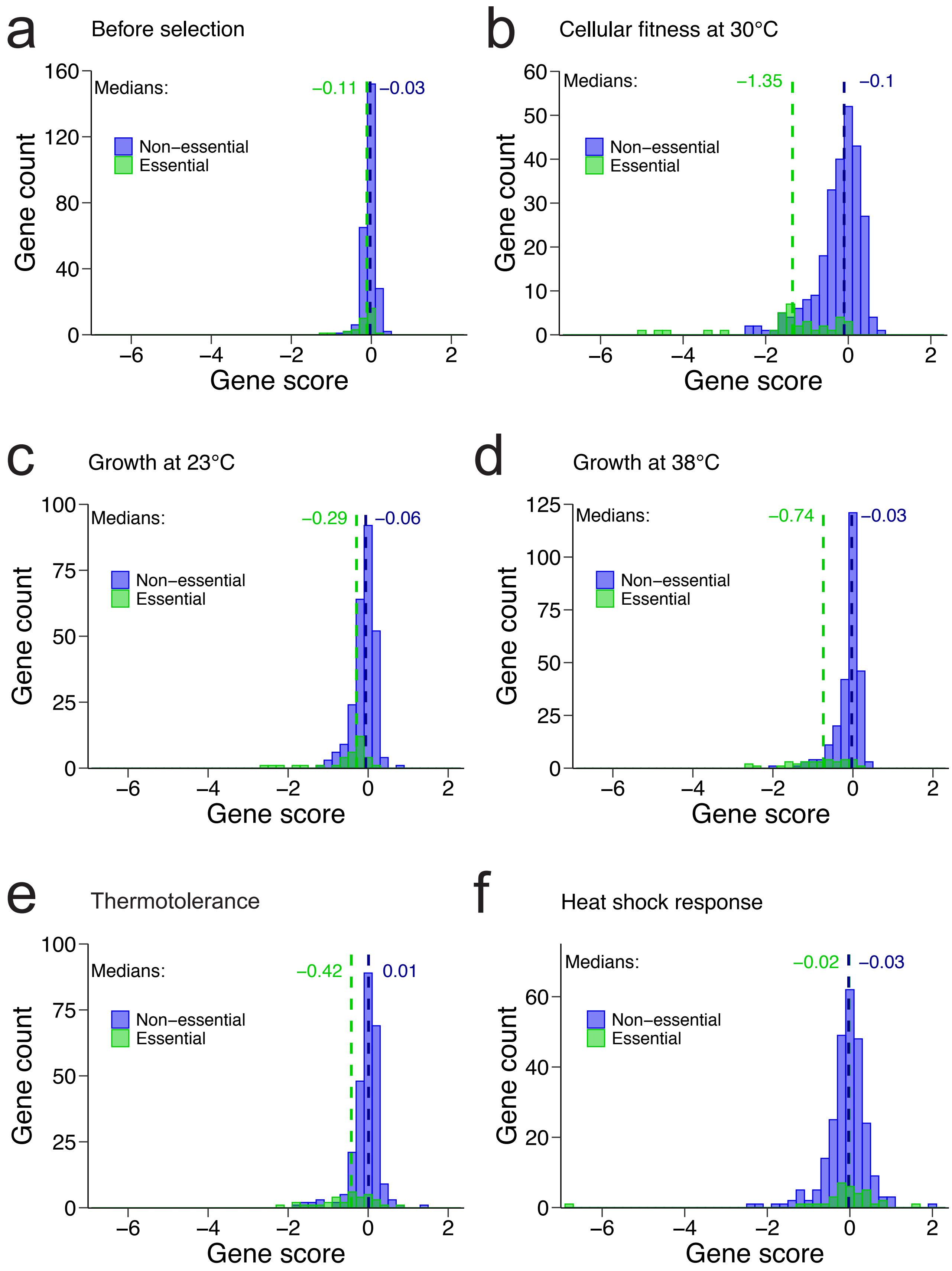


**Suppl. Fig. S11. Distribution of gene fold change distributions.**
Gene fold change distributions are shown for each screen, as indicated in the title for essential genes in green (n=34) and non-essentials in blue (n=256). Dashed lines indicate medians. Gene scores represent mean gRNA log2FC per gene. Histogram bins have size 0.2.

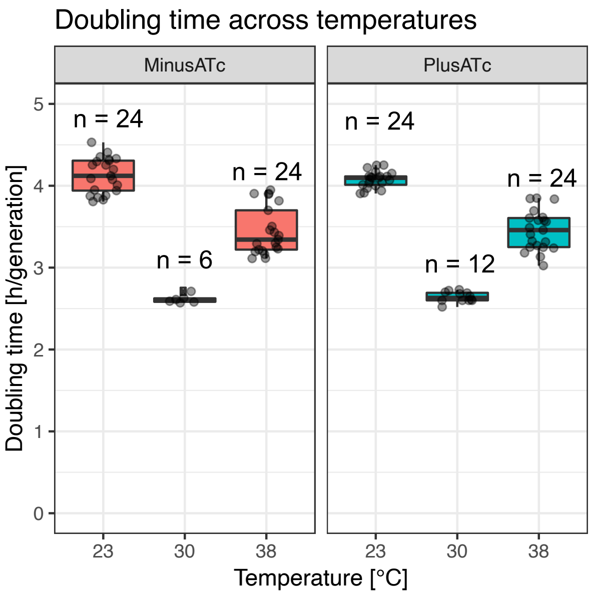


**Suppl. Fig. S12. Doubling times across temperatures.**
Doubling times at temperatures 23, 30 and 38°C were determined by measuring OD600 over time in 96-well cultures with a plate reader. The maximum growth rate (slope of linear fit) was calculated with the cellGrowth R package (Gagneur and Neudecker, 2019).

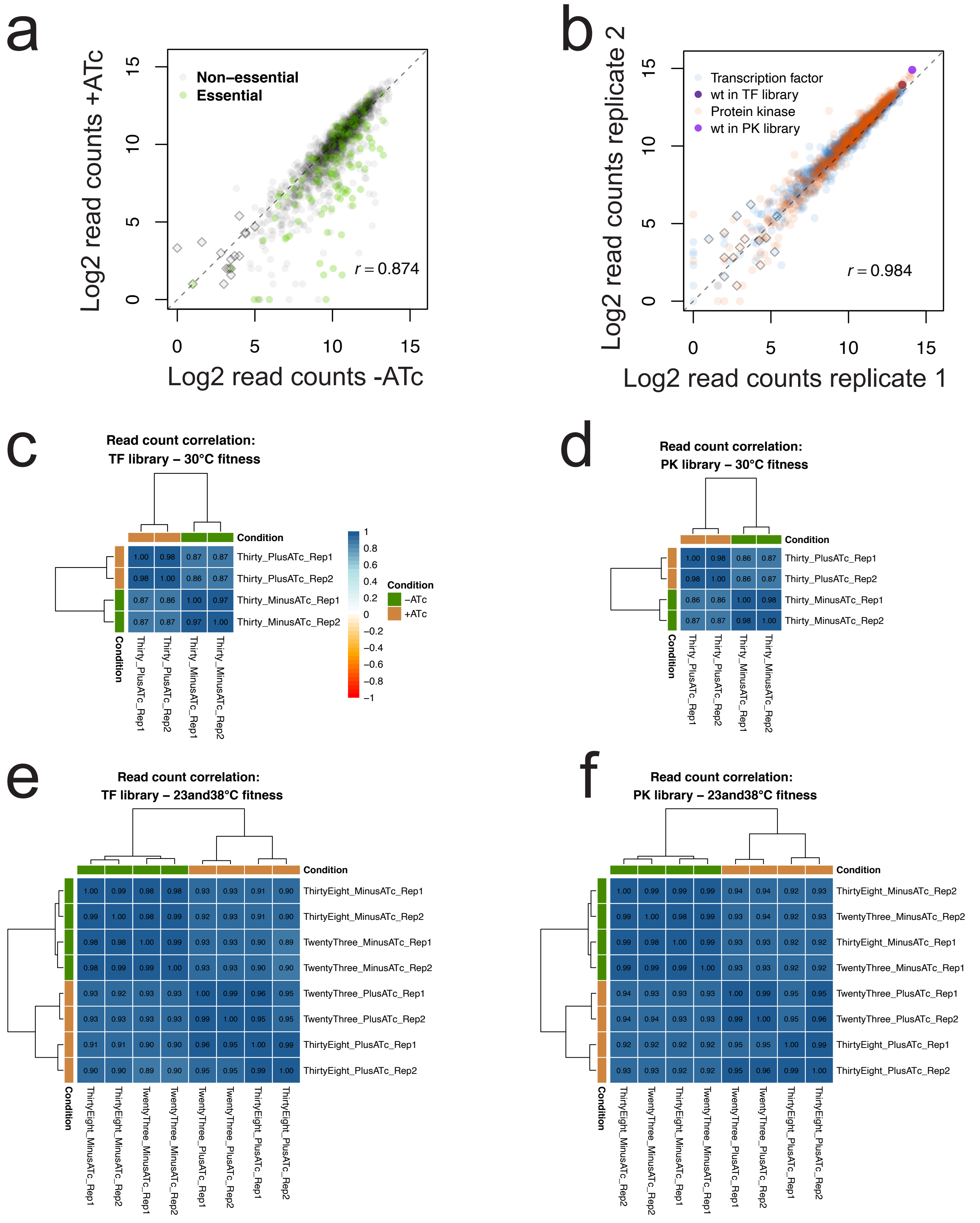


**Suppl. Fig. S13. CRISPRi effect and replicate correlation. [a-f]**
a) Log2-transformed barcode read counts of the 30°C screen, comparing CRISPRi-induced (+ATc) and reference samples (-ATc) for n=1573 gRNAs and two wildtype barcodes (no gRNA) of data merged from TF and PK library screens. Essential gene targets are marked green and low read count barcodes (filtered) are denoted as diamonds. Spearman correlation is shown.
b) Log2-transformed barcode read counts of the 30°C screen, comparing +ATc replicate samples. TF (blue) and PK library (orange) data is merged for n=1573 gRNAs and two wildtype barcodes (no gRNA). Diamonds mark low read counts (filtered) and wildtype barcodes are coloured in light and dark violet to show their enrichment during competitive fitness selection relative to other barcodes. Spearman correlation is shown.
Spearman correlations between read count samples of all fitness screens: Fitness at 30°C screen for c) the TF and d) the PK library samples, and for fitness at 23 and 38°C for samples of e) the TF and f) the PK library.

g

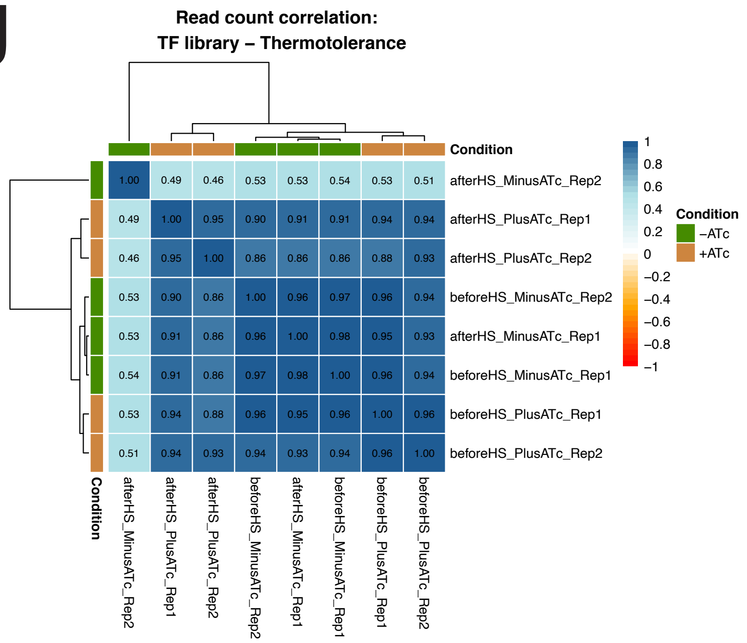


h

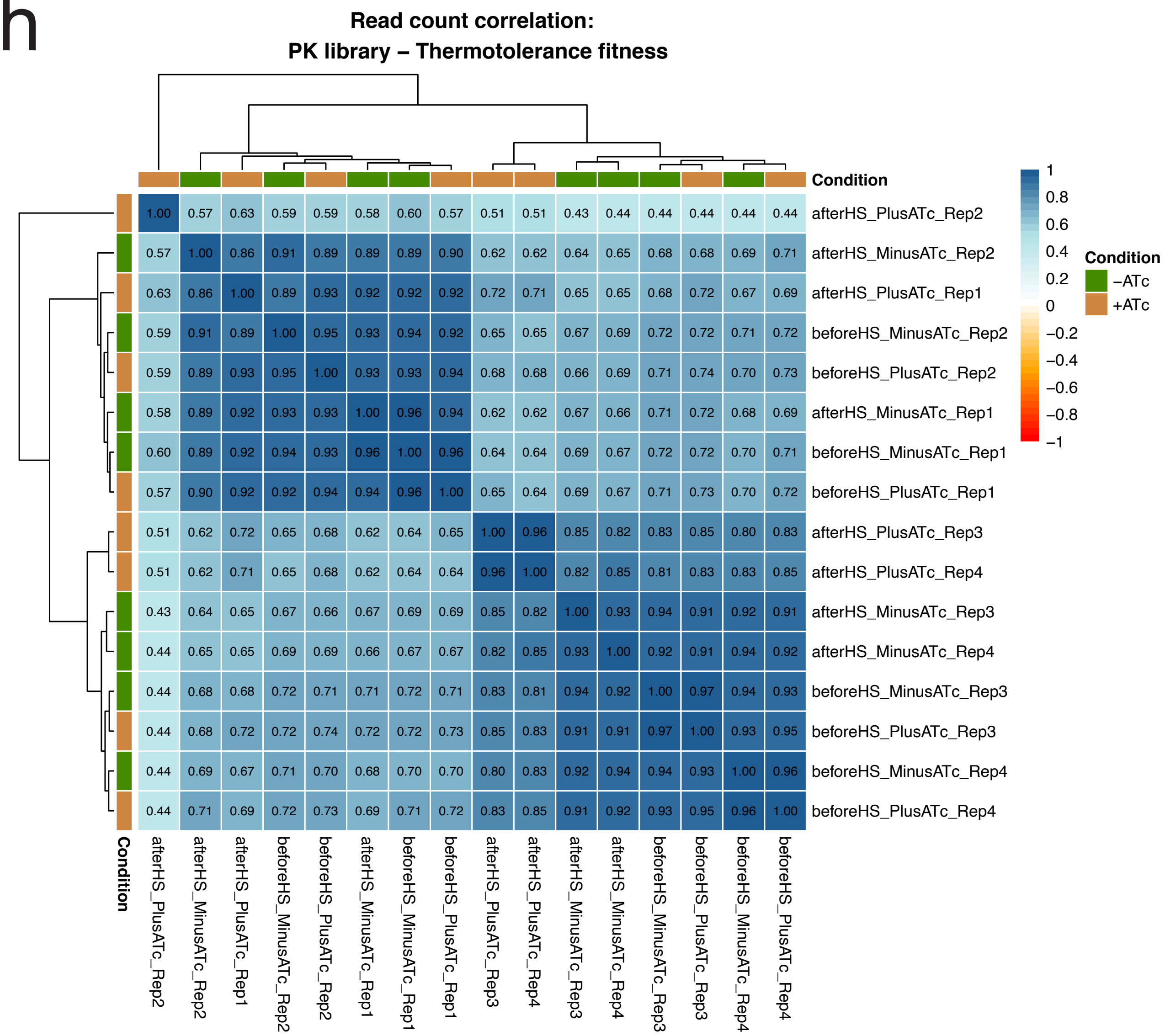


**Suppl. Fig. S13. Read count correlations. [continued for g and h]**
Spearman correlations between read count samples of the thermotolerance screen for samples of g) the TF and h) the PK library.

i

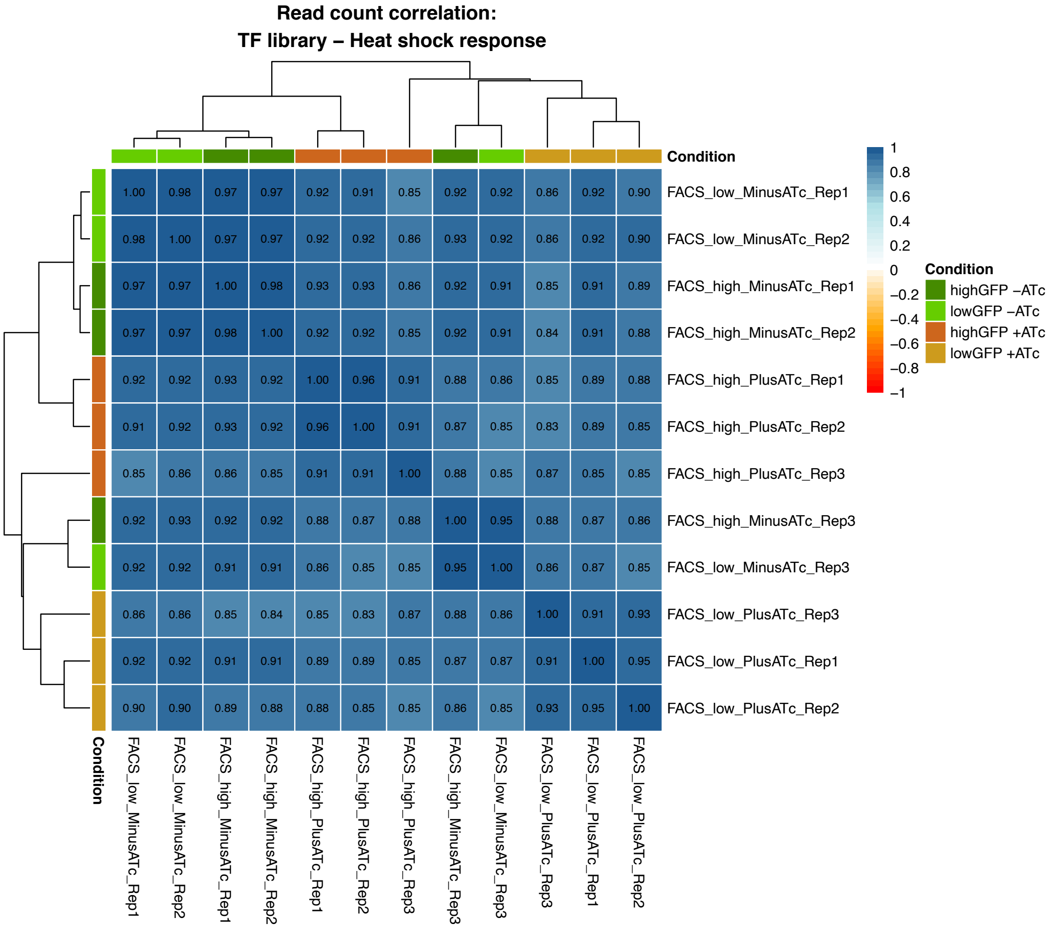


j

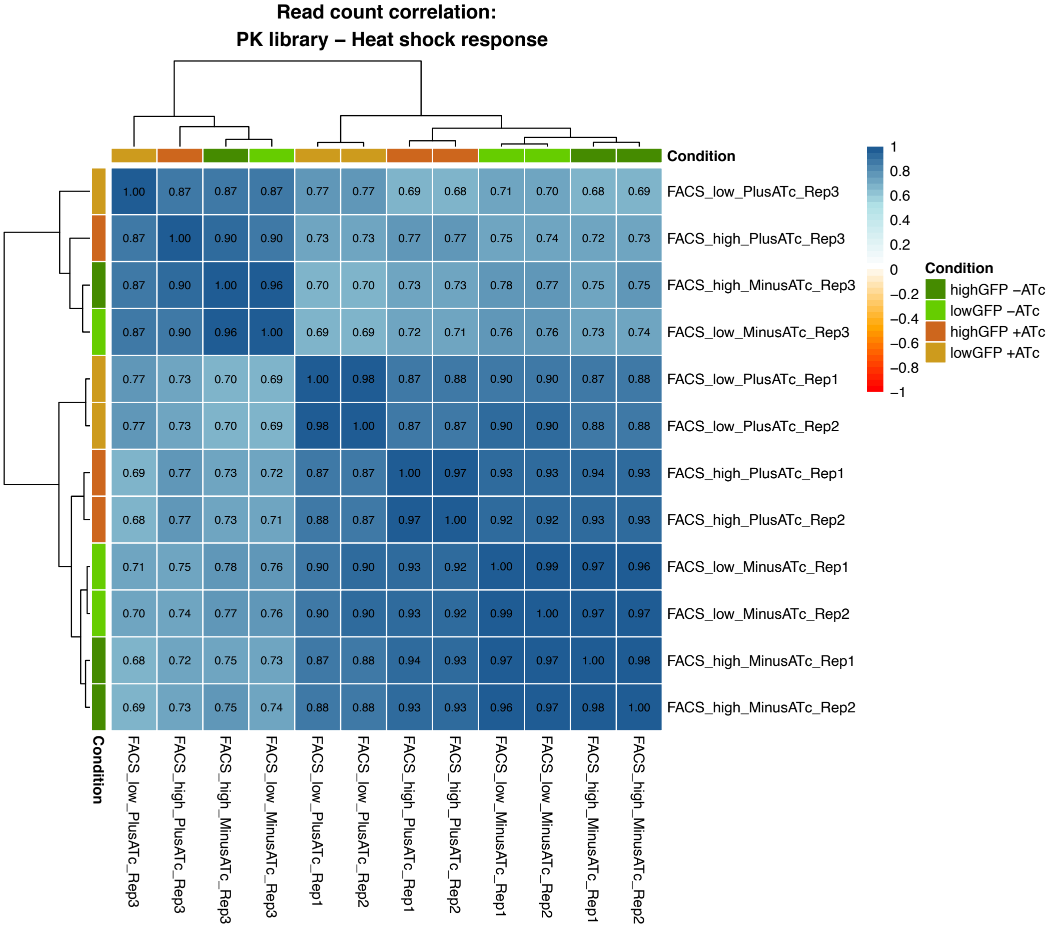


**Suppl. Fig. 13. Read count correlations. [continued for i and j]**
Spearman correlations between read count samples of the HSR screen for i) TF and j) PK library samples.

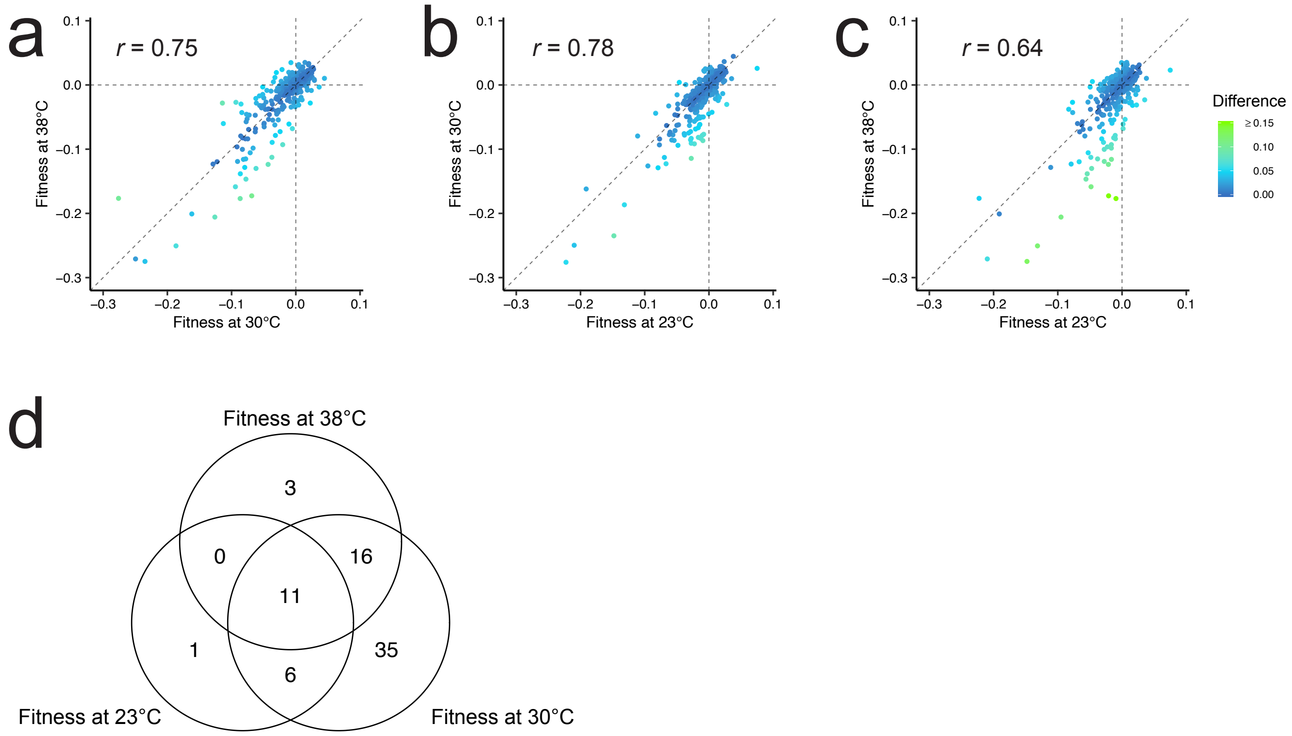


**Suppl. Fig. S14. Fitness at different temperatures is largely modulated by a common set of genes.** Generation-normalized gene log2FCs (n=290) of cellular fitness screens at 23, 30 and 38°C compared to each other (a, b, c). Spearman correlations are shown. d) Overlap of genes with significant fold change between fitness screens.

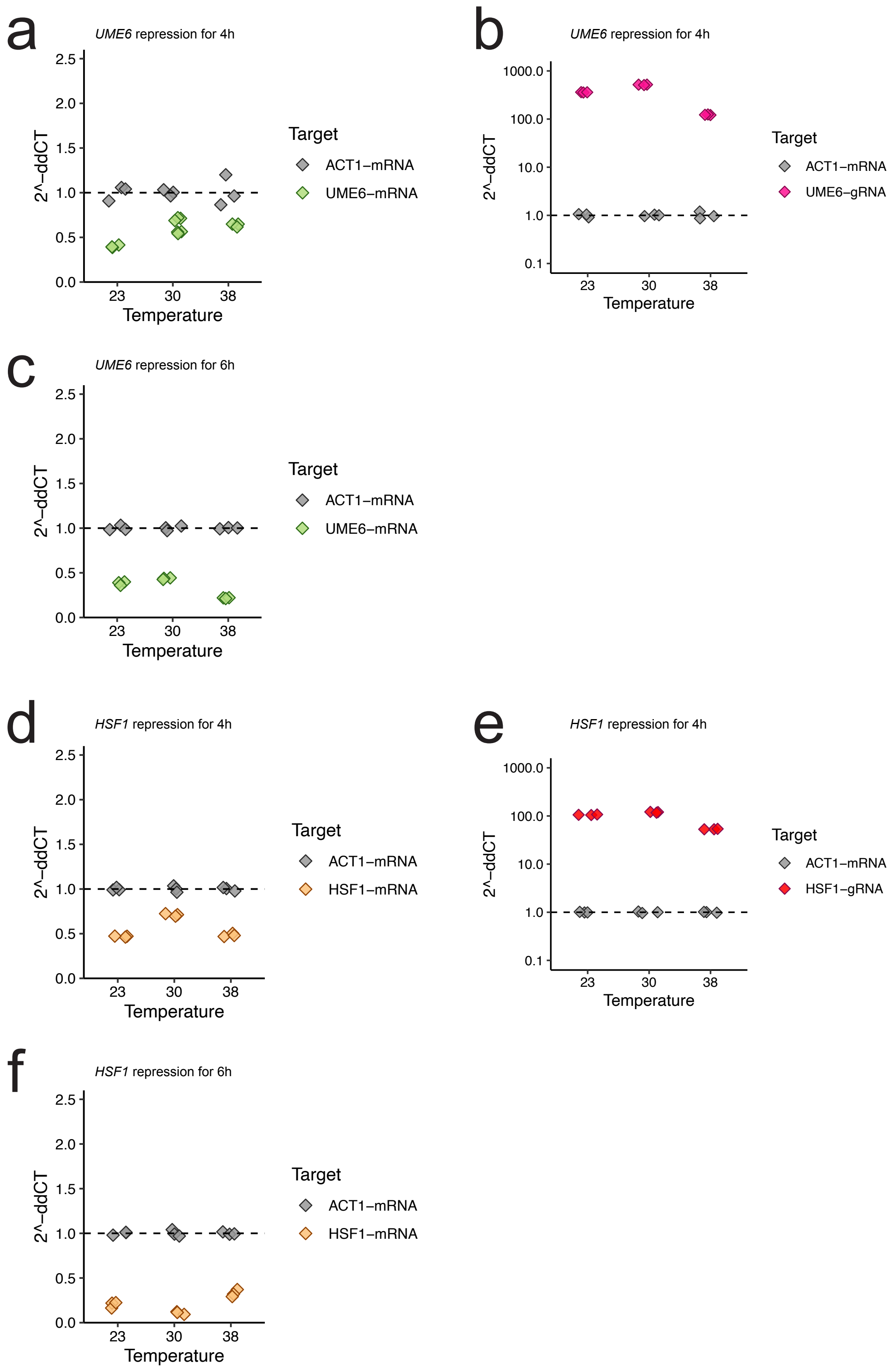


**Suppl. Fig. S15. CRISPRi efficacy across temperatures and induction time.**
Quantitative reverse-transcription PCR was performed to quantify mRNA and gRNA levels in induced and non-induced *UME6* and *HSF1* CRISPRi strains grown at temperatures 23, 30 and 38°C. *ACT1* was used as reference to determine dCT values, and ddCT values were calculated by subtracting -ATc from +ATc dCT values (see Methods). a) *UME6* mRNA after 4h of *UME6* repression, b) *UME6* gRNA after 4h of *UME6* repression, c) *UME6* mRNA after 6h of UME6 repression, d) *HSF1* mRNA after 4h of *HSF1* repression, e) *HSF1* gRNA after 4h of *HSF1* repression, f) *HSF1* mRNA after 6h of *HSF1* repression. Measurements are performed in n=3 (or n=6 in case of *UME6* mRNA 4h time point) technical replicates based on individual measurements but from the same mRNA sample.

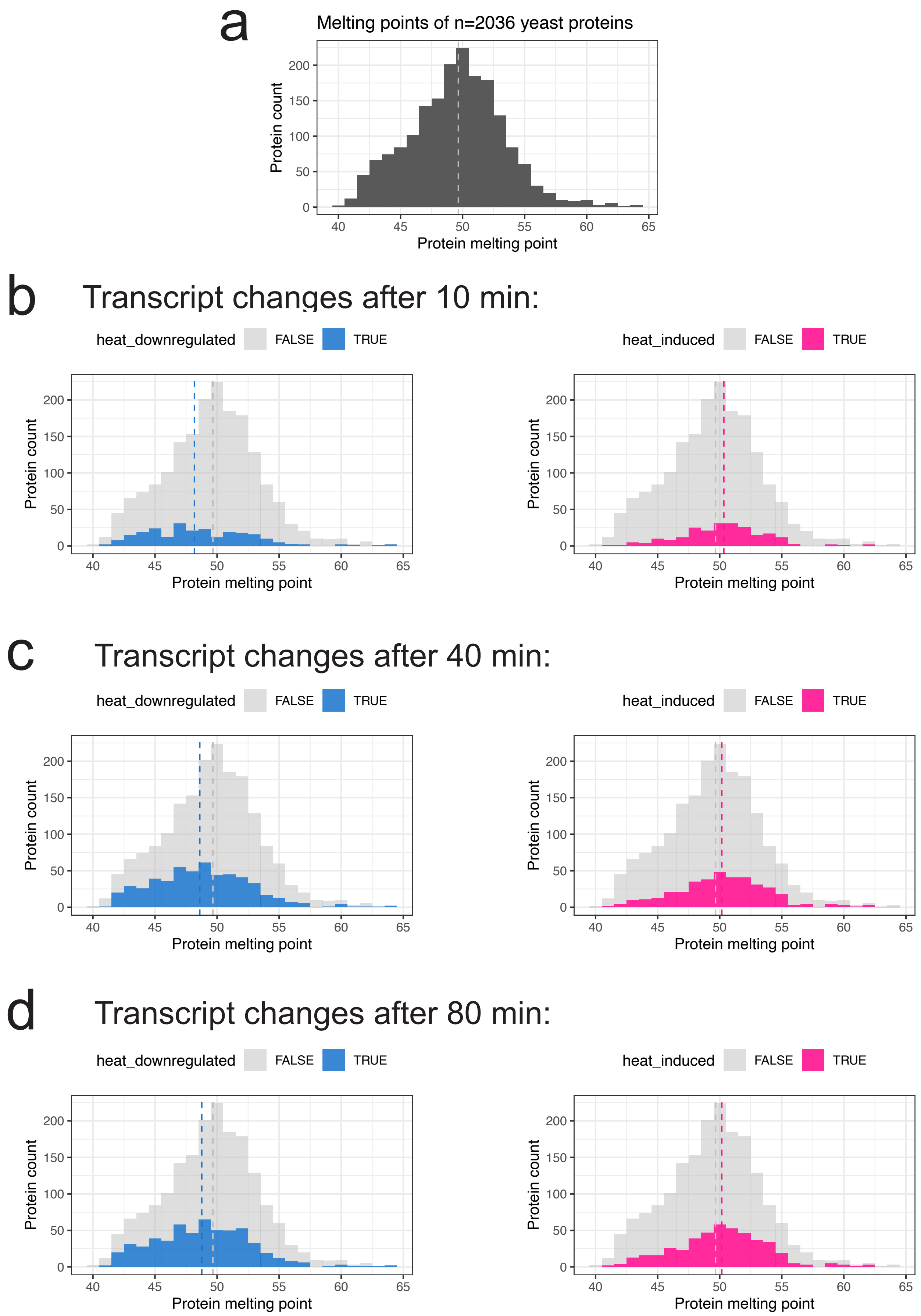


**Suppl. Fig. S16. Protein melting points of genes induced or down-regulated upon heat shock.**
a) Distribution of protein melting points obtained from thermal protein profiling (measured in yeast cell lysate; Jarzab et al., 2020) for proteins with transcripts quantified in microarray data on heat shock expression (Mühlhofer et al., 2019). Melting point distributions are shown for genes (n=2036) that are down- (left panels) or upregulated (right panels) upon a shift from 25 to 42°C with 10 min (b), 40 min (c) or 80 min (d) exposure.

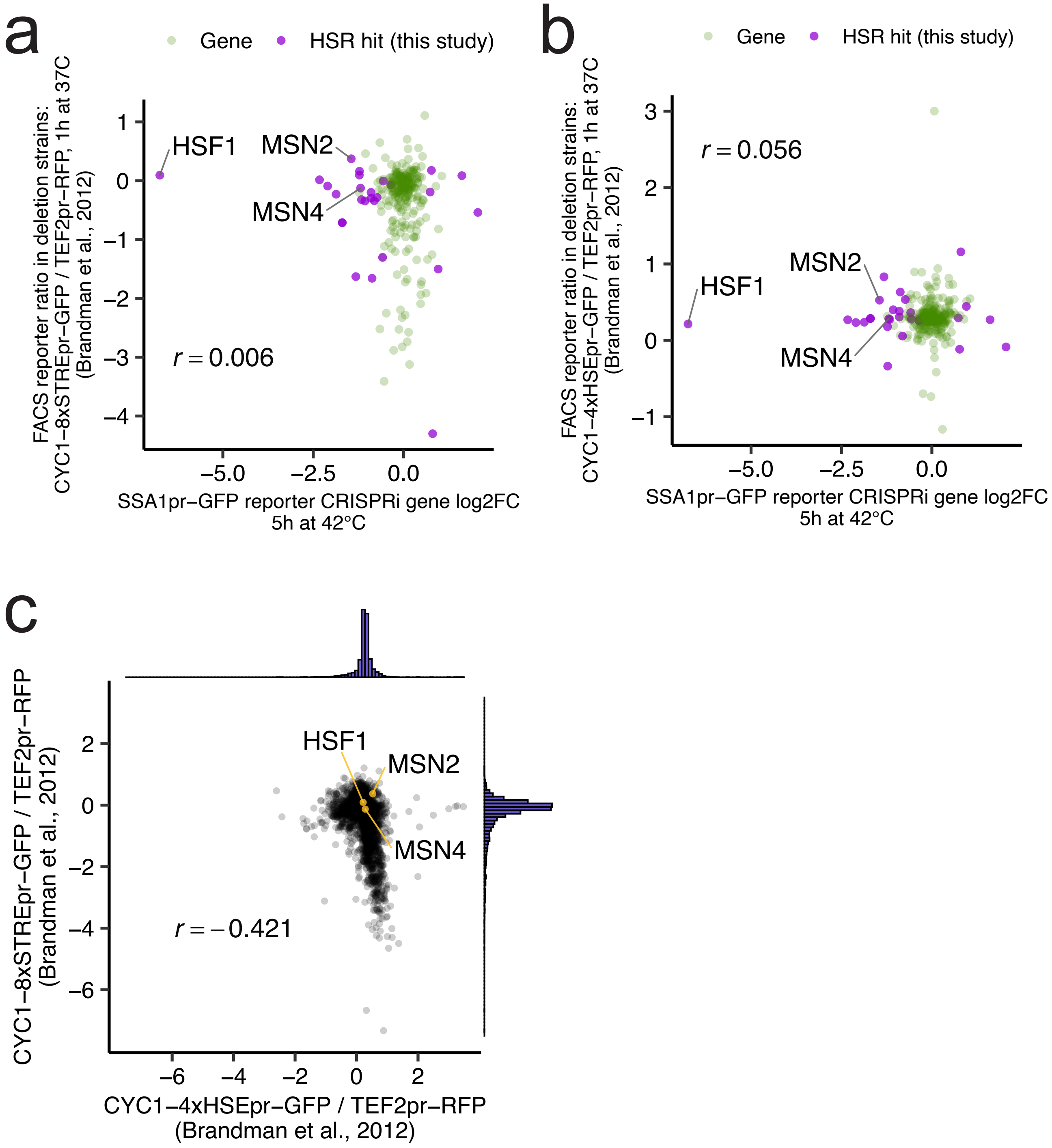


**Suppl. Fig. S17. Regulation of the SSA1 Hsp70 promoter are incompletely explained by artificial HSE or STRE reporters.**
CRISPRi screen HSR gene log2FCs are compared to deletion strain signals of a) stress response element or b) heat shock element reporters (n=274) from Brandman et al. (2012). The CRISPRi screen log2FCs are based on FACS-enrichment of cells depending on expression of the Hsp70 reporter gene (*SSA1pr-GFP*), after 5h incubation at 42°C. The deletion strain HSE and STRE scores are the ratio between FACS fluorescence measurements of green to red fluorescent protein (*GFP/RFP*), where *GFP* is expressed by a crippled 225 nucleotide CYC1 promoter (Guarente and Mason, 1983) combined with either a 4x Heat Shock Element (HSE) repeat sequence (Sorger and Pelham, 1987) or a 8x STress Response Element (STRE) repeat sequence (Marchler et al., 1993), and *RFP* expression is controlled by the yeast *TEF2* promoter, and measurements were taken after 1h incubation at 37°C (Brandman et al., 2012). c) Comparison of HSE and STRE reporter signals (n=5977) using data from Brandman et al. (2012). Marginal histograms display distribution of data points on axes. Genes encoding the regulators of HSE (*HSF1*) and STRE expression (*MSN2* and *MSN4*) are labelled.

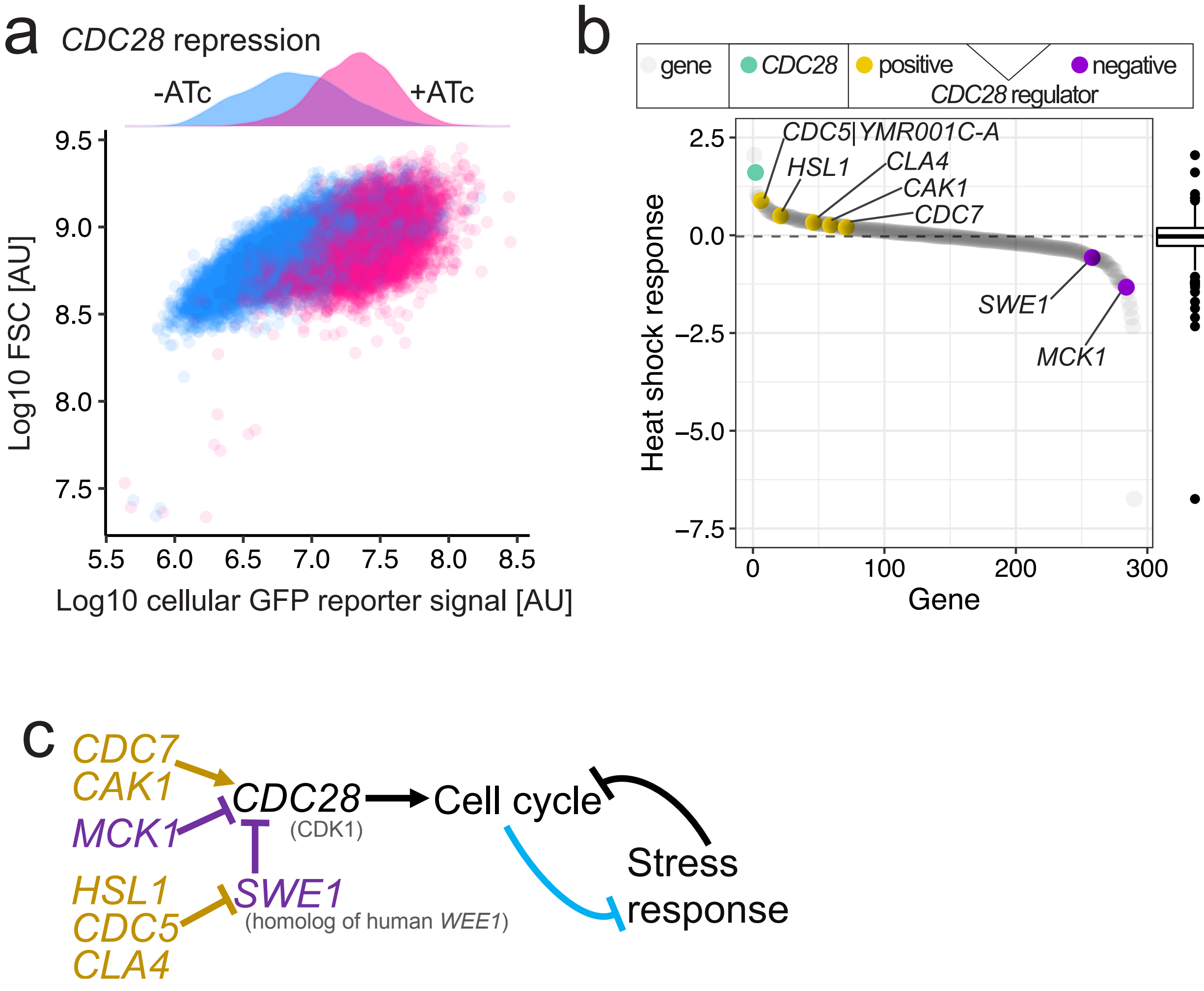


**Suppl. Fig. S18. Repression of CDC28 suggests HSR-antagonizing function.**
a) Cellular GFP intensity (x-axis) versus forward scattering (y-axis) of *CDC28* repression strains after 5h at 42°C measured with FACS, and down-sampled to n=20000 data points. Density distributions of the GFP signal are shown on top. b) Distribution of HSR screen gene fold changes with *CDC28* and its positive and negative regulators marked in yellow and purple, respectively. The boxplot on the right denotes median, quantiles and outliers for all PKs and TFs (n=290). c) Known molecular interactions regulating *CDC28* activity with stimulators and inhibitors colored purple and yellow, repectively. Heat exposure and other stresses are known to result in a cell cycle arrest. The repressive role of *CDC28* on HSR activity suggests that, vice-versa, the cell cycle also inhibits the stress response, marked blue.

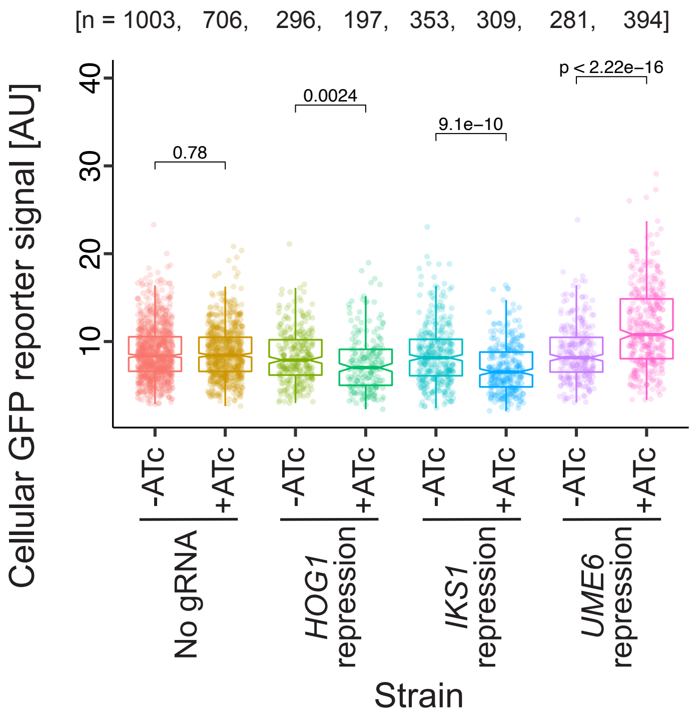


**Suppl. Fig. S19. Modulating Hsp70 expression via CRISPRi**
Cellular SSA1pr-GFP reporter intensity (y-axis) of individual CRISPRa strains cultured with or without ATc (x-axis) after 5h exposure to 42°C, and imaged with fluorescence microscopy. Dots denote single cells. Used gRNAs are Hog1_g2, Iks1_g5, Ume6_g5. Two-sided Wilcoxon adjusted p-values are given for tests between samples.

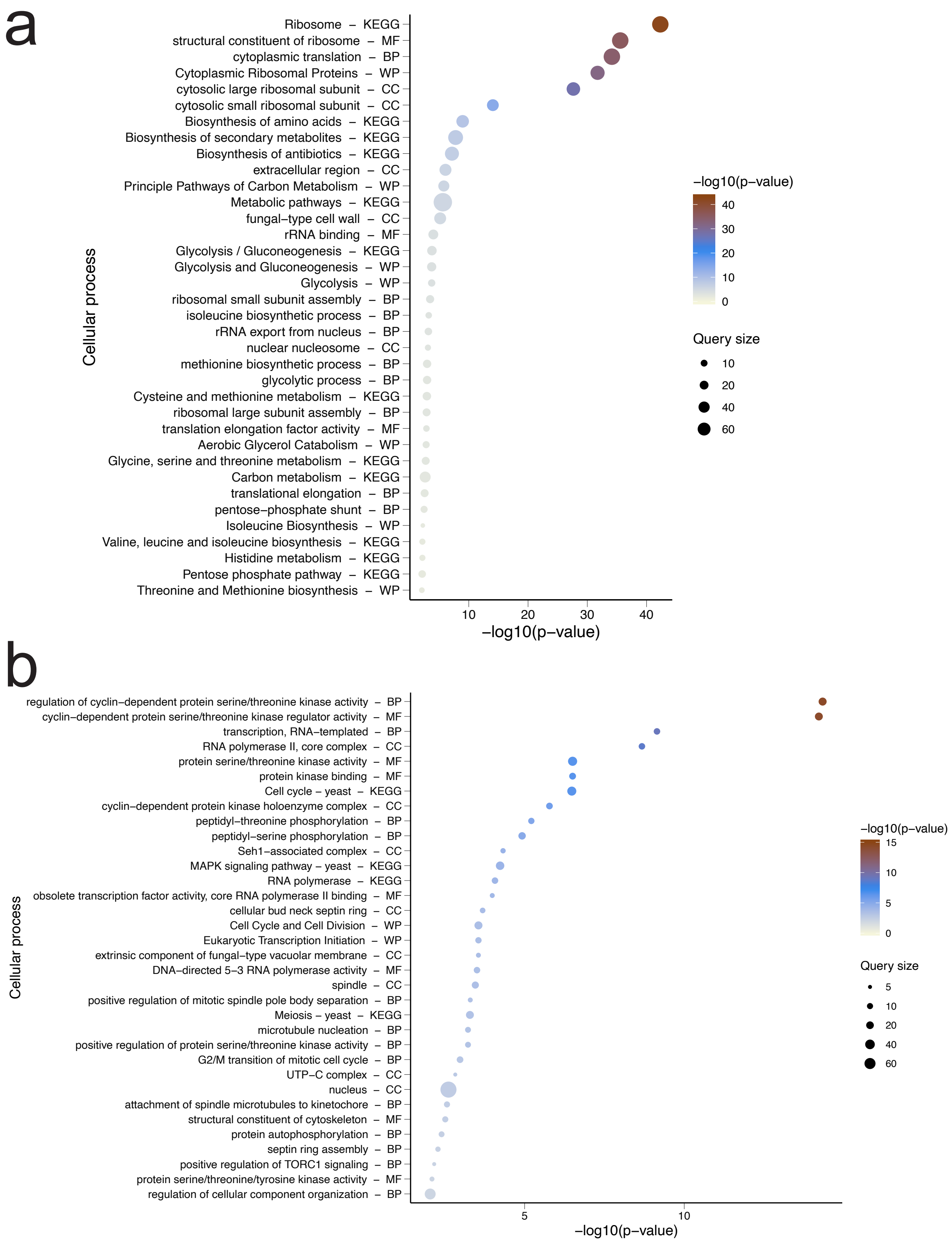


**Suppl. Fig. S20. Target GO-Enrichments. [a & b from a-h]**

Gene Ontology (GO) enrichments for high confidence TF target genes inferred from chromatin-immunoprecipitation data (Gonçalves et al., 2017) and PK phosphorylation targets derived from phospho-proteomics data (Sadowski et al., 2013) of identified modulators, generated with the gProfiler 2 R package (Kolberg et al., 2019). Additional annotations mark entichments based on cellular components (CC), biological processes (BP), molecular functions (MF), as well as Wiki-Pathways (WP) and KEGG-derived pathways (KEGG). Query size and -log10 p-values are indicated by dot size and color gradient, respectively.

a) GO enrichment for target genes (n=664) of TFs (n=33) which modulate cellular fitness at 30°C.

b) GO enrichment for interactors (n = 297) of PKs (n=35) which modulate cellular fitness at 30°C.

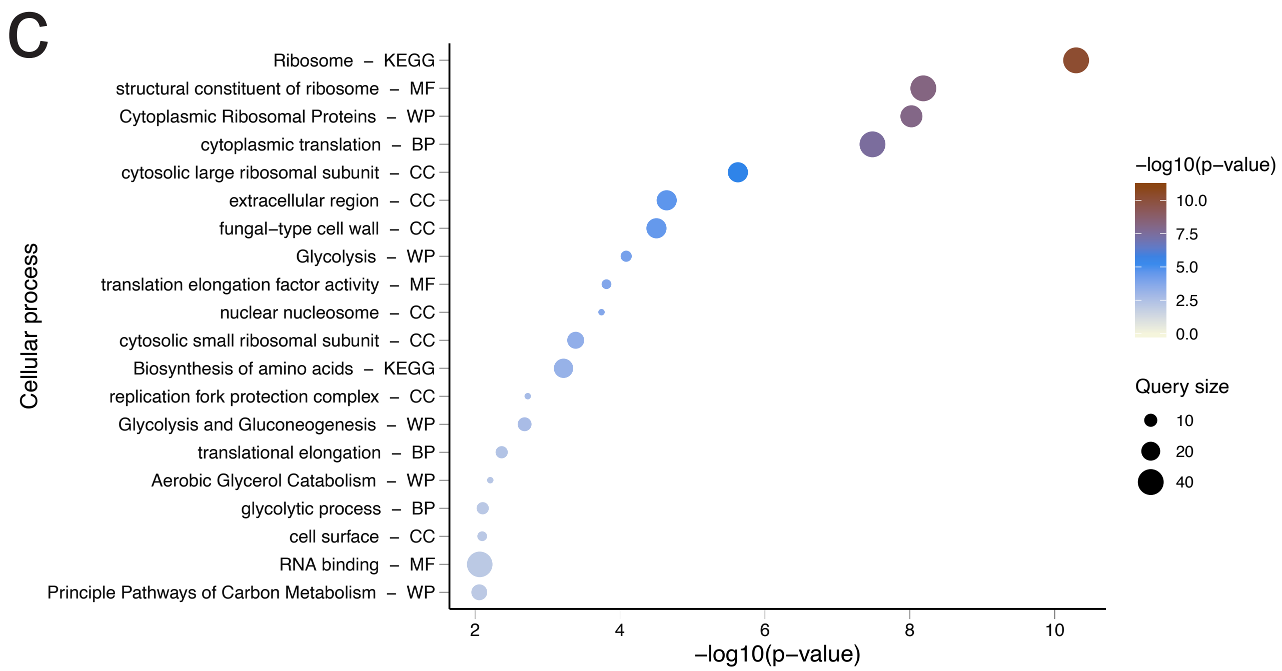


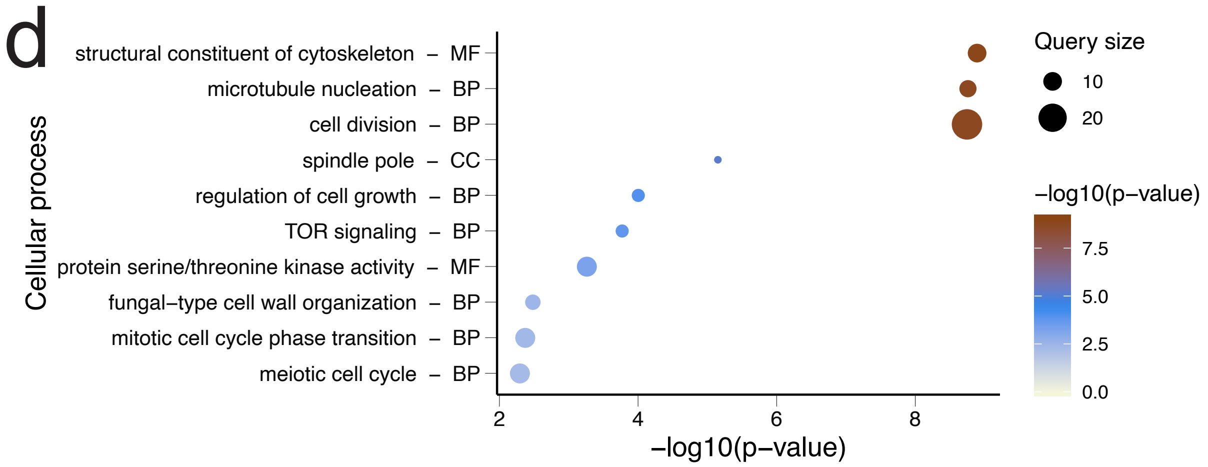


**Suppl. Fig. S20. Target GO-Enrichments. [continued for c & d from a-h]**
c) GO enrichment for target genes (n=207) of TFs (n=11) which modulate cellular fitness specifically at 38°C.
d) GO enrichment for interactors (n=56) of PKs (n=4) which modulate cellular fitness specifically at 38°C.

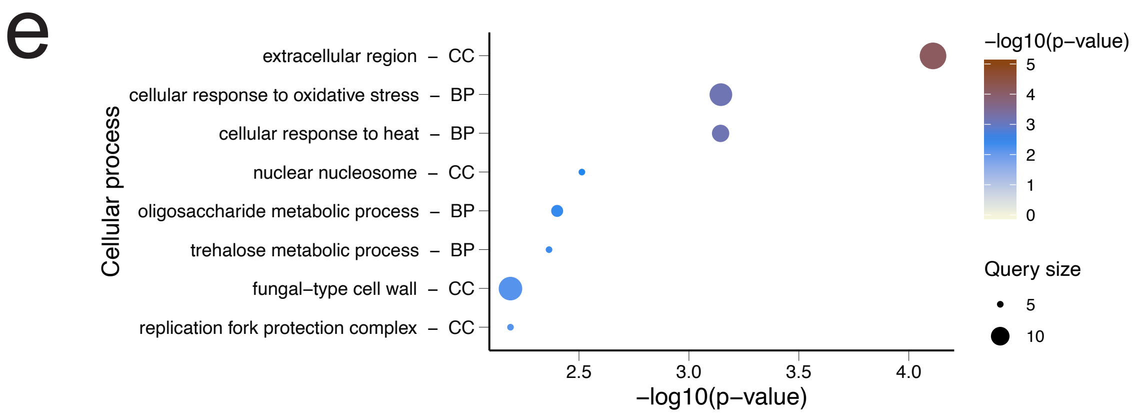


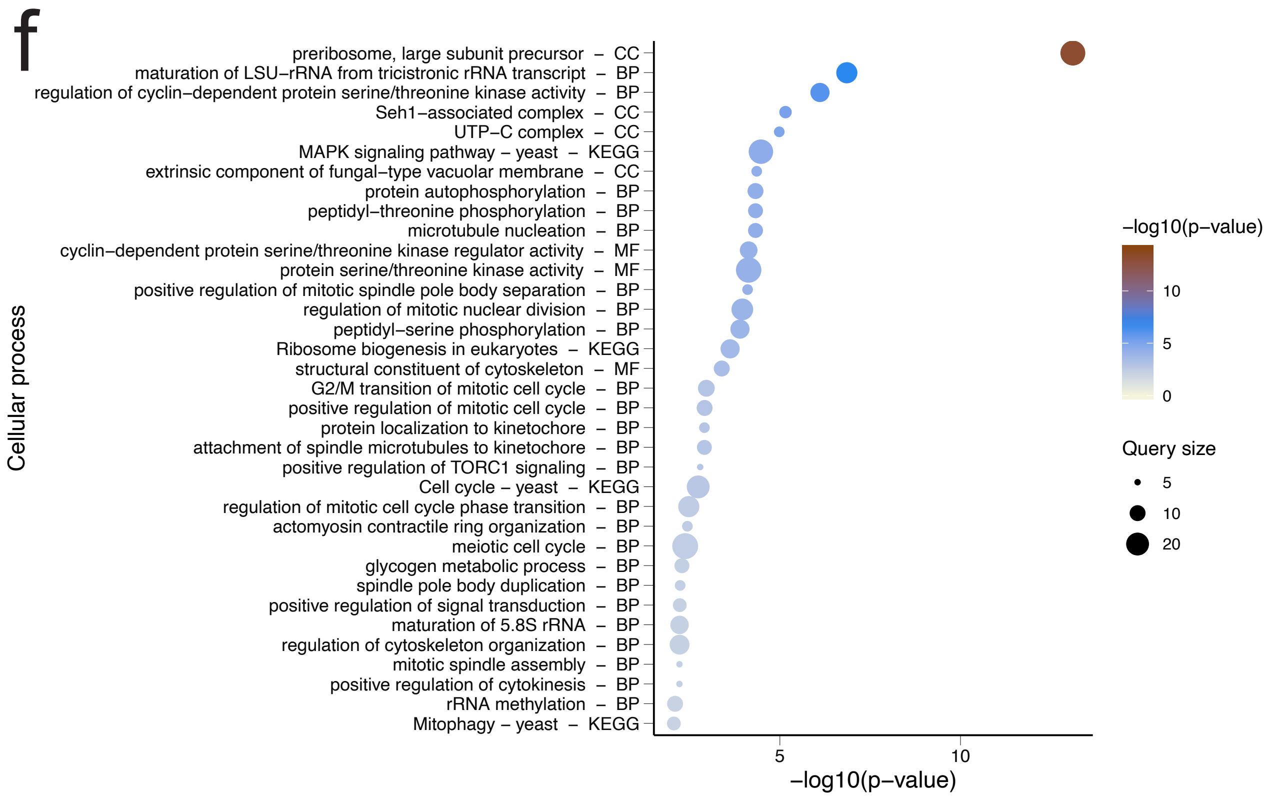


**Suppl. Fig. S20. Target GO-Enrichments. [continued for e & f from a-h]**
e) GO enrichment for target genes (n=141) of TFs (n=10) which modulate the HSR.
f) GO enrichment for interactors (n=216) of PKs (n=17) which modulate the HSR.

g

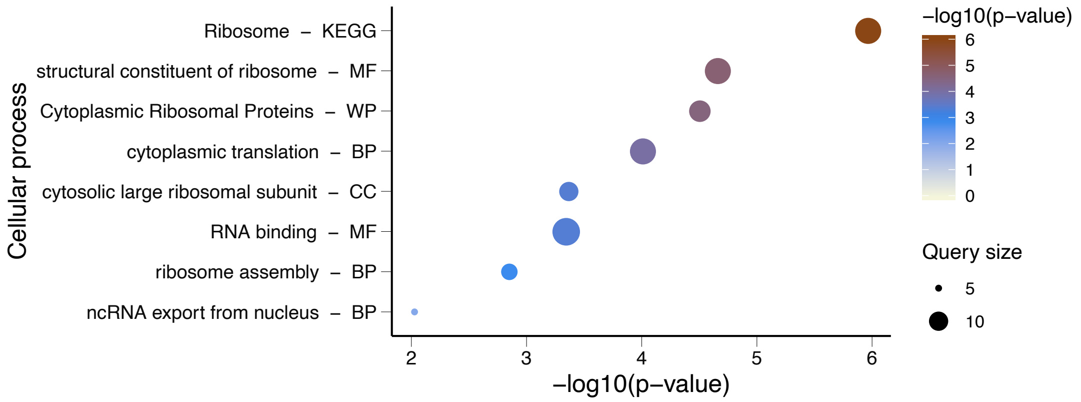


h

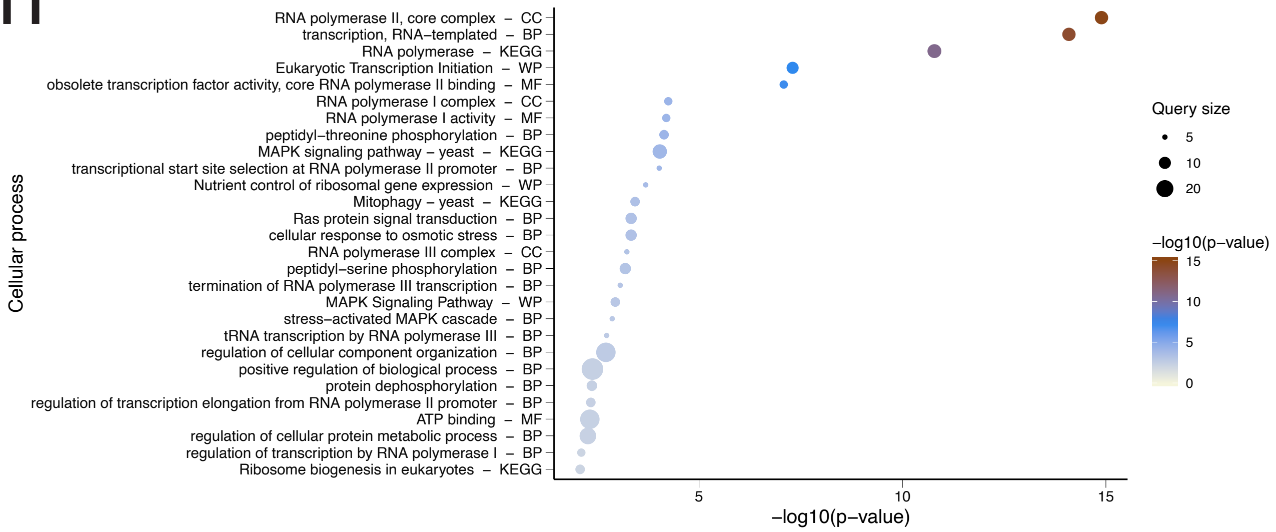


**Suppl. Fig. S20. Target GO-Enrichments. [continued for g & h from a-h]**
g) GO enrichment for target genes (n=62) of TFs (n=5) which modulate thermotolerance.
h) GO enrichment for interactors (n=99) of PKs (n=10) which modulate thermotolerance.

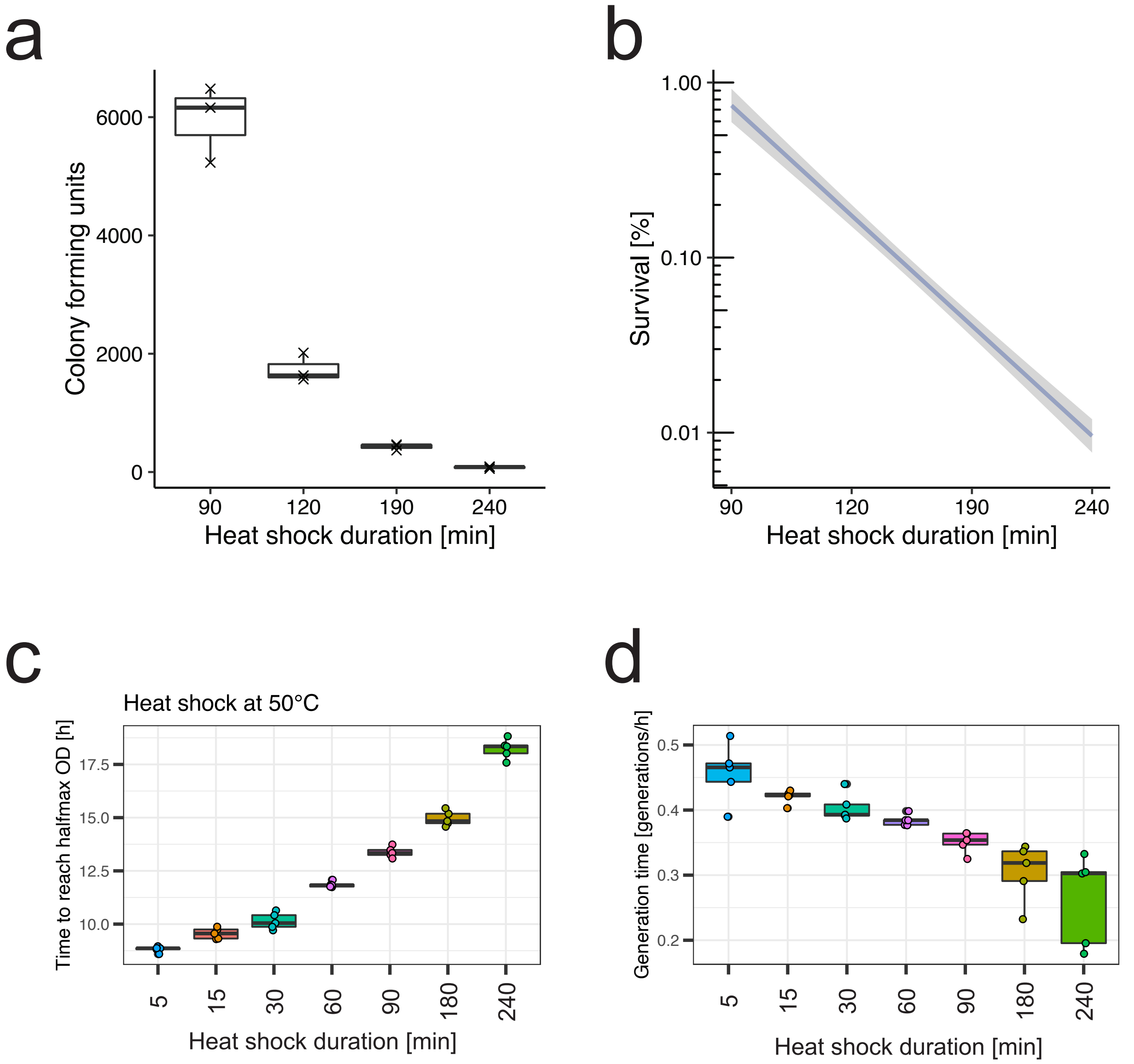


**Suppl. Fig. S21. Heat shock effects on survival and resumed growth.**
The BY4743 SSA1pr-GFP strain was cultured at 30°C, diluted to OD600=0.3 in YPD medium and incubated for different time intervals (x-axis) at 50°C with n=3 biological replicates. After exposure, 100µl cell suspension was spread on YPD plates. After 3 days incubation at 30°C, colony-forming units were counted (a) and survival rates determined (b). The smooth fit was generated with the ggplot2 geom_smooth function with method="lm" and default settings for span. An exposure time of 150 min results in less than 0.1 % surviving cells, which was chosen as screen condition.
For populations that have been exposed to a 50°C for different time intervals, the time to reach half-maximum OD600 (c) and the generation time (d) has been calculated from n=5 replicate wells by OD600 measurements in a plate reader.

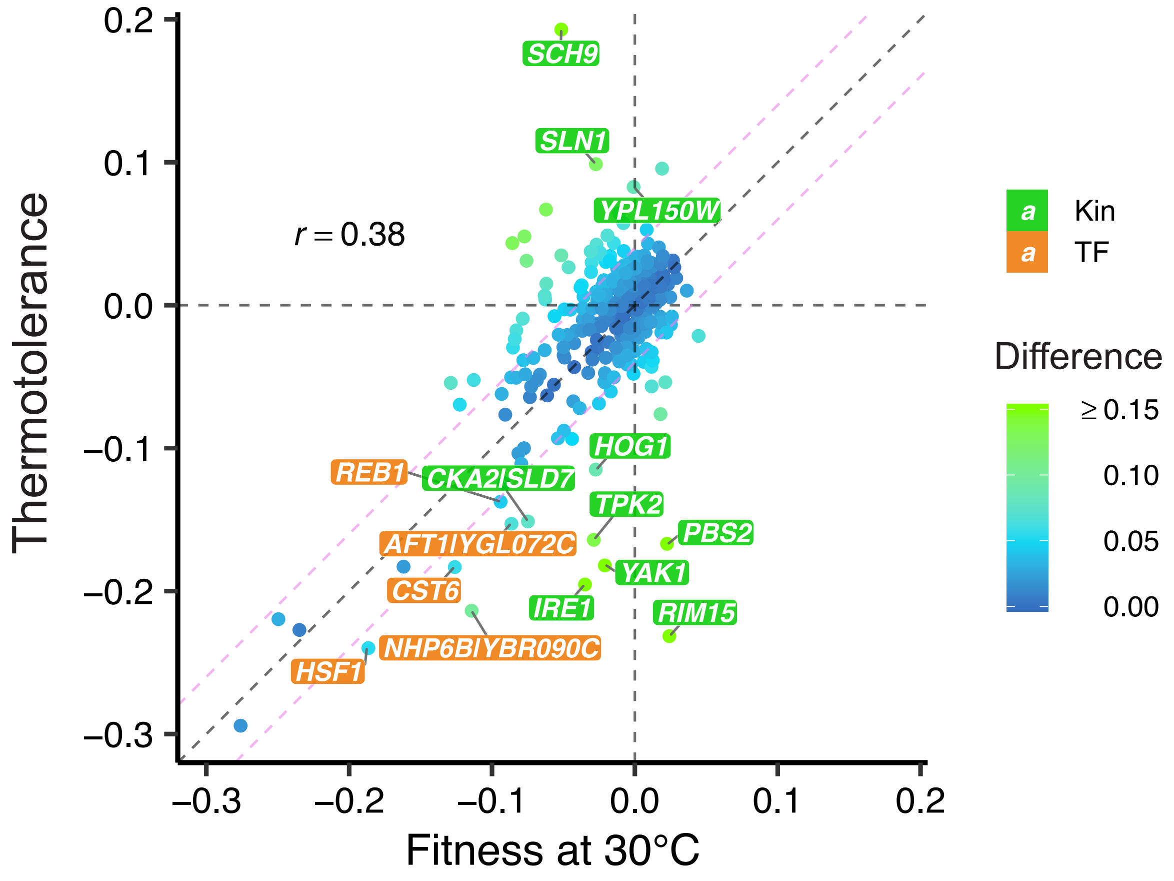


**Suppl. Fig. S22. Thermotolerance effects can be readily separated from fitness effects.**
Comparison of generation-normalized gene log2FCs (n=290) between screens for fitness at 30°C and thermotolerance (Survival of a sudden exposure to 50°C for 150 min and recovery). Dots denote genes colored by difference in normalized fold change. Dashed grey lines are intercepts marking a normalized fold change of 0 and the diagonal. Dashed violet lines mark difference thresholds of 0.05 and -0.05. Genes with specific function in thermotolerance which exceed the difference threshold (indicating that thermotolerance effects are stronger than fitness effects) are labelled on orange (TFs) or green background (PKs).

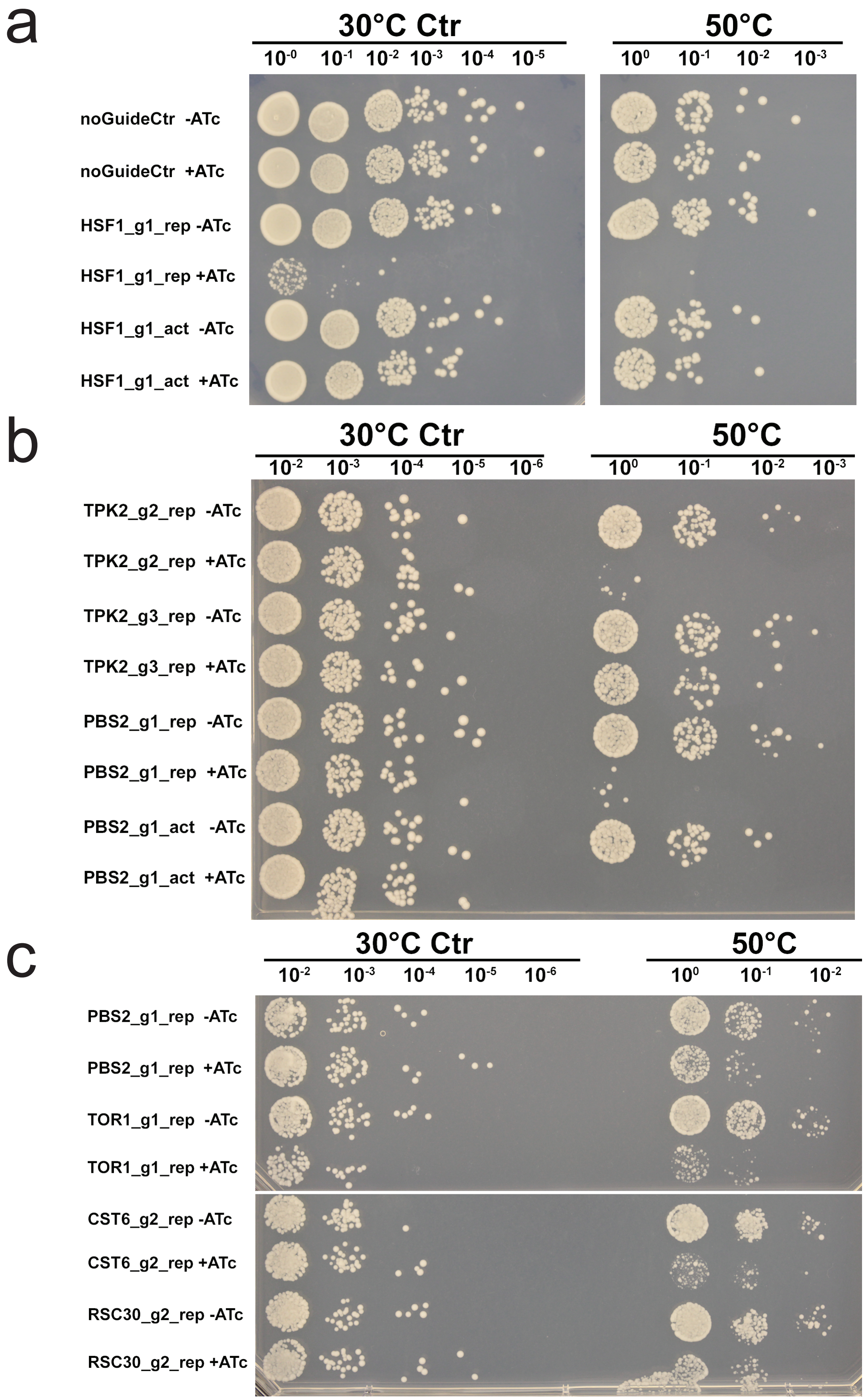


**Suppl. Fig. S23. Dilution spot plating thermotolerance assay.**
a) and b) are dilution spot plating of cultures exposed to 50°C for 150 min, as applied in the thermotolerance screen, or remained at 30°C as reference. The noGuideCtr strain has a non-functional gRNA. Dilutions and treatment are indicated on top. Targeted genes, used guide RNAs (g1-3), repression or activation strain (rep/act) and ATc conditions are indicated on the left. Experiments were performed in triplicate and exemplary results are shown. c) Dilution spot plating as in (a) and (b), but with cultures exposed to 50°C for 90 min.

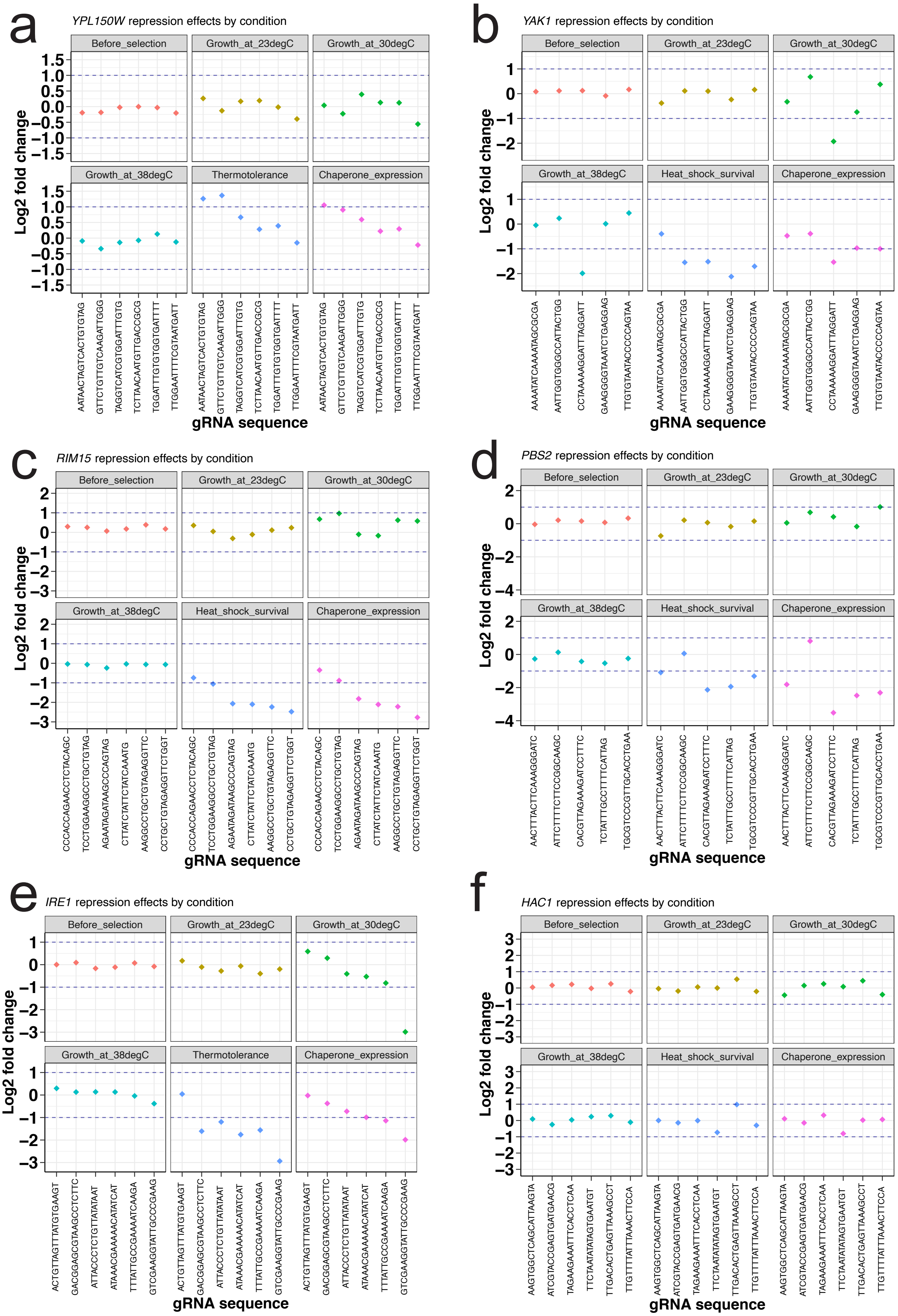


**Suppl. Fig. S24. Detailed guide fold changes for selected genes across screens.**
Log2FCs (y-axis) of gRNAs (x-axis) are shown across all screens for repression of *YPL150W* (a), *YAK1* (b), *RIM15* (c), *PBS2* (d), *IRE1* (e) and *HAC1* (f). Dashed blue lines denote log2FC cutoffs 1 and -1.

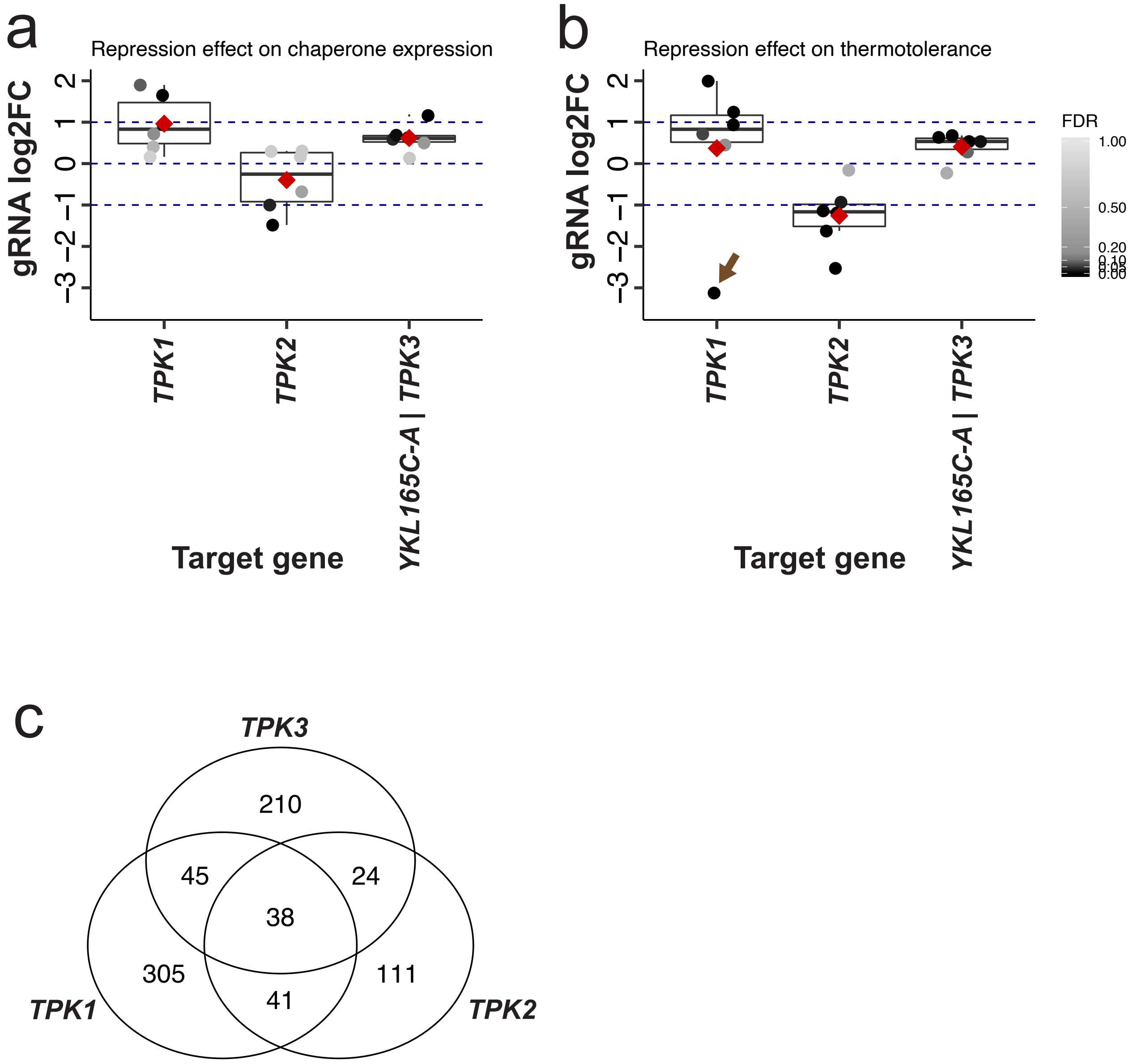


**Suppl. Fig. S25. Catalytic protein kinase A subunits differentially regulate HSR and thermotolerance.** a) HSR screen gRNA log2FCs for *TPK1*, *TPK2* and the bidirectionally regulated *TPK3*|*YKL165C-A* loci. Dots denote gRNA barcodes and are colored by FDR. Boxplots denote median and quantiles of gRNA log2FCs. Gene fold changes are added as red diamonds, representing mean gRNA log2FCs. b) Thermotolerance screen gRNA log2FCs for genes as shown in a). Brown arrows in b) point to a GC-rich gRNA (GCGCTATTAG-GGGGGAGGGA) which likely caused fitness effects through an off-target thus masking increased thermotolerance effects. c) Overlap of protein-protein interactions between catalytic PKA subunits based on phospho-proteomics data from the phosphogrid database 2.0 (Sadowski et al., 2013).

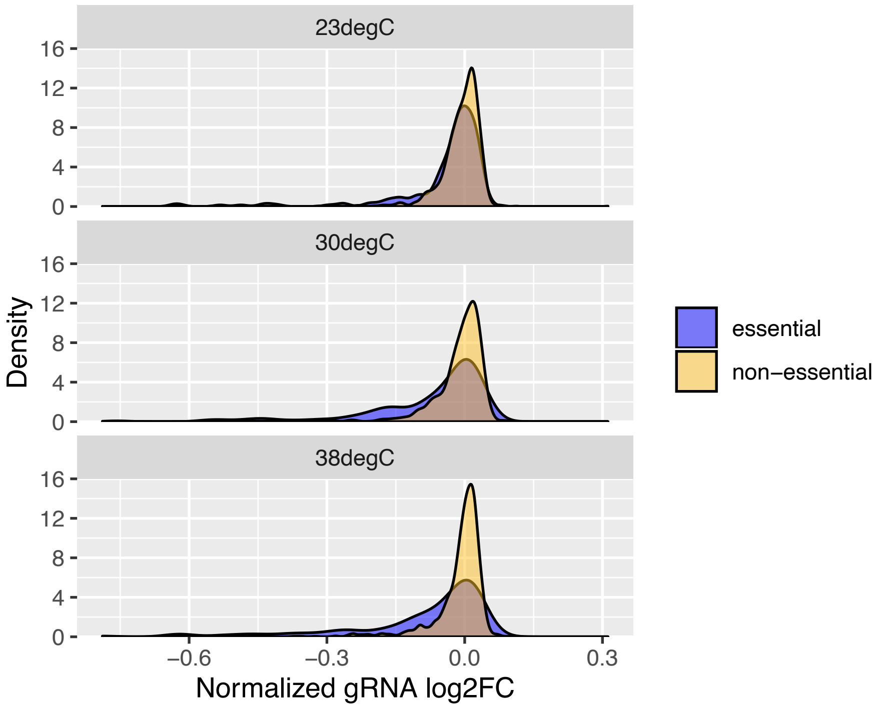


**Suppl. Fig. S26. Temperature-dependency of gRNA barcode dropout**
Generation-normalized guide log2FC density (y-axis) of fitness screens at 23, 30 and 38°C are shown for essential and non-essential genes as defined by non-viable knock-out annotations according to SGD (Cherry et al., 1998) for n=1526 gRNA barcodes quantified across screens.

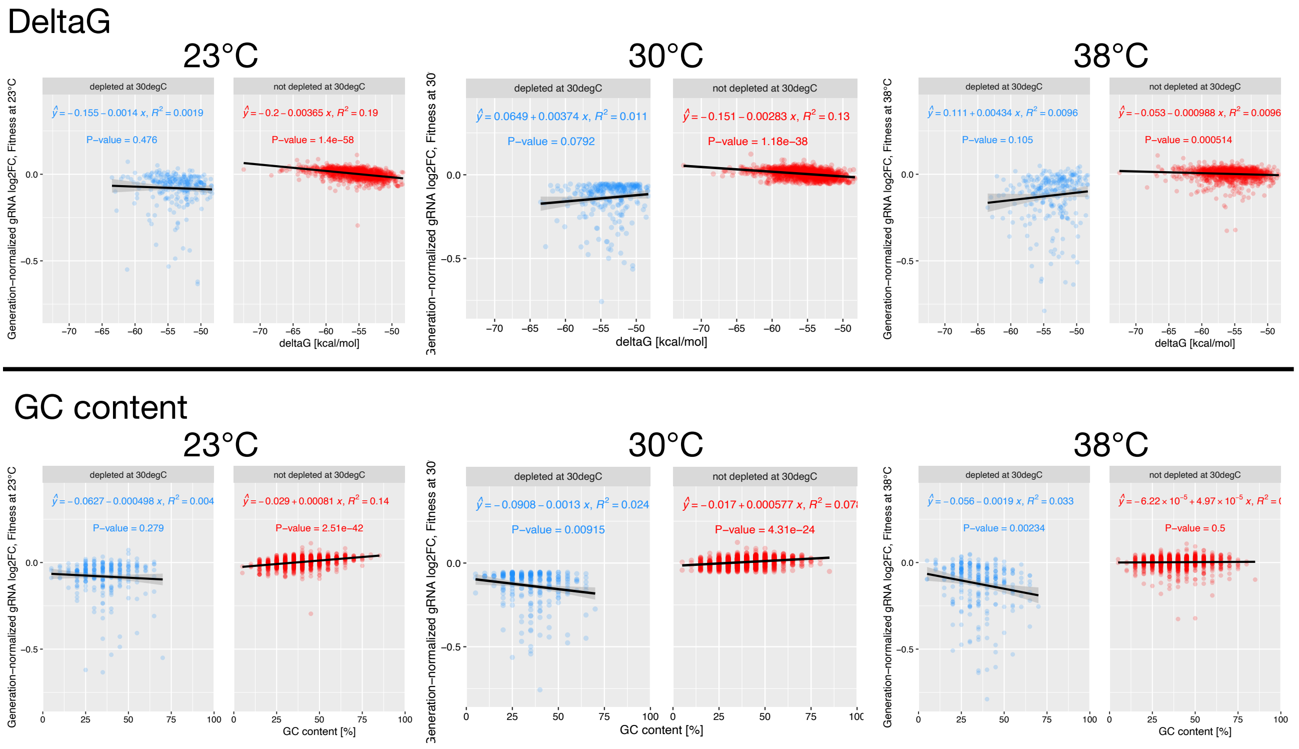


**Suppl. Fig. S27. The gRNA sequence contributes to temperature-dependent CRISPRi efficiency.**
Normalized gRNA log2FCs (y-axis) of fitness screens at 23, 30 and 38°C are shown depending on the deltaG Gibbs free energy of their most probable secondary structure (determined with Vienna RNAfold) in the top panel or their GC content in the bottom panel, facetted for depleted and non-depleted gRNAs in blue and red, respectively. Dots denote gRNAs quantified across screens (n=1526).

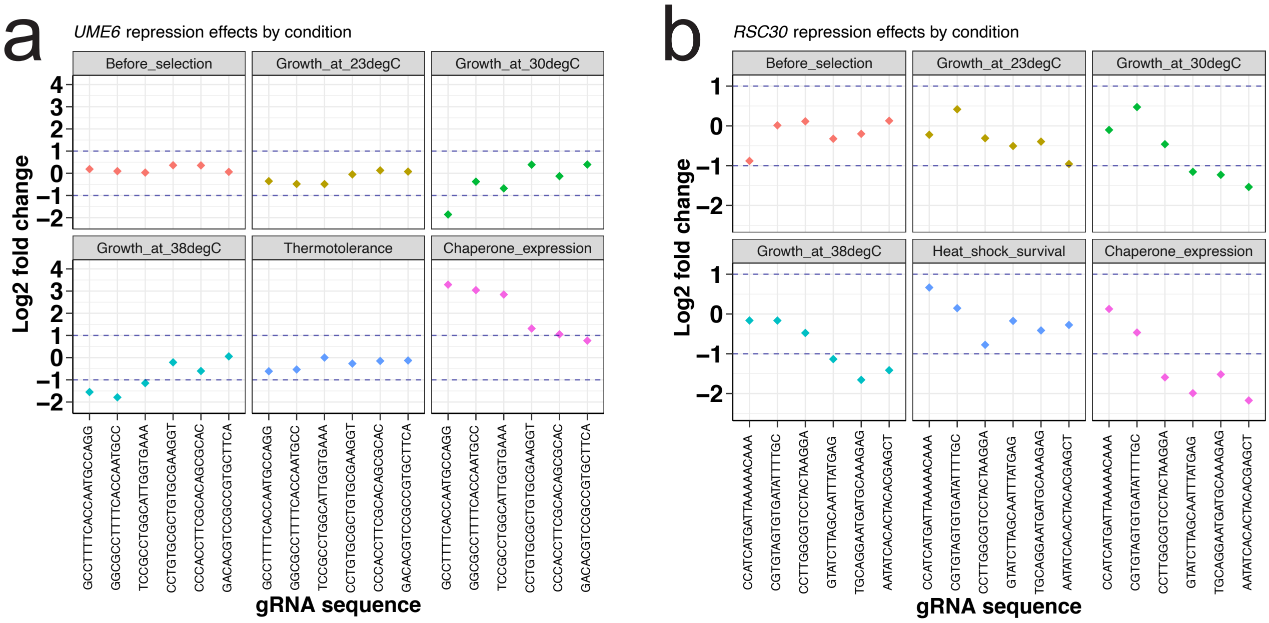


**Suppl. Fig. S28. Detailed guide fold changes for *UME6* and *RSC30* across screens.** Log2FCs (y-axis) of gRNAs (x-axis) are shown across all screens for repression of a) *UME6* and b) *RSC30*. Dashed blue lines denote log2FC cutoffs 1 and -1.

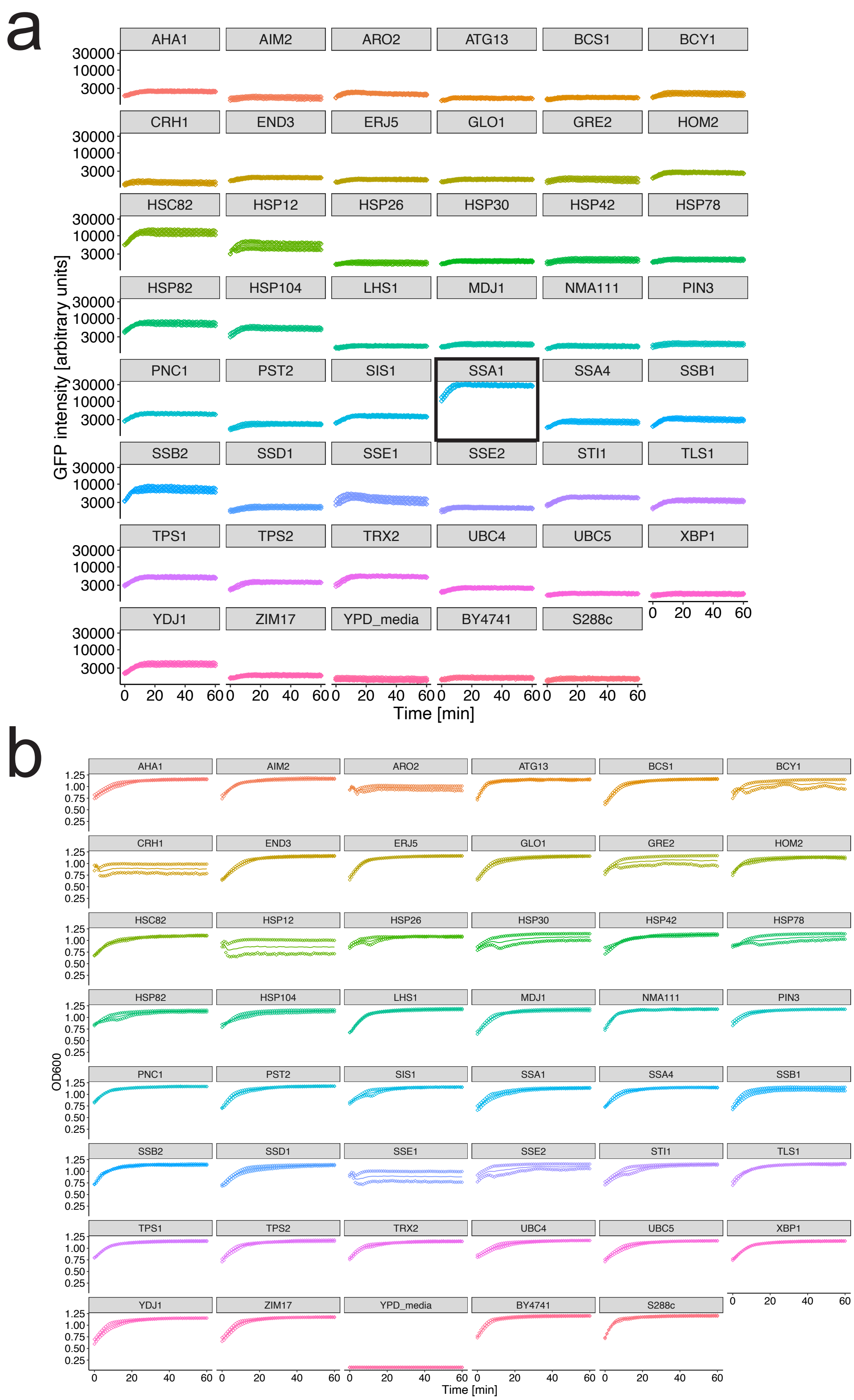


**Suppl. Fig. S29. GFP collection strain induction upon heat shock.**
In order to identify a suitable reporter gene for HSR activity, 44 strains of the yeast GFP-tag collection (Hu et al., 2003) were profiled for their (a) GFP intensity and in parallel quantified (b) OD600 over time upon shift from 30 to 40°C at time 0. Diamonds denote measurements of duplicate wells (n=2) and the lines denotes their mean.

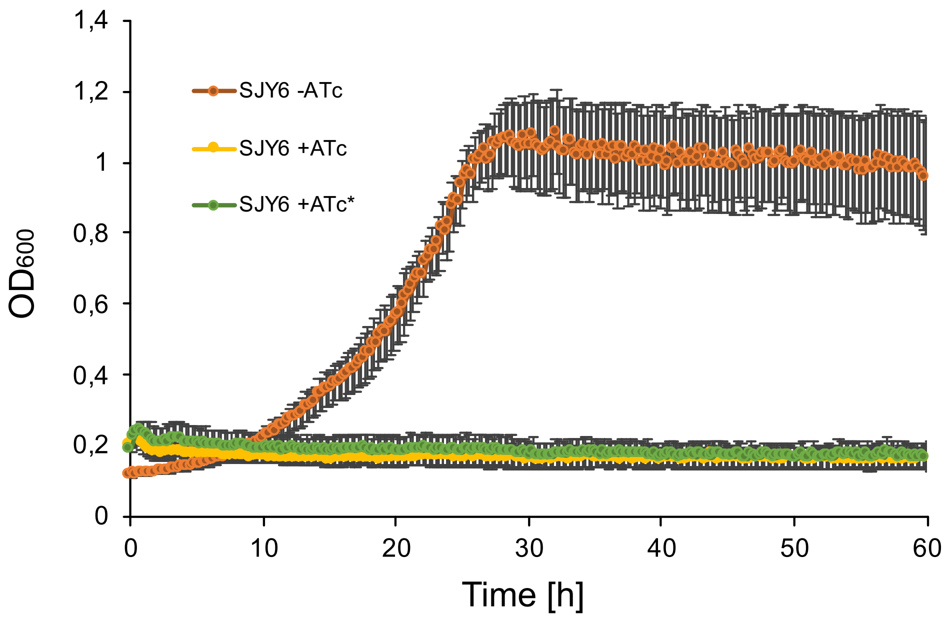


**Suppl. Fig. S30. ATc maintains biological activity at 38°C.**
Growth curves depict OD600 (y-axis) over time (x-axis) for cultures of the *S. cerevisiae* SJY6 strain, grown at 30 °C in different conditions: in presence of 250 ng/ml Anhydrotetracycline (+ATc), in presence of 250 ng/ml ATc that has been incubated for two days at 38°C (+ATc*), and without addition of ATc as reference. Dots denote averaged OD600 values of n=12 replicate well cultures with standard deviations shown as error bars. *In SJY6, transcription of the essential SPT16 gene is inhibited by Anhydrotetracycline (ATc) due to an upstream TetO regulatory system (TetOperonOFF::SPT16) (Jimeno Gonzales et al., 2006).*

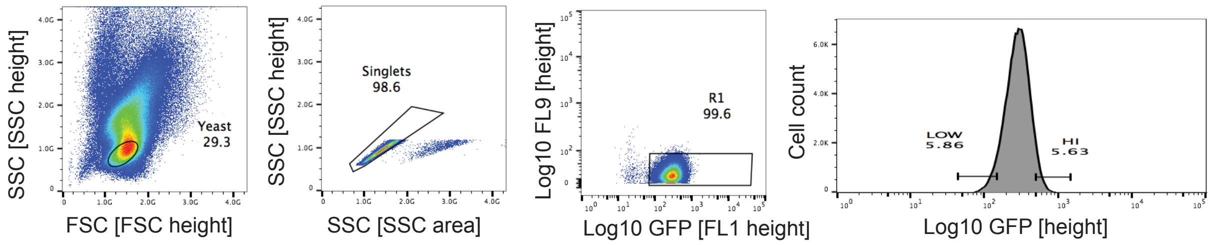


**Suppl. Fig. S31. FACS gating scheme.**
Gating was used to select cells representing the bulk population regarding the forward and sideward scatter profiles (cells with similar size and texture), to exclude dividing cells, to select for alive GFP-positive cell, and to collect 250.000 – 500.000 cells within the top and bottom 5% of cellular GFP reporter intensity.

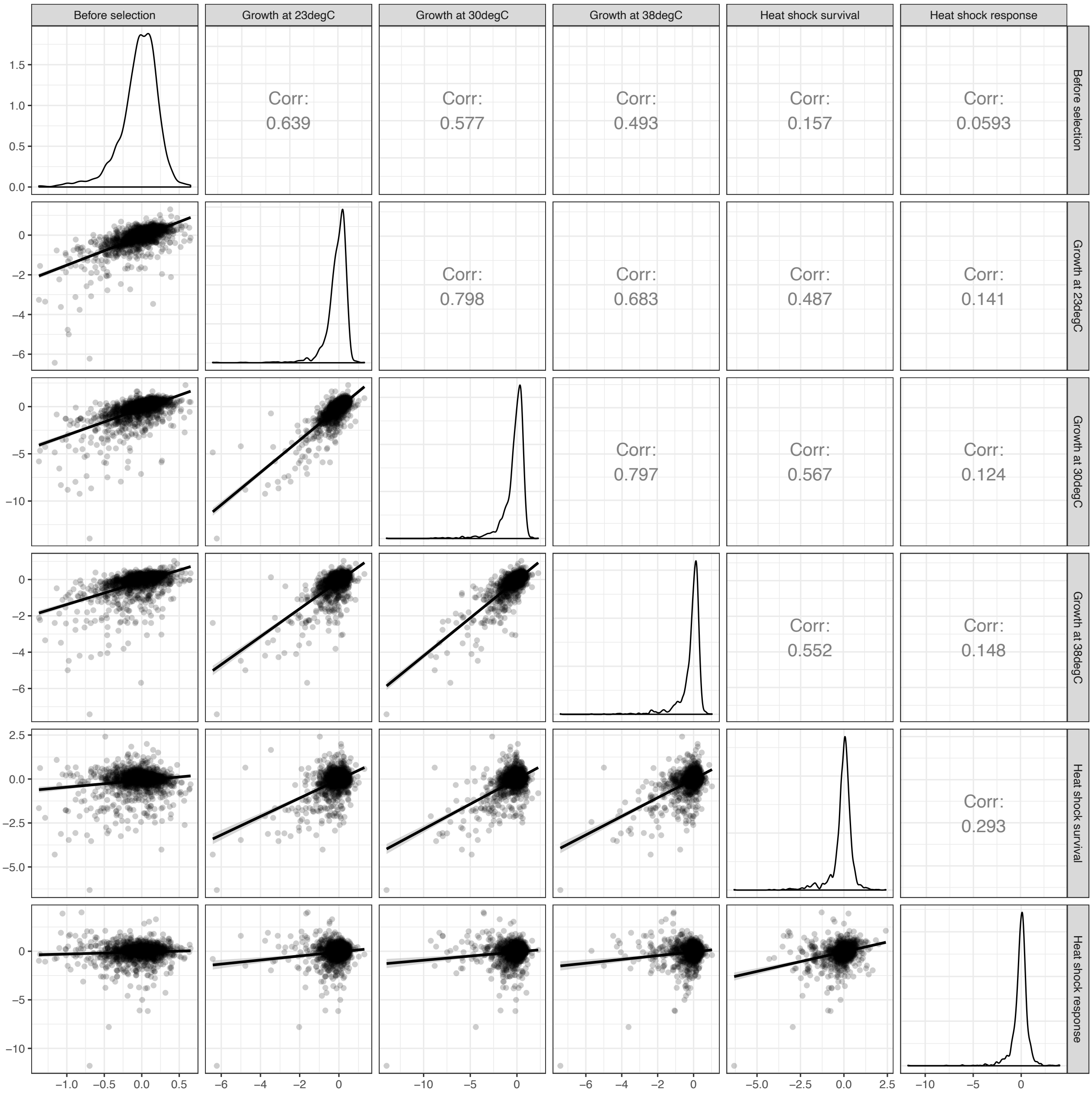


**Suppl. Fig. S32. Guide RNA barcode log2FCs across screens.**
Guide RNA log2FCs are compared between screens as indicated in the panel strips, merged for TF and PK libraries. Dots denote gRNA barcodes quantified across screens (n=1526). The graph was generated with the ggpairs function of the ggally R package (Schloerke et al., 2020) and displays Pearson correlations. Linear model for smoothed line is generated in R ggplot using geom_smooth with method="lm" and default settings. Please note that fitness screens are not normalized for generation number during selection.

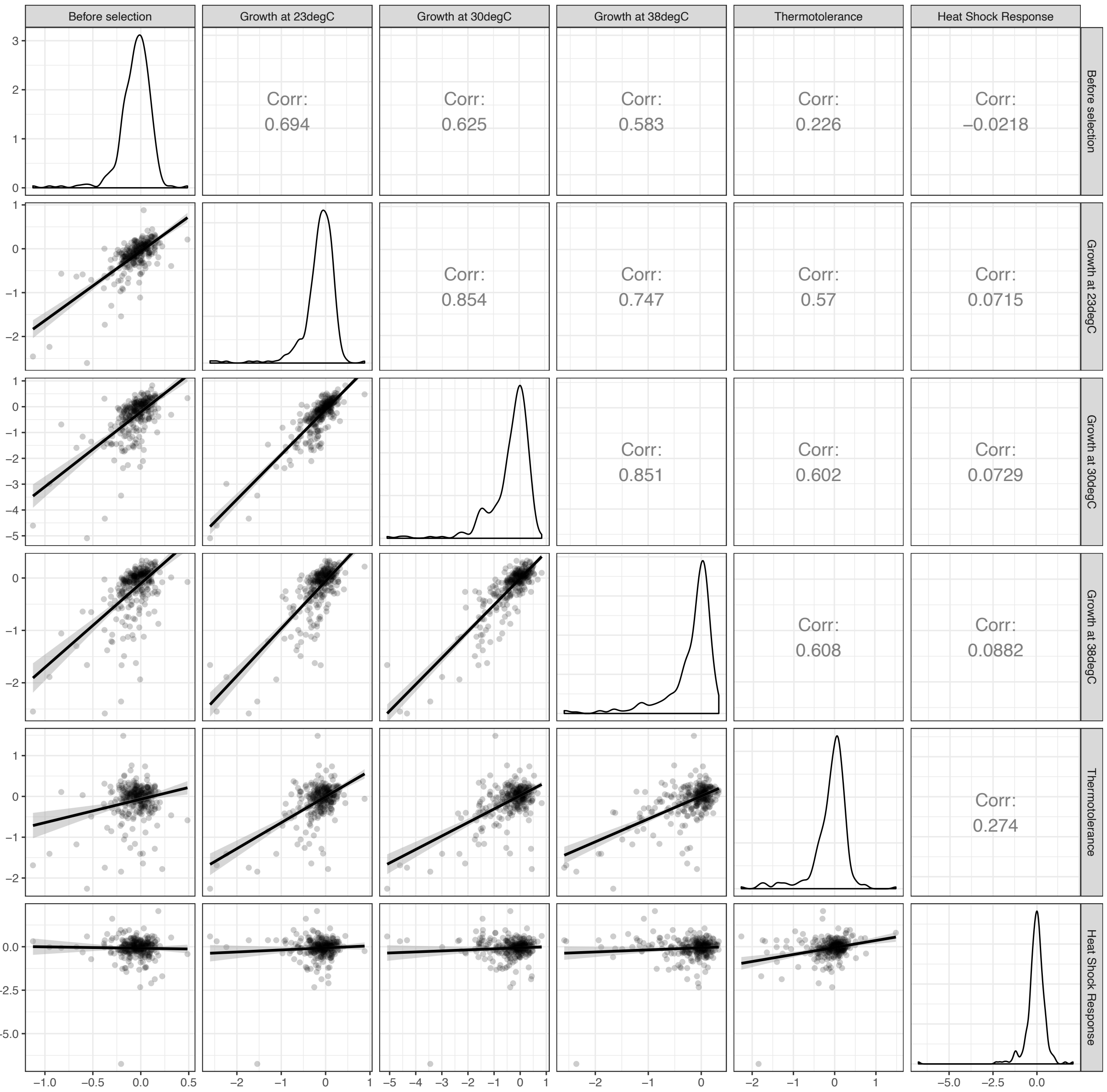


**Suppl. Fig. S33. Gene log2FCs across screens.**
Gene log2FCs (mean of gRNA log2FCs per gene) are compared between screens as indicated in the panel strips, merged for TF and PK libraries (n=290). Dots denote genes. The graph was generated with the ggpairs function of the ggally R package (Schloerke et al., 2020) and displays Pearson correlations. Linear model for smoothed line is generated in R ggplot using geom_smooth with method="lm" and default settings. Please note that fitness screens are not normalized for generation number during selection.

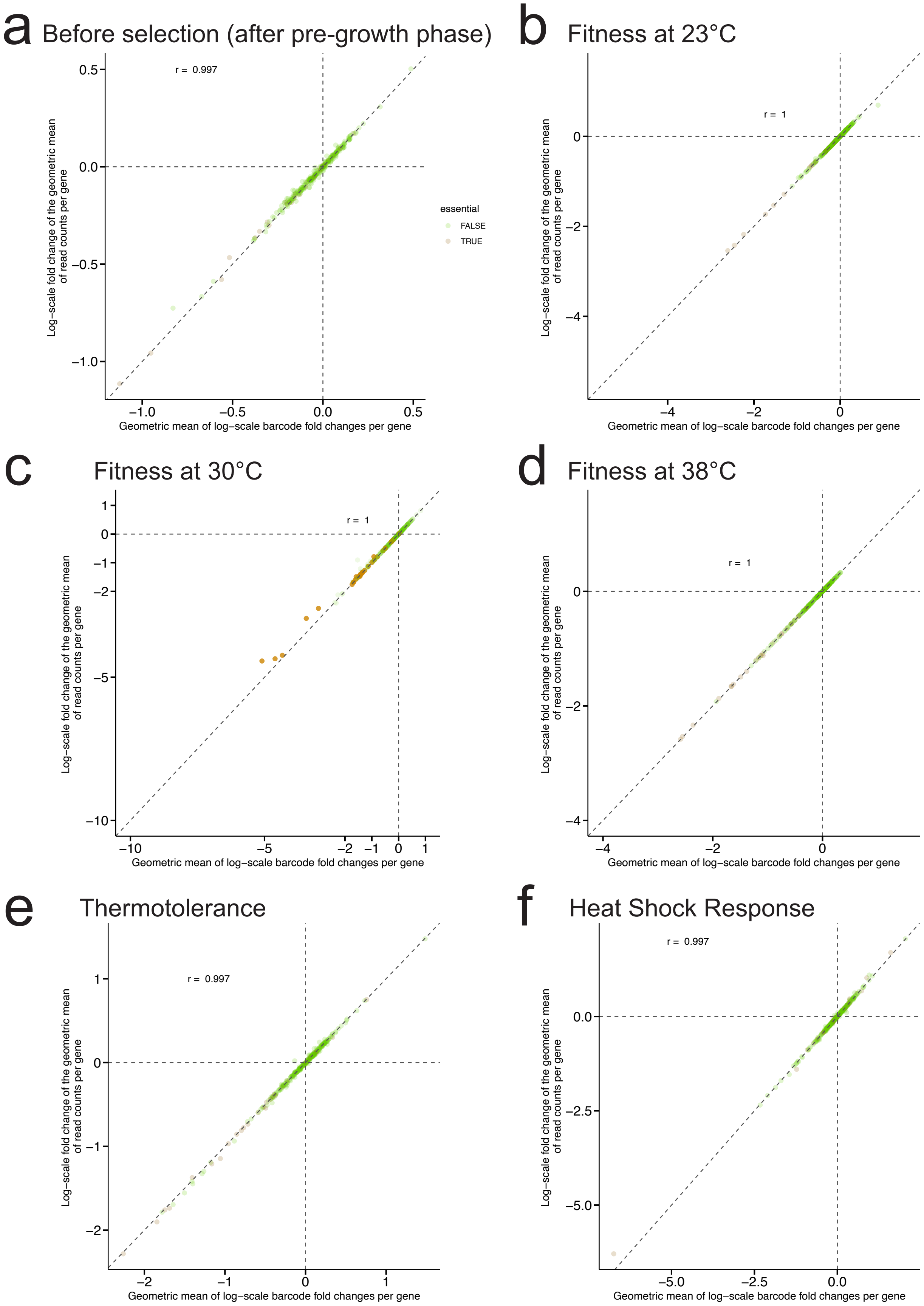


**Suppl. Fig. S34. Correlation of gene scores computed from read counts or gRNA effects.**
Our analysis pipeline uses DNA barcode read-counts as input and computes both, gRNA and gene fold changes along with adjusted p-values based on analysis with edgeR (Robsinson et al., 2009) or other read count-based methods. The gene log2FC can be computed as the average of pre-calculated gRNA log2FCs (x-axis), or directly from raw read counts (y-axis) when using the geometric mean of read counts as input .Comparison of these gene scores is shown for all screens, as indicated in titles for n=290 genes.

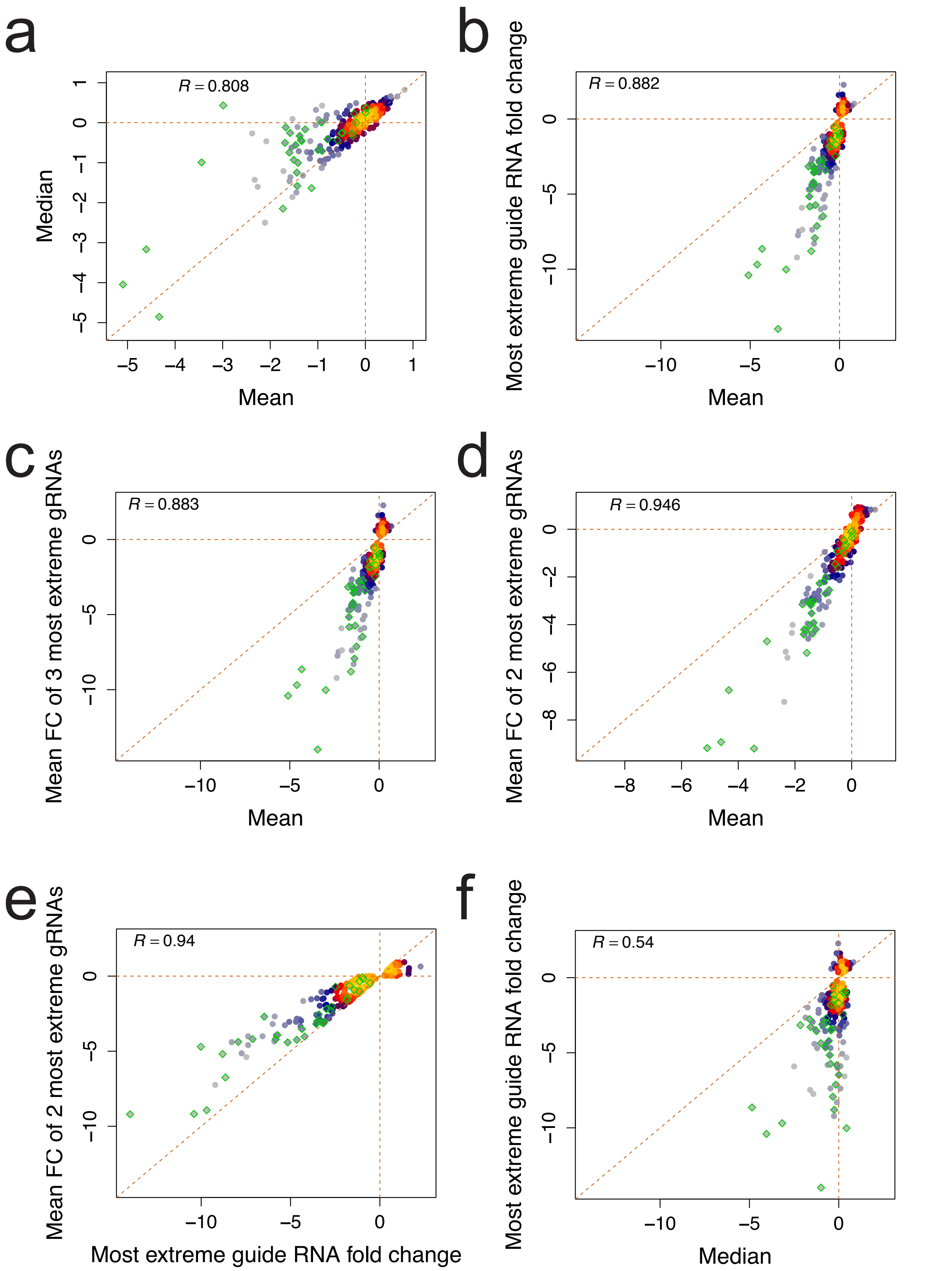


**Suppl. Fig. S35. Comparison of gene score measures and their capture of essential genes.** Different approaches for computation of gene scores from individual gRNA log2FCs are compared in scatter plots based on data of the 30°C fitness screen, including the mean (a-d) and median of all gRNAs (a and f), the mean of the 3 (c) and 2 most extreme gRNAs (d and e), and only the most extreme gRNA log2FC (b and e). Dots denote genes (n=290) and are colored based on density with the LSD R package (Schwalb et al., 2018). Essential genes are marked as green diamonds. Dashed orange lines denote intercepts at 0 and the diagonal. Spearman correlations are shown.

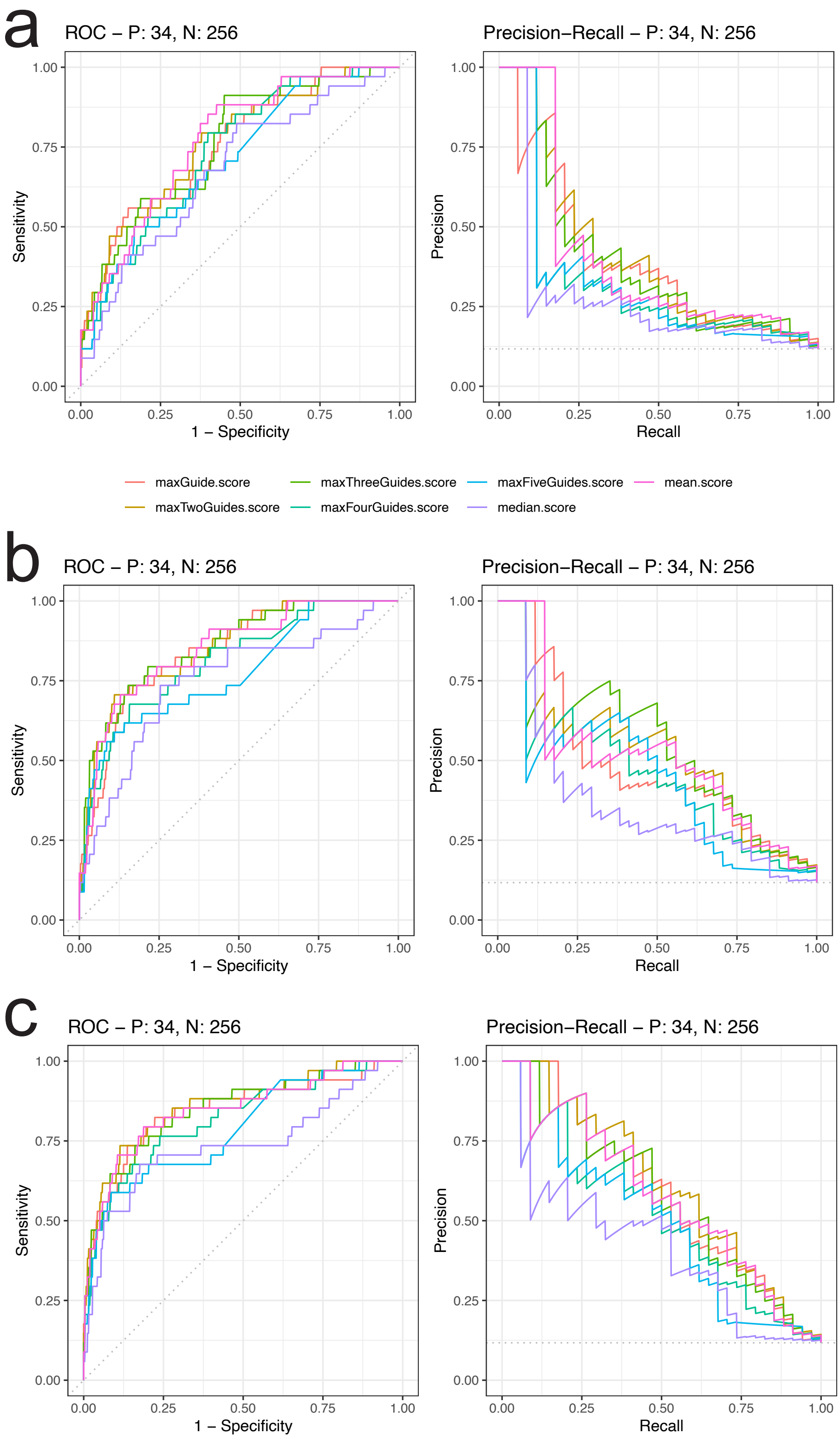


| Fitness at 23°C | | |
|---|---|---|
| Score | AUC ROC | AUC PRC |
| maxGuide.score | 0,75 | 0,39 |
| maxTwoGuides.score | 0,76 | 0,41 |
| maxThreeGuides.score | 0,76 | 0,39 |
| maxFourGuides.score | 0,74 | 0,33 |
| maxFiveGuides.score | 0,71 | 0,32 |
| median.score | 0,68 | 0,27 |
| mean.score | 0,77 | 0,39 |

| Fitness at 30°C | | |
|---|---|---|
| Score | AUC ROC | AUC PRC |
| maxGuide.score | 0,84 | 0,48 |
| maxTwoGuides.score | 0,85 | 0,52 |
| maxThreeGuides.score | 0,86 | 0,53 |
| maxFourGuides.score | 0,81 | 0,45 |
| maxFiveGuides.score | 0,77 | 0,44 |
| median.score | 0,75 | 0,37 |
| mean.score | 0,85 | 0,50 |

| Fitness at 38°C | | |
|---|---|---|
| Score | AUC ROC | AUC PRC |
| maxGuide.score | 0,84 | 0,58 |
| maxTwoGuides.score | 0,86 | 0,61 |
| maxThreeGuides.score | 0,85 | 0,58 |
| maxFourGuides.score | 0,82 | 0,51 |
| maxFiveGuides.score | 0,79 | 0,48 |
| median.score | 0,74 | 0,41 |
| mean.score | 0,85 | 0,57 |

**Suppl. Fig. S36 ROC and PRC curves of gene score measures.**
The Receiver Operatic Characteristic (ROC) and Precision Recall Curve (PRC) is shown for gene score measures indicated in the legend. Their integrals (area under the curve: AUC) are listed. Plots are based on data of the a) 23°C, b) 30°C and c) 38°C fitness screens. Essential genes are used as positive controls (n=34), and non-essentials as negative controls (n=256).
*Please note that positive and negative controls are not perfect. Positive controls are defined by knock-out and may not cause fitness defects upon repression if a low amount of residual mRNA is sufficient for viability. Likewise, many negative control genes will have fitness defects also without being defined as essential.*

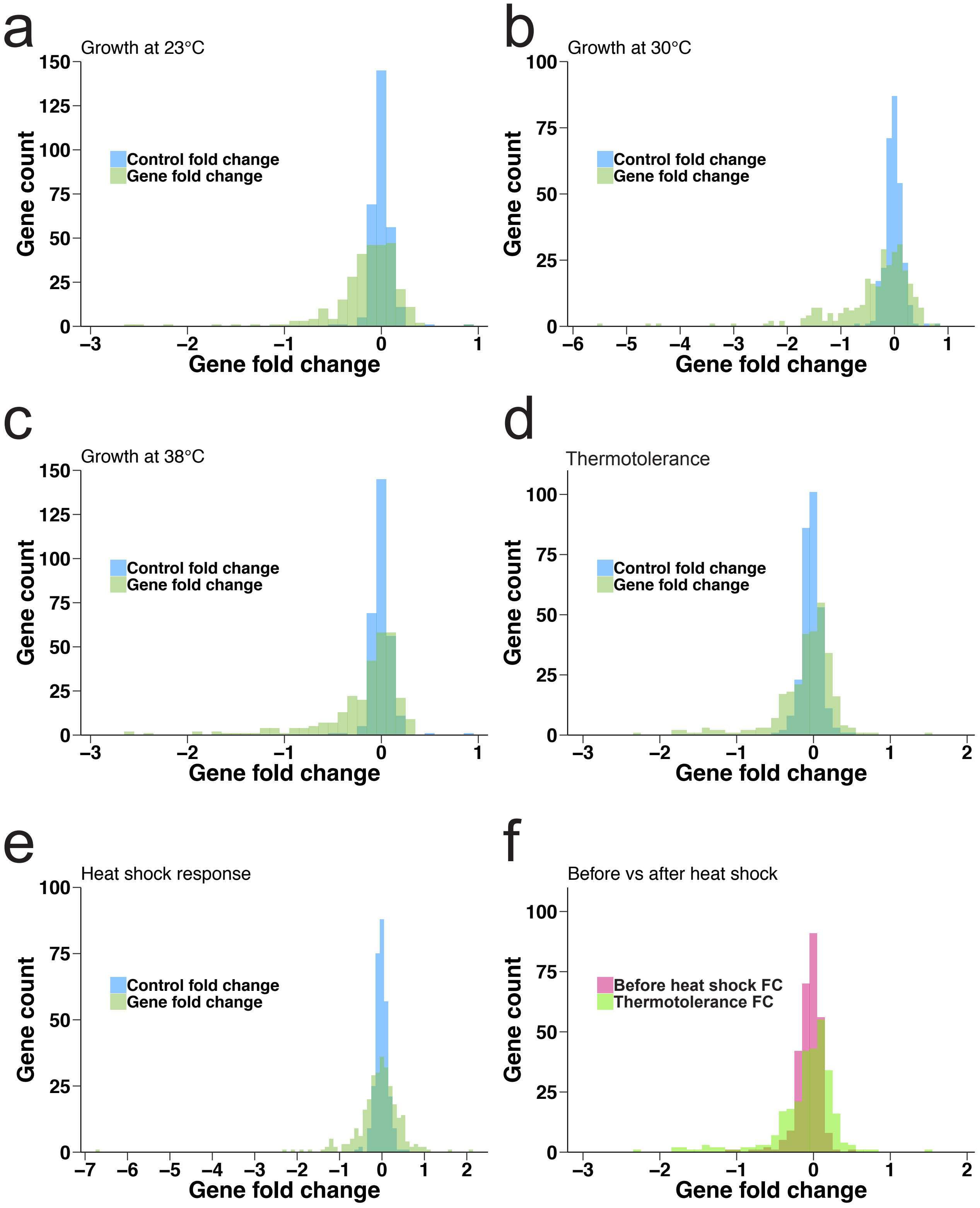


**Suppl. Fig. S37. Control gene scores to estimate technical variation.**
Comparison of CRISPRi gene fold changes with control scores for each screen, as indicated in the title: Fitness screens at a) 23, b) 30, c) 38°C, d) thermotolerance screen and e) HSR screen. Gene scores represent mean gRNA log2FC per gene (n=290). Control scores are fold changes between -ATc samples and represent background variation without CRISPRi induction. f) CRISPRi gene fold changes before and after heat shock from the thermotolerance screen.
